# Systematic revision of the family Tetillidae (Porifera: Demospongiae) in the temperate Northeast Pacific

**DOI:** 10.1101/2025.09.30.679634

**Authors:** Thomas L. Turner

## Abstract

This study presents the first comprehensive systematic revision of the sponge family Tetillidae in the temperate Northeast Pacific. The findings reveal a previously unrecognized clade of *Tetilla* species that informs the family’s generic classification, including a common California species whose identity has long been uncertain. Remarkably, eight species were discovered from only 11 samples from the Aleutian Islands of Alaska, suggesting these islands harbor considerable undiscovered sponge diversity. I also report the first likely case of species introduction in the family: the Asian species *Tetilla japonica* introduced to Southern California. In total, this work refines our understanding of nine previously described species and formally describes eight new species as ***Craniella amlia* sp. nov., *Craniella shemya* sp. nov., *Craniella columbiana* sp. nov., *Craniella rocheta* sp. nov., *Craniella uniiguni* sp. nov., *Craniella vermisigma* sp. nov., *Tetilla losangelensis* sp. nov.,** and ***Tetilla vancouverensis* sp. nov.** Multi-locus DNA sequence data are presented for 13 species, including several low-coverage mitochondrial genomes and complete ribosomal sequences assembled de novo from Illumina sequencing. Together, these results significantly advance our understanding of global tetillid diversity, systematics, and the biogeography of sponge diversity in the Northeast Pacific.

## Introduction

The family Tetillidae Sollas, 1886 comprises 10 genera of spherical or subspherical sponges distributed worldwide, from the deep ocean to the intertidal zone (Van Soest & Rützler, 2002; de Voogd, *et al*., 2025). Two of these genera are known from the temperate Northeast Pacific: *Tetilla* Schmidt, 1868 and *Craniella* Schmidt, 1870. Distinguishing between these two genera has long been difficult and controversial. Shifting definitions have resulted in species being repeatedly reassigned (e.g. Lehnert & Stone 2011). However, recent studies integrating DNA and morphological data have brought greater clarity (Szitenberg *et al*., 2013; Carella *et al*., 2016; Carella & Uriz, 2018). The most recent diagnosis of *Craniella* states that these are globular sponges, lacking the porocalices found in some tetillid genera, and possessing a prominent cortex. This cortex is visible to the naked eye, with a collagenous inner layer reinforced with cortical oxeas, overlain by a thinner outer layer often separated by subdermal cavities (Carella *et al*., 2016). *Tetilla* are said to also lack porocalices, but in contrast to *Craniella*, they lack cortical specialization and lack cortical oxeas (Carella *et al*., 2016). These diagnoses differ from older ones mainly in delineating what constitutes a specialized cortical layer (Van Soest & Rützler, 2002). Intermediates are also found: some species exhibit a “pseudocortex”, which lacks the double-layered structure of *Craniella* but has cortical oxeas. Sponges with this arrangement were found to be distantly related to *Tetilla* and *Craniella*, and have been placed to their own genus, the *Antarctotetilla* Carella et al., 2016.

Seven species of *Tetilla* and *Craniella* were previously known in the temperate Northeast Pacific (these are also the only local representatives of the entire suborder Spirophorina). Two species are described from California, *Tetilla mutabilis* de Laubenfels 1930 and *Tetilla arb* de Laubenfels 1930. *Tetilla mutabilis* is a clavate species that was described from shallow, soft sediments in a single California bay (de Laubenfels, 1932). Its morphology is similar to the type species *T. euplocamos* Schmidt, 1868. Both species are poorly known, but they are united by the absence of sigmaspires—spicules that typically define the suborder but are sometimes missing.

In contrast, *T. arb* was described as a roughly spherical sponge with a cortex, and was therefore moved to the genus *Craniella* by subsequent authors. This species is also very poorly known, with confusion stemming from several issues: poor preservation of the type specimen, an apparent typo in the original description, and the presence of different spicules depending on life stage (see results below). A Master’s thesis previously attempted to resolve these issues, but the effort was ultimately unsuccessful (Ristau, 1977). Although other sponges described in the thesis were later published in a peer-reviewed paper, the tetillids were excluded and never formally described (Ristau, 1978). I interpret this omission to mean that the author or reviewers judged the data insufficient for valid taxonomic publication. This also makes it clear that the thesis was not intended as the formal publication of new names under the rules of zoological nomenclature.

Several *Craniella* species are also known from farther north in the Pacific. Two species are described from British Columbia: *C. spinosa* Lambe 1893 and *C. villosa* Lambe 1893. Their type specimens have not been reexamined since the original descriptions, and a modern analysis of their status and distribution is needed. More recently, *C. sputnika* Lehnert & Stone, 2011 was described from the Aleutian Islands, Alaska. The same year, *C. sigmoancoratum* (Koltun, 1966) was also reported from the Aleutians (Stone *et al*., 2011).

We therefore begin with six species of Tetillidae known from the temperate Northeast Pacific—most described more than a century ago and with little subsequent study. Later surveys of intertidal Tetillidae in California failed to relocate the two originally described species and instead identified two distinct morphotypes, provisionally referred to as *Tetilla* sp. A and *Tetilla* sp. B (Hartman, 1975; Lee *et al*., 2007). These gaps underscore the need for a comprehensive regional revision, which is undertaken here. For this study, loans were obtained from six museum collections, including the holotypes of *T. mutabilis, T. arb, C. villosa,* and *C. spinosa* (as well as the holotype of the Arctic species *C. craniana* de Laubenfels 1953, for comparison). In addition, 30 new samples were collected from across Southern and Central California, Washington, and British Columbia. Alongside detailed morphological descriptions, DNA sequence data from these samples are integrated with publicly available datasets to produce an updated global phylogeny of the family.

## Materials & Methods

### Collections

SCUBA-based collections were conducted by the author at 128 sites spanning San Diego, California to Juneau, Alaska (2019–2025). Search effort was concentrated in Southern and Central California, with additional sampling at five sites in Washington, four in British Columbia, and five in Alaska. Intertidal collections were made at 15 additional sites across the same range. Floating docks in 23 harbors and marinas were surveyed, but no Tetillidae were found at these sites. Brandon Stidum, Siena McKim, and Jeff Goddard each collected and contributed an additional sample.

Archived material was obtained on loan from the Smithsonian National Museum of Natural History (USNM), the Royal British Columbia Museum (RBC), the California Academy of Sciences (CASIZ), the Santa Barbara Museum of Natural History (SBMNH), the Natural History Museum of Los Angeles (NHMLA), and the Canadian Museum of Nature (CMNI). Newly collected material was vouchered at these institutions as well as the Cheadle Center at the University of California, Santa Barbara (UCSB), the Florida Museum of Natural History (BULA), and the Museum of Evolution in Uppsala (UPSZMC). Voucher numbers are listed in the Systematics section, tabulated in Table 1, and provided in an expanded table S1 in the supporting information. The expanded table includes expanded metadata in a convenient format, including field numbers, museum vouchers, GenBank accession numbers, collection locations and depths, iNaturalist links, and other notes.

**Table 1.** Vouchering information for samples and DNA sequences. See supporting information for an expanded version including metadata for each sample.

| Species | Type status | Museum vouchers | Mitochondrial Voucher(s) | Ribosomal voucher(s) |
| --- | --- | --- | --- | --- |
| *Craniella amlia* | Holotype | USNM 1478667 | PX230100 | n/a |
| *Craniella shemya* | Holotype | USNM 1478666 | PX230103 | PX227227 |
| *Craniella columbiana* | Paratype | RBC A-094-00006 | n/a | n/a |
| *Craniella columbiana* | Holotype | RBC 980-00259-006 | PX230104 | PX227225 |
| *Craniella craniana* | Holotype | USNM 23233 | n/a | n/a |
| *Craniella hamatum* | n/a | USNM 1478661 | PX230105 | PX227224 |
| *Craniella columbiana* | Paratype | RBC 975-00072-024 | PX238412 | n/a |
| *Craniella rocheta* | Holotype | USNM 1517729 | PX230106 | PX227208 |
| *Craniella sigmoanchoratum* | n/a | USNM 1478663 | PX230101 | PX227230 |
| *Craniella spinosa* | Holotype | CMNI 1900-2814 | n/a | n/a |
| *Craniella sputnika* | n/a | USNM 1478668 | PX230107 | PX227226 |
| *Craniella sputnika* | n/a | USNM 1478669 | PX230116 | PX227233 |
| *Craniella uniiguni* | Paratype | USNM 1478664 | PX230102 | PX227229 |
| *Craniella uniiguni* | Holotype | USNM 1478662 | PX230108 | PX227228 |
| *Craniella vermisigma* | Holotype | USNM 1478660 | PX230109 | PX227231 |
| *Craniella vermisigma* | Paratype | USNM 1478700 | PX230110 | PX227232 |
| *Tetilla arb* | Holotype | USNM 21496 | n/a | n/a |
| *Tetilla arb* | n/a | TBD before publication | n/a | n/a |
| *Tetilla arb* | n/a | CASIZ 302844 | PX230090 | PX227210 |
| *Tetilla arb* | n/a | TBD before publication | PX230086 | PX220134 |
| *Tetilla arb* | n/a | BULA-0164. NHMLA 15491 | n/a | PX227209 |
| *Tetilla arb* | n/a | NHMLA 16558 | PX230087, PX232379 | PX220135 |
| *Tetilla arb* | n/a | TBD before publication | PX230089 | PX227202 |
| Tetilla arb | n/a | TBD before publication | n/a | PX227200 |
| Tetilla arb | n/a | TBD before publication | n/a | PX227203 |
| Tetilla arb | n/a | TBD before publication | n/a | PX227204 |
| Tetilla arb | n/a | TBD before publication | n/a | n/a |
| Tetilla arb | n/a | TBD before publication | n/a | n/a |
| Tetilla arb | n/a | TBD before publication | n/a | n/a |
| Tetilla arb | n/a | TBD before publication | n/a | n/a |
| Tetilla arb | n/a | TBD before publication | n/a | n/a |
| Tetilla arb | n/a | TBD before publication | PX230098 | PX227214 |
| Tetilla arb | n/a | TBD before publication | n/a | PX227215 |
| Tetilla arb | n/a | TBD before publication | PX230083 | PX227216 |
| Tetilla arb | n/a | TBD before publication | PX230096 | PX227211 |
| Tetilla arb | n/a | TBD before publication | PX230085 | PX227212 |
| Tetilla arb | n/a | TBD before publication | PX230097 | PX227213 |
| Tetilla arb | n/a | TBD before publication | PX230094 | PX227201 |
| Tetilla japonica | n/a | SBMNH 47765 | n/a | n/a |
| Tetilla japonica | n/a | SBMNH 659949 | n/a | n/a |
| Tetilla japonica | n/a | SBMNH 659950 | n/a | n/a |
| Tetilla japonica | n/a | TBD before publication | PX230115 | PX227205 |
| Tetilla losangelensis | Holotype | NHMLA 15495, BULA-0180 | PX230111 | PX227217 |
| Tetilla mutabilis | Holotype | USNM 21498 | n/a | n/a |
| Tetilla vancouverensis | n/a | RBC 981-00216-014 | PX238413 | n/a |
| Tetilla vancouverensis | n/a | RBC 018-00212-001 | PX230091 | PX227206 |
| Tetilla vancouverensis | n/a | TBD before publication | PX230092 | PX227218 |
| Tetilla vancouverensis | Paratype | TBD before publication | n/a | n/a |
| Tetilla vancouverensis | Paratype | TBD before publication | PX230112 | PX227219 |
| Tetilla vancouverensis | Paratype | TBD before publication | PX230113 | PX227220 |
| Tetilla vancouverensis | Holotype | TBD before publication | PX230114 | PX227221 |
| Tetilla villosa | Holotype | CMNI 1900-2815 | PX238414 | n/a |
| Tetilla villosa | n/a | RBC 981-00216-015 | PX238415 | n/a |
| Tetilla villosa | n/a | USNM 21497 | n/a | n/a |
| Tetilla villosa | n/a | RBC 980-00335-008 | PX238416 | n/a |
| Tetilla villosa | n/a | RBC 007-00011-002 | PX230088 | n/a |
| Tetilla villosa | n/a | CASIZ 235692 | n/a | n/a |
| Tetilla villosa | n/a | TBD before publication | PX238417 | n/a |
| Tetilla villosa | n/a | TBD before publication | PX230095 | PX227207 |
| Tetilla villosa | n/a | TBD before publication | PX230099 | PX227223 |
| Tetilla villosa | n/a | TBD before publication | PX230084,<br>PX232378 | PX220136 |
| Tetilla villosa | n/a | TBD before publication | PX230093 | PX227222 |

### Morphology

Skeletal architecture was examined by hand-cutting tissue sections and digesting them in 97% Nuclei Lysis Solution (Promega, Wizard Genomic DNA Purification Kit) and 3% Proteinase K (20 mg/ml, Promega). Spicules were isolated by digesting sponge subsamples in bleach. Light micrographs were taken with a Nikon D3500 SLR and an Amscope NDPL-1 microscope adaptor; scanning electron micrographs (SEMs) were taken with a FEI Quanta400F Mk2 after coating spicules with 20 nm of carbon. Measurements were made in ImageJ (Schneider *et al*., 2012). For simple spicules (e.g., oxeas), length was measured as the longest possible straight line from tip to tip, even when curved; some very long spicules are an exception, as they were measured in several straight intervals across multiple photos using landmarks. Spicule width was measured at the widest point. Spicules like anatriaenes and protriaenes have a complex morphology, and additional measurements were made on each. Anatriaenes: rhabd length (excluding clads), rhabd width (widest point near the clad end), clad length (tip of clad to top of spicule), clad width (measured where the clad meets the rhabd), and clad angle (the angle from rclad tip to the top of the spicule to center of rhabd). Protriaenes: rhabd length and width as above, longest and shortest clad lengths (tip of clad to rhabd junction), clad width (widest point), and clad angle (180° minus the angle from rhabd center to clad junction to clad tip). Figures S1-S7 provide a visual guide to these measurements in the supporting information. Several measurements are subject to considerable error (e.g., the central location where clads gather with the rhabd can only be approximated, and the measurements of angles will vary based on the orientation of the spicule in the image), but error was minimized by recording maximum values and averaging across many spicules. Despite many potential sources of variability, measurements consistently distinguished species and were useful for diagnoses.

For statistical comparisons among *Craniella* species, ANOVAs were performed using the *aov* function in R, with pairwise differences tested using Tukey’s HSD (*emmeans* and *TukeyHSD*). For *Tetilla*, t-tests (*t.test*) or Wilcoxon rank-sum tests (*wilcox.test*) were used. Measurements are reported as minimum–mean–maximum with *n* equal to the number of spicules measured (which may differ across traits due to spicules with broken portions). All 21,039 spicule measurements made are provided in the supporting table S2.

### Illumina sequencing

DNA was extracted using the Qiagen Blood & Tissue kit, treated with RNase A, and repurified with the Zymo DNA Clean & Concentrator. Library preparation and sequencing were performed at the UC Davis DNA Technologies Core Facility using the Super-High-Throughput (SHT) protocol, with dual indexing of 96 pooled samples. Three tetillid sponges were included, alongside other sponges analyzed elsewhere (Turner *et al*., 2025). Libraries were sequenced on an Illumina NovaSeq, generating paired-end 150 bp reads with a median of 2.1 million reads per sample (range: 208–17 million). This variability likely reflected differences in DNA preservation and secondary chemistry.

Reads were trimmed with fastP (Chen *et al*., 2018) and assembled de novo with Megahit (Li *et al*., 2015). Assemblies yielded 0–1,080 contigs per sample (median: 22). Blast searches indicated that many contigs were microbial, or occasionally from fish, mollusks, or other taxa, but the nuclear ribosomal locus was typically the highest-coverage contig. Nearly complete ribosomal loci were assembled for all three tetillid sponges. These contigs were then used as references: reads were aligned with bwa-mem (Li & Durbin, 2009), and consensus sequences were generated with SAMtools (Danecek *et al*., 2021). This served as the final sequence for that sample. Mitochondrial contigs were present in some of the Megahit assemblies but were less complete than ribosomal loci. To generate more complete mitochondrial genomes, reads were aligned to the reference mitochondrial genome of *Cinachyrella kuekenthali* (NC_010198) using bwa-mem, followed by the same consensus-building approach.

### Sanger sequencing

I attempted to sequence the Folmer region of cox1 and the D1D2 and D3D5 regions of 28S for all species. However, commonly used poriferan primers performed poorly in tetillids: Folmer primers worked only in *T. japonica*, and the D3D5 region was the only 28S locus to amplify consistently. Using Illumina assemblies and GenBank data, I designed new primer sets. Amplicons 700–850 bp in length worked well in recent collections but often failed in older samples, so additional primers targeting 100-300 bp regions were developed. In some cases, separate primers were designed for *Tetilla* and *Craniella*. Primer sequences are provided in the supporting information. Sequences have been deposited in GenBank, except for fragments rejected due to short length; these are available on request. Accession numbers are listed in Table 1 and in the supporting table S1.

### Phylogenetic methods

Tetillid sequences covering the Folmer region of cox1 and the C2–D2 region of 28S were compiled via BLAST searches, along with relevant outgroups. These data include unpublished datasets and numerous previous studies (Lavrov *et al*., 2008; Szitenberg *et al*., 2010, 2013; Cárdenas *et al*., 2011; Carella *et al*., 2016; Erpenbeck *et al*., 2016; Schuster *et al*., 2017, 2018). Alignments were generated in MAFFT v.7 (Katoh *et al*., 2017). Phylogenies were inferred with maximum likelihood in IQ-TREE (Nguyen *et al*., 2015; Trifinopoulos *et al*., 2016), with nodal support assessed using ultrafast bootstrapping and SH-aLRT tests (Hoang *et al*., 2018). Optimal substitution models were selected with ModelFinder (Kalyaanamoorthy *et al*., 2017); TN+F+I+G4 was chosen for both loci. A combined dataset was analyzed by concatenating loci in a partitioned analysis. Final trees were visualized with the Interactive Tree of Life webserver (Letunic & Bork, 2019).

## Results

### Molecular phylogenies

Figure 1 shows the concatenated, partitioned phylogeny of Tetillidae based on 28S and cox1 loci. Separate trees for each locus (which include additional taxa) are provided in the supporting information. All temperate Northeast Pacific species are represented in Figure 1, with three exceptions. One species described here (*C. amlia* sp. nov.) is represented only in the cox1 tree, as 28S data could not be obtained. Two previously described species (*T. mutabilis* and *C. spinosa*) lacked DNA amplification entirely but are treated in the systematics section.

**Figure 1.**
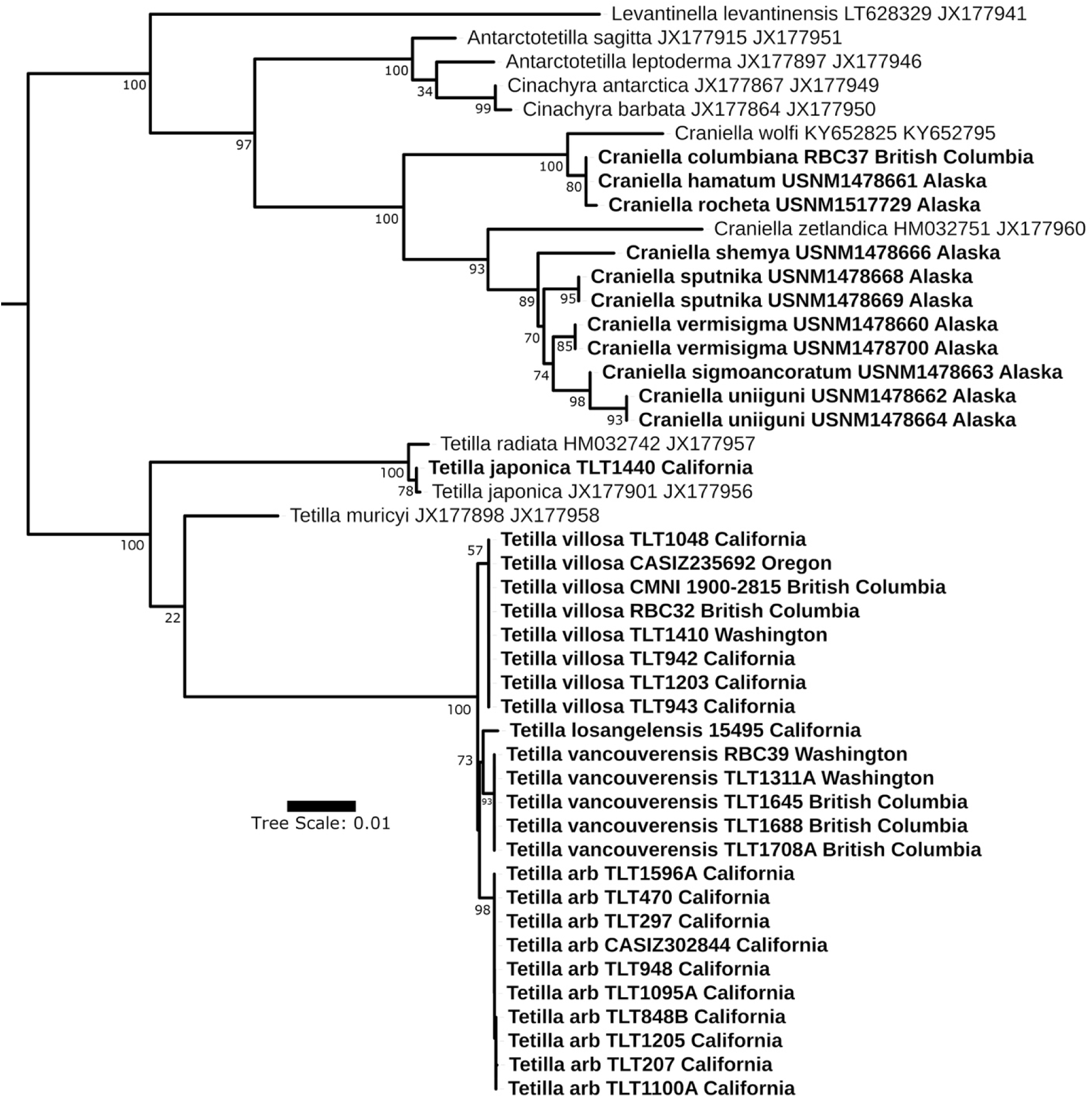
(below). Maximum likelihood phylogeny of the Tetillidae. All species with data at both cox1 and 28S are shown. New sequences are shown in bold, along with collection location and voucher number. Data from Genbank includes accession numbers and collection locations (when known). In some cases, cox1 and 28S data from Genbank are combined from different samples of the same species. Node confidence is indicated with bootstrap values. Scale bar indicates substitutions per site.

The type species of *Tetilla*, *T. euplocamos* Schmidt, 1868, lacks DNA data. Based on morphology, it likely belongs in the clade containing *T. japonica* Lampe, 1886 and *T. radiata* Selenka, 1879. *T. mutabilis* likely belongs here as well, but the holotype was the only sample I could locate, and DNA amplification from it was unsuccessful. Another representative of this clade was found in San Diego Bay (sample TLT1440; Figure 1). Morphologically similar sponges have been known in California bays since the 1970s, but their identity was previously unresolved (Hartman, 1975; Lee *et al*., 2007). Unexpectedly, this sample is nearly identical to sequences from a Thailand sample at both loci: they differ by one base pair at cox1 and are identical at 28S (Szitenberg *et al*., 2013). The Thai specimen was identified as *T. japonica* by Sumait Putchakarn, TLT1440 matches descriptions of *T. japonica* from Japan (Lebwohl, 1914), and does not match any other described species (see systematics section). I therefore assign the California sample to *T. japonica*, though I acknowledge that this is somewhat tentative, as I have not examined the Thai sample or the type material for *T. japonica*. If correct, this would be the first likely case of human-mediated range expansion known in Tetillidae, joining a growing list of introduced sponges (Turner, 2020; Cavalcanti *et al*., 2020; Bettcher *et al*., 2024). California bays have some of the highest proportions of non-native species known (Ruiz *et al*., 2013), making introduction from Asia a likely possibility. Introduction from California into Asia cannot be excluded, but is less likely, as this species was described from Japan in 1886 (Lampe, 1886). It is also possible that future genetic and/or morphological analyses of Japanese *Tetilla* will reveal that the Thai and California samples belong to an undescribed species similar to *T. japonica,* but in that case, it remains likely that the current distribution of this species is a result of human-mediated introduction.

Globular sponges like *T. japonica* have been consistently placed in *Tetilla* for many years, but defining the differences between *Tetilla* and *Craniella* for spherical sponges has been fraught. *Craniella villosa*, described from British Columbia, has remained in *Craniella* since its description, whereas *Tetilla arb* was originally described in *Tetilla* but later moved to *Craniella* (de Laubenfels, 1932; de Voogd, *et al*., 2025). Morphologically, both lack the prominent double-layered cortex of *Craniella*, but they do possess cortical oxeas, said to be absent in *Tetilla* (Carella *et al*., 2016). Though DNA data is not available from the type species of either genus, the phylogenies place both these species in the clade with globular *Tetilla*, while placing all species with a double-layered cortex in a distant clade with other *Craniella*. This is true at both loci individually (figures S8 & S9) and in the concatenated tree (figure 1), so I am confident that these species do not belong in *Craniella*. I therefore propose a slight modification to the diagnosis of the genus *Tetilla* in the systematic section so that these species can be accommodated in this genus. This change makes it harder to differentiate *Tetilla* and the recently erected *Antarctotetilla* Carella, Agell, Cárdenas & Uriz, 2016, which are said to possess a “pseudocortex” that is similar to what is seen in this clade of North Pacific *Tetilla*. It is likely that a more robust set of diagnoses for the genera within Tetillidae will require additional characters, perhaps in the form of DNA characters, but this higher-level taxonomic revision is beyond the scope of the current paper.

### Species delimitation within *Tetilla*

I attempted Illumina sequencing of three California tetillids with several goals in mind. First, I wished to assemble reference sequences of the mitochondrial genome and nuclear ribosomal locus to aid in both phylogenomic comparisons and primer design for Sanger sequencing. This goal was successful, with complete (6,085 bp) haplotypes of the ribosomal locus from *T. arb* and *T. villosa*, and another *T. arb* sample nearly (99%) complete. Mitochondrial genomes were also assembled, but were less complete, with a 17,948 bp contig for *T. arb* with 9% uncalled bases and a 18,050 bp contig for *T. villosa* with 11% missing data. The only previous mitochondrial genome for the family Tetillidae was from *Cinachyrella kuekenthali* (Lavrov *et al*., 2008), so these sequences greatly increase the data available for the family.

My second goal was to increase the genomic scope of data available to investigate species boundaries among California tetillids by sequencing representative samples from three distinct but similar morphotypes: a small yellow *Tetilla* with a single osculum from the intertidal zone (figure 6A), a larger black *Tetilla* with a single osculum from the subtidal (similar to figure 6B, but with a single osculum), and a large gray *Tetilla* without prominent oscula from the subtidal (figure 7A). Coverage was insufficient to assemble the mitochondrial genome from the intertidal sample, but the ribosomal locus of this sample was nearly identical to the subtidal sponge with a single osculum. Across the 6805 bp ribosomal haplotype, these two sponges differ at only 4 bp (0.06%; samples TLT207 and TLT297 in figure 1). Combined with a lack of differences at the Sanger-sequenced cox1 locus, it seems likely that these two morphotypes are part of a single species. As detailed in the systematics section, morphological taxonomy assigns them to the name *Tetilla arb*.

Divergence (D_xy_) between the two *T. arb* samples and the sponge without prominent oscula — which was subsequently determined to be the species *T. villosa* — is also quite modest, but it is an order of magnitude higher (0.6%) at 28S than seen between the two samples of *T. arb*. The mitochondrial genomes of these samples are very similar, however, differing at only 10 of the aligned base pairs (0.04%). Fortuitously, 2 of these single nucleotide polymorphisms (SNPs) are within the cox1 gene, so I designed sequencing primers that would genotype this region in a larger sample of individuals. Genotyping of 10 *T. villosa* and 10 *T. arb* at cox1 found that both SNPs were fixed differences between species. Likewise, sequencing 8 *T. villosa* and 16 *T. arb* at 28S verified 4 fixed differences across all samples. Because the mitochondrial and nuclear genomes freely recombine during meiosis, reciprocal monophyly across these compartments is strong evidence of reproductive isolation when species are sympatric (Jennings, 1917; Rannala & Yang, 2020). These species are sympatric over a large range (figure 2): I collected them both at a single site in Carmel Bay, Central California, and at several sites off Point Loma, in Southern California (600 km and 4° latitude from Carmel Bay).

**Figure 2.**
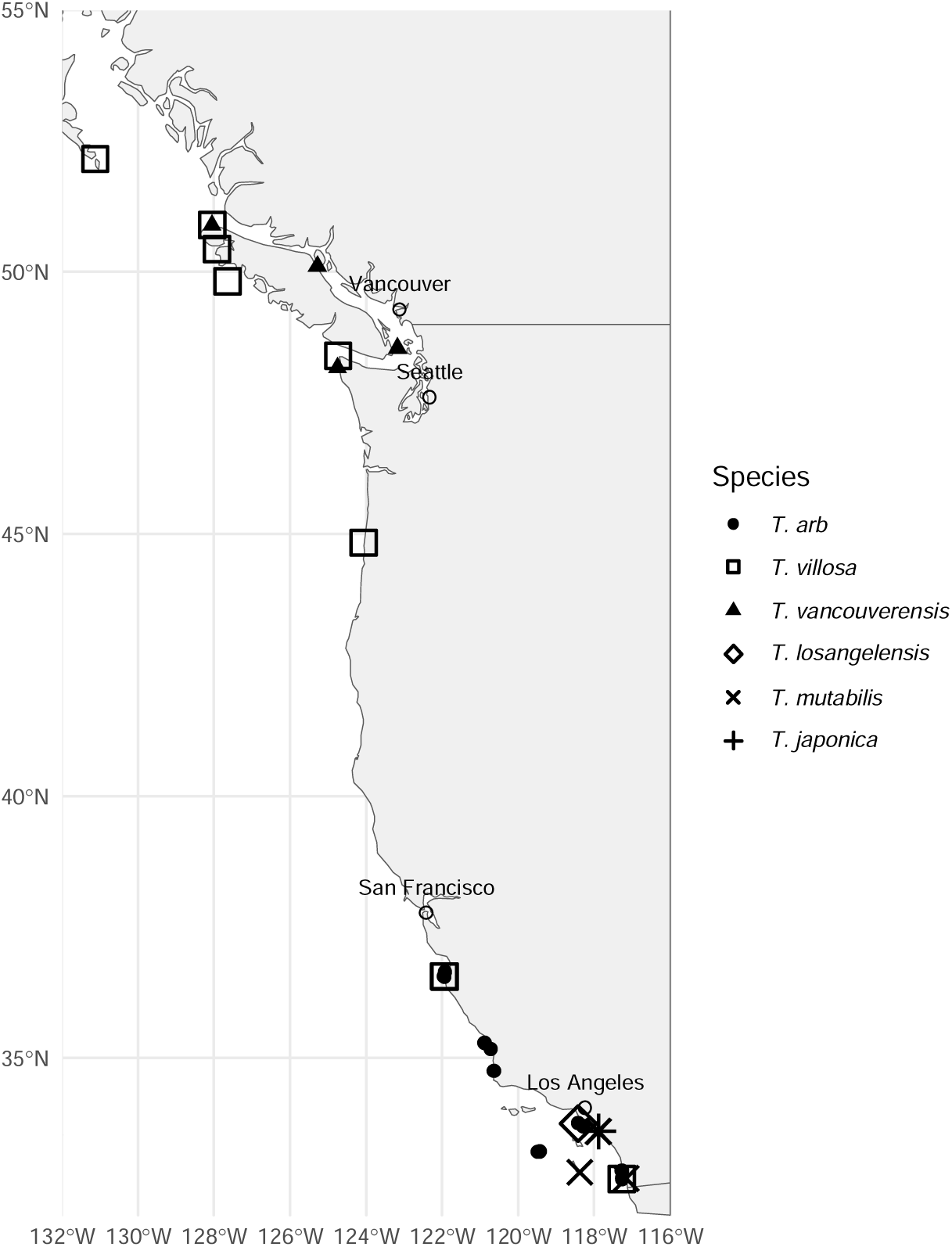
Range map of samples examined for all *Tetilla* species. *T. villosa* was found to have a considerable latitudinal range, while other species were limited to British Columbia (*T. vancouverensis*) or California (all other species). Northern Mexico is ecologically similar to Southern California, and California species are expected to be found there in future work. Photos seen by the author also indicate that *T. arb* likely extends into Northern California, but these samples were not available for verification.

Sequencing of these two loci in all available samples unexpectedly revealed two more reproductively isolated lineages. Six British Columbia sponges initially identified as *T. villosa* showed fixed cox1 and 28S differences relative to both *T. villosa* and *T. arb*. This new species, broadly sympatric with *T. villosa*, is described as *T. vancouverensis* sp. nov. in the systematic section. The other lineage is represented by a single sample, genetically and morphologically distinct from all others. Though naming a species based on a single sample is never ideal, convincing evidence of reproductive isolation among the other three members of the clade make it likely that this samples constituted a fourth species, and it is described below as *T. losangelensis* sp. nov.

### Variation in spicule complement in *Tetilla*

The species concepts inferred from genetic data indicate that some spicule traits varied within species, in correlation with sponge size. The largest population sample is for *T. arb*, with 22 sponges ranging from 7–55 mm in diameter. Additionally, the holotype is estimated to have been the largest known sample, approximately 72 cm in diameter (based on a 36 cm distance from the sponge center to the ectosome in the preserved fragment, and assuming it was spherical). Comparing sponge size to the length of the largest-size class of oxeas (oxeas I) in a subset of these sponges reveals a strong and apparently non-linear relationship (figure 3; Wilcoxon p_=_0.008). The length of oxeas II and oxeas III were also significantly associated with sponge size (Wilcoxon p_<_0.005 in both cases). I only found one small sample of *T. vancouverensis* sp. nov., but comparing this sponge to larger samples reveals a similar pattern (figure 3; Wilcoxon p_<_0.005). This clearly limits our ability to use the dimensions of these spicules as taxonomic traits, at least until enough samples have been characterized to calibrate the relationship between sponge size and spicule length (for sponges 50–60 mm in diameter, *T. vancouverensis* sp. nov. have oxeas 40% larger than *T. arb*, but the small *T. vancouverensis* sp. nov. would be indistinguishable from *T. arb* using the same statistic). Thankfully, other spicule dimensions did not vary with sponge size, such as the sizes and shapes of protriaene and anatriaene clads. I therefore focus on these traits in the systematic section below.

**Figure 3.**
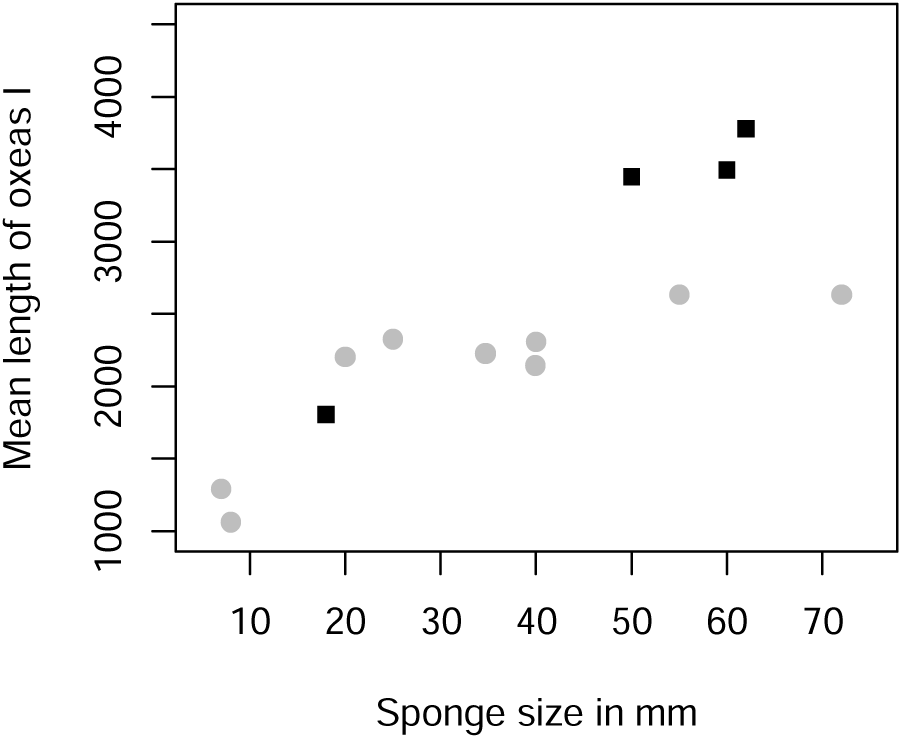
Spicule lengths vary with sponge size. *Tetilla arb* (gray) and *Tetilla vancouverensis* (black) both have a highly significant relationship between the sponge diameter and the mean length of the largest class of oxeas (oxeas I) in that sample.

Sponge size also affected the categorical types of spicules that were present. Anamonaenes (”boathook” spicules, figure 6J) were abundant in small (< 3 cm) *T. arb* individuals, but were rare or absent in large individuals. Conversely, anatriaenes (anchor-shaped spicules, figure 6K) were common in large individuals, but absent or rare in small individuals. The pattern in *T. vancouverensis* sp. nov. was similar. The one small individual had anamonaenes in the basal ectosome, and these were not seen in larger individuals. In contrast to *T. arb*, this species has two types of anatriaenes, and both were present in the small and the large individuals.

To my knowledge, this is the first reported case of a spicule complement that differs among settled sponges of different sizes Some larval sponges are known to have different spicules than adults (Maldonado, 2006). Notably, It has been reported that juvenile *Craniella*, which have direct development within the adult, have different spicules when they are developing within the mother vs. after settling (Burton, 1931). Among the reported differences is a preponderance of anamonaenes rather than anatriaenes, similar to what was found here in *Tetilla*, but this difference was not seen among settled sponges of different sizes.

One hypothesis for this change in spicule type is that anamonaenes facilitate sponge settlement on host sponges. Size is presumably correlated with age, and *T. arb* seems to settle exclusively on other sponges. Small individuals of this species were only observed growing on other sponges; indeed, sampling the substrate under these sponges has proven to be a good way to discover cryptic sponges that resemble inorganic substrate. Larger *T. arb* are often, but not aways, attached to another sponge, perhaps because the tetillid sometimes outlives the death of its host sponge. A similar substrate preference seems likely in related *Tetilla*: 5 of 7 *T. vancouverensis* sp. nov., including the only small sample, were clearly attached other sponges, and 3 of 11 *T. villosa* were attached to other sponges, despite not having any small sponge samples from this species. If young sponges use anamonaenes to anchor themselves to other sponges, this could explain why they produce them mainly when small, and especially in the basal region of the sponge. This explanation is not entirely satisfactory, because larger *T. arb* are found to have anatriaenes embedded deeply into their host sponges, so it is unclear why the sponges would switch the type of embedded anchoring spicules with age. Regardless of the reason, this is an additional observation that indicates that spicule-based taxonomy of the Tetillidae is best done with caution, preferably in combination with DNA data.

### Remarkable *Craniella* diversity in the far North Pacific

The Aleutian Islands are a 1,700 km volcanic island chain stretching across the far north Pacific, separating this region from the Bering Sea. Deep-water habitats in the Aleutian Islands are known to support animal forests and sponge grounds that are among the most diverse and ecologically important deep water habitats in the world (Stone, 2014; Beckmann *et al*., 2025). I examined 11 vouchers of *Craniella* that had been collected in the Aleutians, mostly by the NOAA Alaska Fisheries Science Center. Remarkably, these 11 *Craniella* samples yielded 8 species: one known Aleutian species, two originally described from the nearby Kuril Islands, and five new species. Most were deep-water taxa, with only *C. rocheta* sp. nov. collected on SCUBA. Eight species from only 11 samples is extraordinary, but these species concepts are supported by both genetic data and unique morphological traits, as described in the systematics section. Despite considerable recent progress describing the diversity of sponges in this region (e.g. (Stone *et al*., 2011; Lehnert & Stone, 2015)), the findings reported here suggest that many sponges in the region remain to be characterized.

Two additional *Craniella* were found in British Columbia: the previously described *C. spinosa* and *C. columbiana* sp. nov. With *T. arb* and *T. villosa* reassigned to *Tetilla*, no *Craniella* were found South of 48°N in the region. To the South, the next known *Craniella* is *C. wolfi* Schuster *et al*. 2018, just north of the equator, in the Galapagos Islands (Schuster *et al*., 2018). *Craniella lissi* Sim-Smith *et al*. 2021 is also described from the Galapagos, but appears to be a sediment-rooting *Tetilla* species.

### DNA sequencing of museum vouchers

The 28S and cox1 loci are the most commonly used genes for sponge systematics, because they are conserved enough to amplify and align, but variable enough to be informative (Erpenbeck *et al*., 2006; Thacker *et al*., 2013). Sequencing these loci in tetillids was challenging because “universal” poriferean primers failed to amplify a product in most samples. This is likely due to sequence mismatches in the priming region, as has been found in other studies of the Tetracinellida (Cárdenas *et al*., 2011). To circumvent this problem, I used the Illumina contigs and data from Genbank to design custom primers to amplify 700-850 bp regions of these loci (table S3). These worked well in nearly all of the samples recently collected by the author, but failed to amplify a product in many vouchers that were loaned from museums. This is likely because the DNA in older samples is degraded, either because of how they were collected, how they were initially preserved, how they have been archived since, or some combination of the three. Though the causes of this phenomenon are not well understood, many others have also reported difficulty in sequencing DNA from older sponges; more recently, sequencing of smaller fragments has yielded successes (Cárdenas & Moore, 2019; Agne *et al*., 2022; Erpenbeck *et al*., 2025). To further investigate, I designed primers to amplify subregions of the target loci, varying from 500 bp to only 100 bp in length. I attempted to amplify longer regions first, then progressively smaller ones if these failed. There is a clear relationship between sample age and the longest amplicon that was successfully sequenced for a sample (figure S10). This result should be considered hypothesis-forming rather than hypothesis-testing, as these points are non-independent due to partial correlations between age, collection location, collector, museum, and taxonomic subgroup. Several results are interesting to note, however. First, I was able to amplify small regions even from one of the oldest samples, the holotype of *T. villosa,* collected in 1879. This supports other findings that have successfully obtained DNA data from old type specimens (Agne *et al*., 2022; van der Sprong *et al*., 2025). Preservation of DNA was not universal, however, as I was unable to amplify even a 100 bp fragment from the type of *T. spinosa* which is from the same collection and a similar age, nor from either of the type specimens collected by M. W. de Laubenfels in the 1920s. This variable success has also been reported by others (van der Sprong *et al*., 2025).

The utility of sequencing small subregions of these genes was variable. In *Tetilla*, I designed primers to amplify specific SNPs that were diagnostic for species identification, which made even small fragments of DNA very useful. In contrast, when no a priori SNP targets were known, these DNA “mini-barcodes” (Cárdenas & Moore, 2019) sometimes failed to differentiate species that were morphologically distinct. For example, I successfully sequenced two small regions of the cox1 locus in most *Craniella* species; after removing primer sequence, about 200 bp of data remained from one amplicon and 100 bp of data remained from the other. At these 300 bp, *C. columbiana* sp. nov. and *C. hamatum* were indistinguishable from each other and from a sample of *C. wolfi* from the Galapagos Islands (figure S9). This is not surprising, as morphologically distinct sponge species have been found to be undifferentiated at even longer stretches of cox1 (Huang *et al*., 2008; López-Legentil *et al*., 2010; Pöppe *et al*., 2010; Turner & Pankey, 2023). More surprising is that *C. hamatum* and *C. columbiana* sp. nov. remained genetically indistinguishable in the partitioned analysis, which included 427 aligned base pairs from 28S. Only 230 of these base pairs were from the more rapidly-evolving D1D2 region, however, so it is likely that sequencing a larger portion of the locus would reveal genetic differences. These two species are very well differentiated morphologically, and I sequenced these loci with multiple sets of primers from two independent DNA isolations to ensure that this result was not due to contamination. These results support the idea that DNA is available in some old sponges, but that the utility of these small DNA fragments often depends on having newer material for comparison.

## Systematic Section

### Genus *Tetilla* Sollas, 1886

#### Diagnosis

Tetillidae without porocalices, and without a distinct, two-layered cortex, but may have perpendicular oxeas reinforcing the ectosomal layer. (Modified from previous definitions to include perpendicular ectosomal oxeas; Carella *et al*. 2016; Van Soest & Rützler 2002).

#### Diagnosis for Sediment-rooting *Tetilla*

*Tetilla* are found in two distinct types in the Northeast Pacific, one form I dub the “Sediment-rooting *Tetilla*” and the other that I dub the “Sponge-rooting *Tetilla*”. We will first discuss the “Sediment-rooting *Tetilla*”, which can be diagnosed as follows. Shape globular or clavate, rather than nearly spherical; lacking a palisade of stout oxeas II in ectosome; no well-developed radial skeleton.

### *Tetilla mutabilis* de Laubenfels, 1930

#### Material examined

Holotype: USNM 21498, Newport Harbor, California, near low tide mark, November 1924.

#### Diagnosis

No sigmaspires are present and the mean clad:rhabd angle of anatriaenes is > 40°.

#### Morphology

The holotype consists of a small, dull red fragment, of which a rectangular piece approximately 12 x 9 mm was loaned to me. The original description states that the sponge is found in a small “ pedunculate-clavate” form and a larger “irregularly massive” form, up to 8 cm high and 15 cm in diameter. These were dull red alive and in alcohol, possessing a roughly radial symmetry and an apical osculum. The surface was matted with brushes of spicules, especially prodiaenes and protriaenes. No root-like structure was described by de Laubenfels, who stated that “the massive forms lie loose on the soft mud, the clavate forms seem to have been attached to small shells or other solid objects” (de Laubenfels, 1932).

#### Skeleton

Not investigated in the small fragment of the holotype that was available. Originally described as having a central axis of densely packed spicules in the clavate form, but in the massive form, “no central or radiate skeleton, instead the spicules are matted together in sheets or walls around gross chambers” (de Laubenfels, 1932).

#### Spicules

As pointed out by de Laubenfels (1932), the spicules of this species are all very long, thin, and matted together, making the measurements of lengths difficult. Shown in figure 4, except when noted below.

**Figure 4.**
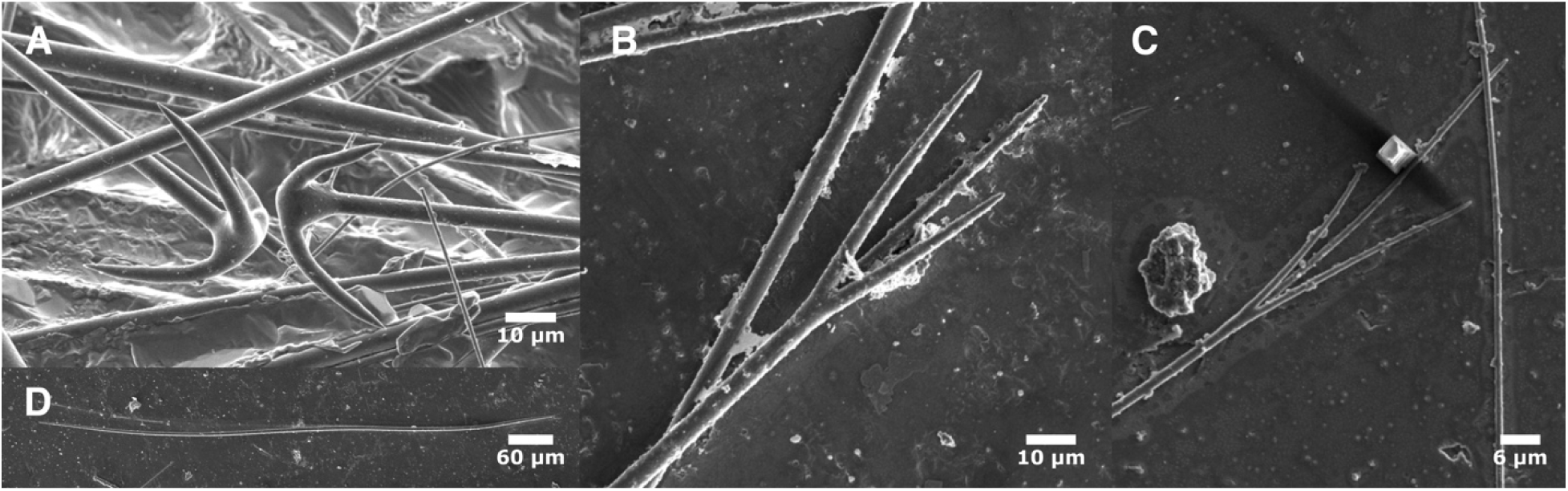
Tetilla mutabilis. A: anatriaenes; B: protriaene I; C: protriaene II. D: oxea II. All photos are of the holotype.

Oxeas I: the longest size-class of oxeas, usually tapering to filiform tips at both ends, but sometimes missing one or both filiform ends; easier to separate from oxeas II by length rather than shape. Longer examples likely occur. 1329–1496–1770 x 6–7–10 μm (n=5). Reported as 2000 x 2–9 μm in the original description. Not shown in figure 4; see figure 5D.

**Figure 5.**
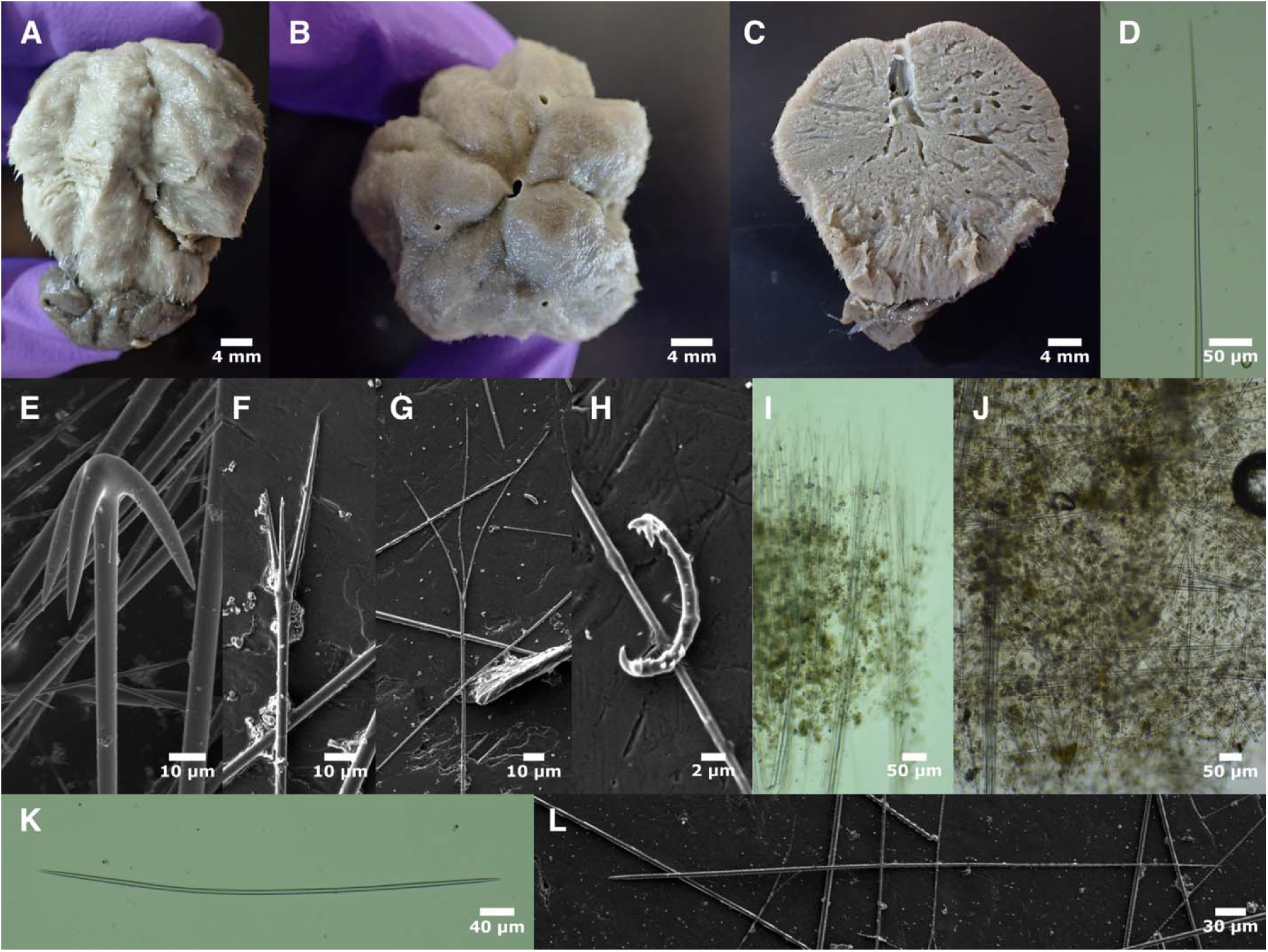
Tetilla japonica. A-C: sponge post-preservation. D: half of oxea I; E: anatriaene; F: protriaene I; G: protriaene II; H: sigmaspire; I: perpendicular surface section showing ectosomal skeleton; J: perpendicular surface showing choanosomal skeleton just below ectosomal region; K: oxea II; L: anisoform oxea III. All are from sample TLT1440.

Oxeas II: shorter than oxeas I and tapering abruptly to pointed tips, rather than filiform ends. 568–787–983 x 4–5–7 μm (n=24). Not reported in the original description.

Oxeas III: anisoxeas, with one end tapering abruptly to a pointed tip and the other tapering very gradually to a filiform end; thickest near the pointed tip. 514–739–969 x 3–4–4 μm (n=7). Not reported in the original description. Not shown in figure 4; see figure 5L.

Anatriaenes: most are slightly bulbus where the clads meet the rhabd. Clads are thin, round and abruptly tapering to points. The angle between clads and rhabd is large. No diaenes or monaenes were seen. Rhabd lengths not measured. Rhabd width 3–5–7 μm (n=25). Clads 22–33–45 x 3–5– 7 μm, with length:width ratios of 6–7–12 (n=25). Clad:rhabd angle 31°–47°–78° (n=40). The original description reports rhabds 3–7 μm wide, clads 20–30 μm long.

Protriaenes/prodiaenes I: those with clad widths over 2 μm were placed in this class. Clads are straight, only slightly spread, and average nearly equal in length. Most (88%, 15/17) were triaenes, while the others were diaenes. Rhabd: three that appeared unbroken were 1663–1893– 2022 μm in length; widths 3–4–5 μm (n=16). Longest clads 44–58–76 x 2–3–4 μm (n=16), with longest:shortest clad ratios of 1.0–1.2–1.5 (n=12). Clad:rhabd angle 8°–12°–18° (n=12). The original description reports all protriaenes/diaenes together with rhabds 1–6 μm wide, clads 30– 90 μm long.

Protriaenes/prodiaenes II: those with clad widths under 2 μm were placed in this class. Clads are hair-like, flexible, and often curved; only the longest clad was measured, due to difficulty in determining if any clads were broken. One of nine was a diaene, while the others were triaenes. Rhabd: 348–513–839 μm long (n=6); 2–2–3 μm wide (n=9). Longest clads 29–57–92 x 1–1–1 μm (n=7).

#### Distribution and habitat

Known with certainly only from the mud flats around Balboa Island in Newport Bay, Southern California. It was abundant there in November of 1924, absent in June of 1926, and abundant again in November of 1926 (de Laubenfels, 1932). I checked this location nearly 100 years later, in January of 2021, and found none. It has been reported from other places in Southern and Central California (Lee *et al*., 2007), but vouchers identified as this species, that I was able to acquire, were a better match to *T. japonica*, including two collected from Newport Bay. A field photo of sponges collected in San Diego Bay during an environmental survey in 2018 are dull red in color, as described by de Laubenfels, and might be this species (inaturalist.org/observations/176515329). It has also been reported that a brighter, reddish-orange *Tetilla* is sometimes abundant in Mission Bay, as shown in a photograph posted online (inaturalist.org/observations/6404736). I was unable to find any vouchers from this location, and no sponges were present when I checked this and other Mission Bay locations May 2021.

#### Remarks

The holotype is small and incomplete, did not yield amplifiable DNA even when amplicons as small as 100 bp were attempted, and I was unable to acquire any other sponges from this species. As a result, all I can add to de Laubenfels’s initial description of this species are more details regarding spicule types and dimensions. As sigmaspires can be rare in *Tetilla*, I searched extensively for them, and like de Laubenfels, found none. The lack of spicules thicker than 9 μm (10 μm in my reanalysis) was noted as distinctive by de Laubenfels, though this is also true of other *Tetilla* such as *T. japonica* Lampe, 1886, *T. serica* (Lebwohl, 1914), *T. hwasunensis* Shim & Sim, 2011, *T. limicola* Dendy, 1905, *T. pedonculata* Lévi, 1967, and *T. rodriguesi* Fernandez, Peixinho, Pinheiro & Menegola, 2011. Of these species, only *T. rodriguesi* also lacks sigmaspires; despite the similarity to *T. mutabilis* in terms of spicule dimensions, differences in gross morphology, climate, habitat, and ocean basin make these species unlikely congeners (Fernandez *et al*., 2011). Nonetheless, because it is possible that *Tetilla* are being transported long distances by humans, finding fresh samples of *T. mutabilis* for DNA sequencing are a high priority for future work.

### Tetilla japonica Lampe, 1886

#### Material examined

TLT1440, San Diego Bay, California, (32.70246, -117.16293), 9.05 m, 23-Aug-2023; SBMNH 47765, Newport Bay, California, (33.60000, -117.88330), depth not recorded, collection date not recorded, but donated to museum in 1976; SBMNH 659949, China Cove, Corona del Mar, California, (33.60000, -117.90000), 6 m, 8-Aug-1957; SBMNH 659950, San Clemente Island, California, (32.82390, -118.37920), in tidepool, Aug-1968.

#### Diagnosis

Sigmaspires are rare but present, and the mean clad:rhabd angle of anatriaenes is < 40°.

#### Morphology

The sequenced sample (TLT1440) is roughly spherical, but comprised of six lobes in a radially symmetric pattern. It is slightly taller (36 mm) than wide (30 mm at the top), with the widest point (35 mm) in the center, and a narrower (20 mm) base. The largest osculum (2 mm) is apical, but there are three additional oscula surrounding it that are about half the size. A root-like tuft of spicules and sand is attached to the base. Tan in ethanol; sample not seen alive by the author, but reported as purplish-red above and yellowish-green on the underside by the collector. The three other samples are tan to pinkish in color post-preservation. All are taller than wide, lobed like TLT1440, with the largest 67 x 38 mm and the smallest 29 x 21 mm. They have a single apical osculum and a basal root of spicules up to 117 mm long. All samples are moderately firm but somewhat compressible.

#### Skeleton

The interior lacks a strongly radial architecture. There are bundles of oxeas that travel towards the ectosome, but also many oxeas in confusion. At the ectosome, bundles of oxeas fan out slightly to form brushes, and are joined by anisoxeas (oxeas III) which were only found upright at the surface. Oxea bundles protrude only slightly, but the sponge surface is densely covered in protruding protriaenes I and II. Anatriaenes were not seen in the interior of the sponge nor in the ectosome on the top face of the sponge, but were present near the ectosome at the base of the sponge and in the root.

#### Spicules

Shown in figure 5, except when noted below.

Oxeas I: the longest size-class of oxeas, usually tapering to filiform tips at both ends, but sometimes missing one or both filiform ends. Separating from oxeas II based on shape was difficult, so oxeas were classified by splitting the bimodal length distribution at 1400 μm. 1422– 1575–1744 x 9–11–12 μm (n=18).

Oxeas II: shorter than oxeas I but otherwise similar; tending to taper abruptly to pointed tips, rather than having filiform ends. 442–897–1363 x 3–6–11 μm (n=81).

Oxeas III: anisoxeas, with one end tapering abruptly to a pointed tip and the other tapering very gradually to a filiform end; difficult to distinguish from oxeas II, but note that they are thickest near the pointed tip rather than in the center. 466–753–1090 x 3–5–7 μm (n=42).

Anatriaenes: one unbroken rhabd measured at 3571 μm in length, but tangled rhabds were likely longer. Rhabd width 3–5–7 μm (n=48). Clads 22–36–52 x 3–6–10 μm, with length:width ratios of 5–6–10 (n=25). Clad:rhabd angle 25°–35°–49° (n=57).

Protriaenes/prodiaenes I: those with clad widths over 2 μm were placed in this class. Clads are straight, only slightly spread, equal to fairly unequal in length. Most (73%, 8/11) were triaenes, while the others were diaenes. Three unbroken rhabds were 1260–1749–2438 μm in length, but broken rhabds were measured up to 2622 μm; widths 3–4–7 μm (n=16). Longest clads were 32– 70–123 x 2–2–4 μm (n=16), with longest:shortest clad ratios of 1.0–1.5–2.0 (n=7). Clad:rhabd angle 9°–13°–20° (n=6).

Protriaenes/prodiaenes II: those with clad widths under 2 μm were placed in this class. Clads are hair-like, flexible, and often curved; only the longest clad was measured, due to difficulty in determining if any clads were broken. Most (88%, 28/32) were triaenes, while the others were diaenes. Rhabd: 902–1366–1814 (n=6) x 1–2–4 μm (n=27). Longest clads 23–58–106 x 1–1–2 μm (n=26).

#### Distribution and habitat

The sequenced sample was collected in San Diego Bay at 9 m. If we are correct in assigning the other three samples to the same species, then it has also been found in Newport Bay and, surprisingly, a tidepool on San Clemente Island, 100 km offshore of mainland Southern California (this habitat seems poorly suited to the species, and I suspect the sample location may be erroneous). *Tetilla* with sigmaspires have also been reported from San Francisco Bay in Northern California, which might be this species (Hartman, 1975; Lee *et al*., 2007). For additional observations of *Tetilla* sp. in Southern California bays, see the *T. mutabilis* section.

#### Remarks

*Tetilla japonica* was described from Japan, and fits the description of the California samples, minus the spherasters, which were not reported by later authors and were presumably foreign (Lampe, 1886; Lebwohl, 1914). Two other species have very similar spicules, but differ in gross morphology. *T. serica* (Lebwohl, 1914) is cup-shaped, with a large cluster of oscula within the cup, and *T. hwasunensis* (Shim & Sim, 2011) is pink, cone-shaped, and has a large (5 mm) apical oscule. Moreover, a sponge from Thailand was identified as *T. japonica* by Sumait Putchakarn, and then sequenced at both the 28S and cox1 loci: our sample is identical at 28S (including the variable D1-D2 region), and differs by only one base pair at cox1 (Szitenberg *et al*., 2013). I therefore assign these California samples to this species. I acknowledge the possibility that both the California and Thai material is instead from an undescribed species similar to *T. japonica*, so it could be argued that *T. cf. japonica* is more appropriate. However, the extreme genetic similarity between the California and Thai samples makes it likely that the California sample is from a species also found in Asia, so a match to the morphological descriptions of *T. japonica* seems sufficient to place the California material in this species, pending further data.

Three *Tetilla* were available from the Santa Barbara Museum of Natural History that were similar to *T. japonica* / *T. mutabilis.* Two were collected in Newport Bay, the type location for *T. mutabilis*, and had been associated with that name; the third was misidentified as *T. arb*. Two clear spicular differences united these samples with *T. japonica* over *T. mutabilis*. First, they all possessed sigmaspires. Sigmaspires were quite rare in these samples, and finding them required considerable searching. This raises the possibility that the presence of sigmaspires is variable in *T. japonica*, which would mean *T. mutabilis* could be a junior synonym. Another spicular difference makes this less likely: the clads of the anatrienes are shaped differently in these species. In *T. mutabilis*, the clads point far out from the rhabd (somewhat like an umbrella), while in *T. japonica*, they curve down towards the rhabd (more like a dome umbrella). This difference can be quantified by measuring the angle from clad tip to top of spicule to center of rhabd (figure S5). This angle averaged 47° in *T. mutabilis* but only 35° in *T. japonica* (t-test p-value = 2.72e-11), and was consistent across all samples assigned to *T. japonica* (averaging 32°-37°). I attempted to confirm this morphological identification with DNA data from the older vouchers of this species from California, but succeeded only in obtaining a 100 bp amplicon from one Newport Bay sample and the San Clemente Island sample. After trimming primer sequence from the amplicon, a mere 40 bp of data remained. This sequence was identical to *T. japonica*, and did differ by one base pair from the most closely related species with data available (*T. radiata*) – but due to the short length, this is hardly conclusive.

These findings make it likely that *T. japonica* has been introduced to California by human activity. Introduction from California into Asia is also a possibility, but one made less likely by the fact that it was described from Japan in 1886 and the observation that California bays have some of the highest proportions of non-native species known (Ruiz *et al*., 2013). If validated by future work, this would make *T. japonica* the first known introduction of a sponge in the family Tetillidae.

Finally, I note that the anisoform oxeas III in this species appear to be limited to the ectosome. As the morphology of this species is otherwise quite similar to the type species for *Tetilla*, this highlights the need for ectosomal oxeas to be included in the diagnosis for the genus.

#### Diagnosis for sponge-rooting *Tetilla*

*Tetilla* are found in two distinct types in the Northeast Pacific, one form I dub the “Sediment-rooting *Tetilla*” and the other that I dub the “Sponge-rooting *Tetilla*”. We will now discuss the “Sponge-rooting *Tetilla*”, which can be diagnosed as follows. Spherical or nearly so, with a well-developed radial skeleton, but lacking a prominent double-layered cortex. Subectosomal layer contains an erect palisade of stout oxeas II and erect anisoform oxeas III. Protriaenes found in two size classes. Oxeas I are primarily symmetrical, with both ends filiform, rather than distinctly anisoform due to one filiform end as seen in most *Craniella*. Abundant anamonaenes usually seen in small individuals.

### *Tetilla arb* de Laubenfels, 1930

*Tetilla* sp. B (Hartman, 1975; Lee *et al*., 2007)

#### Material examined

Holotype: USNM 21496, Pescadero Point, Carmel, California, 36.56000, - 121.95000, Intertidal, July 1925. Other samples: TLT729, Purisima Point, California, 34.75593, - 120.63853, Intertidal, 13-Nov-2000; CASIZ 302844, East End of San Nicolas Island, California, 33.21833, -119.43433, 12 m, 27-Sep-2011; TLT207, Hazard Canyon, California, 35.28959, - 120.88415, Intertidal, 4-Jul-2016; 15491, Point Fermin, California, 33.70170, -118.29160, 12-15 m, 20-Aug-2019; TLT297, Halfway Reef, California, 33.76265, -118.42560, 15-23 m, 23-Aug- 2019; TLT470, Goalpost, California, 32.69438, -117.26860, 12-15 m, 8-Feb-2020; TLT539, Goalpost, California, 32.69438, -117.26860, 12-15 m, 8-Feb-2020; TLT669, La Jolla Cove Reef, California, 32.85227, -117.27239, 10-16 m, 14-Aug-2020; TLT671A, La Jolla Cove Reef, California, 32.85227, -117.27239, 10-16 m, 14-Aug-2020; TLT687a, Hazard Canyon, California, 35.28959, -120.88415, Intertidal, 12-Dec-2020; TLT826b, Cave Landings, California, 35.17535, -120.72240, Intertidal, 6-Feb-2021; TLT829b, Cave Landings, California, 35.17535, -120.72240, Intertidal, 6-Feb-2021; TLT833b, Cave Landings, California, 35.17535, -120.72240, Intertidal, 6-Feb-2021; TLT840b, Cave Landings, California, 35.17535, -120.72240, Intertidal, 6-Feb- 2021; TLT848b, Cave Landings, California, 35.17535, -120.72240, Intertidal, 6-Feb-2021; TLT938, Six Fathoms, California, 32.71000, -117.26860, 9-18 m, 15-May-2021; TLT948, La Jolla Cove Reef, California, 32.85227, -117.27239, 10-16 m, 17-May-2021; TLT1095A, Acropolis Steet, California, 36.64183, -121.93060, 9-18 m, 9-Aug-2021; TLT1100A, CRABS, Carmel, California, 36.55377, -121.93840, 10-17 m, 21-Sep-2021; TLT1205, CRABS, Carmel, California, 36.55377, -121.93840, 10-17 m, 21-Sep-2021; TLT1596A, Dutch Harbor, San Nicolas Island, California, 33.21600, -119.48413, 10-14 m, 10-Oct-2024.

#### Diagnosis

Yellow, black, or gray sponges, roughly spherical, 7–80 mm in diameter, with 1–10 discrete, separated oscula. Long-cladded anatriaenes (length:width ratios > 7.5) are absent. Small sponges have mainly anamonaenes, including “boathook” spicules. The larger size class of protriaenes include prodiaenes and promonaenes, with clads that are only slightly spread (mean angle < 17°) and modestly unequal in length (longest:shortest clad ratio means 1.1–1.7 per sample). The anisoform oxeas III are thinner than the isoform oxeas II. Oxeas I average less than 3000 μm long and oxeas II average less than 1300 μm long. This is the only *Tetilla* common in the intertidal; it is also found in the shallow subtidal. See table 2 for a comparison of key features with other sponge-rooting *Tetilla*.

#### Morphology

Spherical sponges with 1–10 scattered oscula, always discrete and well separated, which remain open and visible in preserved samples. Small sponges have a single apical osculum, while larger sponges vary. The oscula are sometimes surrounded by a tall palisade of spicules, but these are frequently absent from some or all oscula. Samples collected by the author were 7 to 55 mm across; the holotype is incomplete but is estimated to have been 72 mm across. Intertidal samples seen by the author were yellow, 7–20 mm across, were often seen in clusters, and were always attached to other species of sponges. The original description states that intertidal samples were drab and up to 80 mm wide, though at least one of these sponges was unattached and possibly washed up from the subtidal. Subtidal individuals are yellow, black, gray, or a blotchy mix of these colors; samples examined by the author were 10–55 mm across. Smaller sponges are usually yellow while larger ones are gray or black, but one 10 mm subtidal individual was black.

#### Skeleton

Skeleton radial, with bundles of spicules radiating from a central point; bundles can be straight, curved, or spiraling. Bundles contain oxeas I, anatriaenes, anadiaenes, and/or anamonaenes with the clad end near the surface of the sponge, and protriaenes, prodiaenes, and/or promonaenes with their clads near the surface or protruding. Oxeas and anatriaenes may protrude slightly, but most protruding spicules are protriaenes, prodiaenes, and/or promonaenes, which protrude both at spicule bundles and across the surface of the sponge. Anatriaenes, anadiaenes, and/or anamonaenes are more abundant in the lower portions of the sponge than they are in the upper portions of the sponge. They also protrude farther from the base of the sponge, where they are sunk into the host sponge as attachment points. The ectosome of the sponge is reinforced with oxeas II and oxeas III which form an erect palisade just under the surface of the sponge or protruding slightly. Smaller sponges are generally replete with anamonaenes, with anadiaenes and anatriaenes rare or absent, while this is reversed in larger sponges; some sponges of intermediate size have both anamonaenes and anadiaenes in abundance, and in one case, all three types in abundance. Sigmaspires occur throughout the sponge in moderate abundance.

#### Spicules

Shown in figure 6, except when noted below.

**Figure 6.**
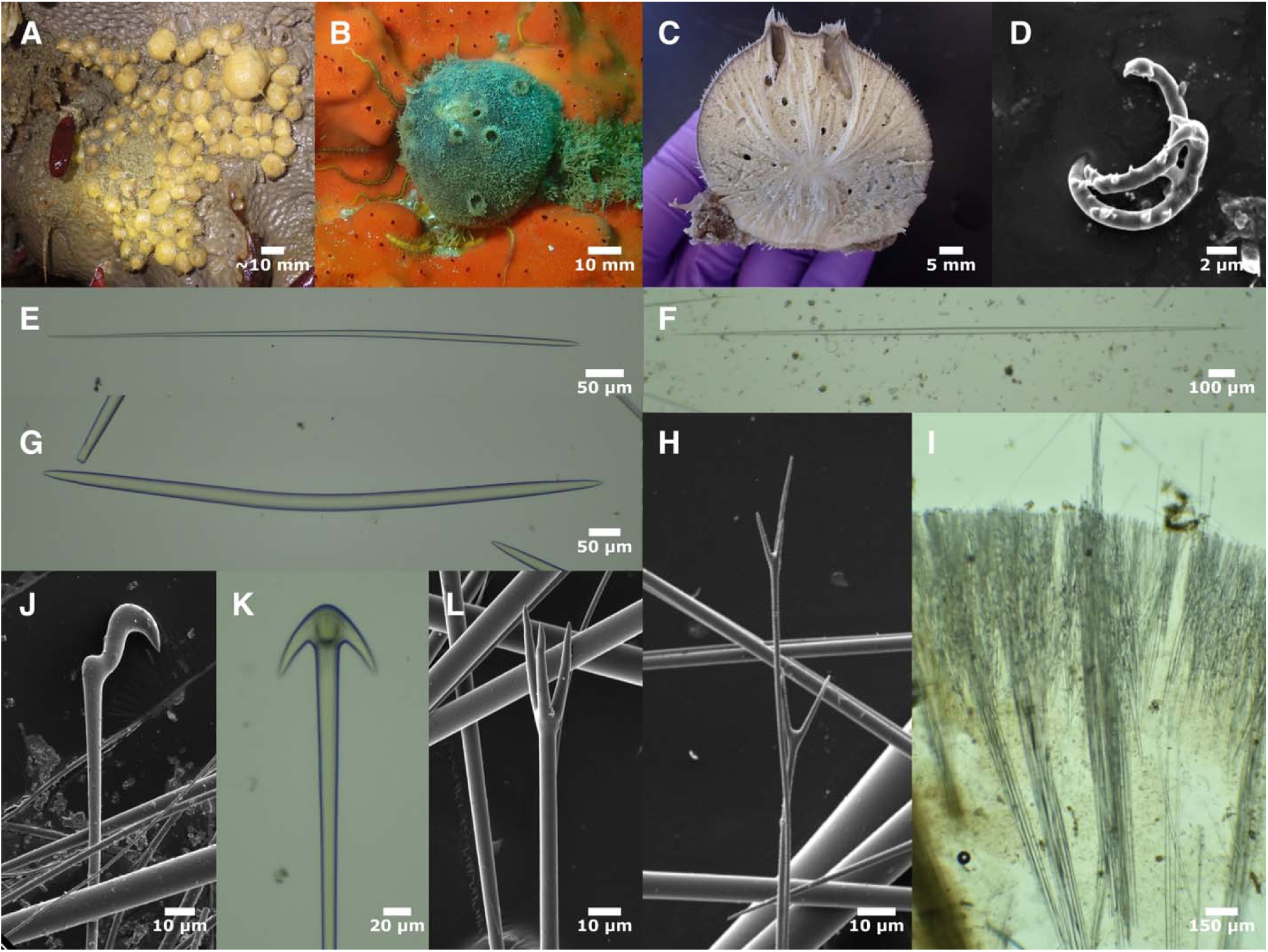
Tetilla arb. A: intertidal sample TLT207 on *Spheciospongia confoederata* (scale approximate, photo curtesy of Jeff Goddard); B: subtidal sample TLT1596A on *Dramgacidon mexicanum*; C: subtidal sample TLT1205 post-preservation, sectioned through oscula; D: sigmaspire from TLT1095A; E: anisoform oxea III from holotype; F: oxea I from holotype; G: oxea II from holotype; H: protriaene & prodiaene II from TLT1095A; I: perpendicular section through ectosome showing vertical oxeas II & III, protruding protriaenes, and anatriaenes (visible mainly in central spicule column) from TLT470; J: boathook anamonaene from TLT829B; K: anatriaene from holotype; L: protriaene I from TLT1095A.

Oxeas I: the longest size-class of oxeas. The largest are fusiform, tapering very gradually to filiform ends. Smaller ones are less dramatically fusiform, but still mostly symmetrical, thickest in the center, and gradually tapering. Some oxeas are hard to differentiate among categories, especially small oxeas I (especially if one filliform end is broken) and large oxeas III; see figure S11 for a comparison of oxea categories in the holotype. All sponges pooled: 477–2186–4479 x 4–22–44 μm (n=396). Mean length per sponge 1062–2633 μm, correlated with sponge size (figure 3).

Oxeas II: stout ectosomal oxeas, often curved or bent, tapering abruptly to sharp tips. One was seen that was modified into a style. All sponges pooled: 337–832–1570 x 5–18–44 μm (n=318). Mean length per sponge 479–1225 μm, correlated with sponge size.

Oxeas III: ectosomal anisoxeas, with one end tapering abruptly to a pointed tip and the other tapering very gradually to a filiform end; difficult to distinguish from oxeas I, but note that they are thickest near the pointed tip rather than in the center. All sponges pooled: 323–773–1605 x 3–9–19 μm (n=211). Mean length per sponge 434–1029 μm, correlated with sponge size.

Anatriaenes/anadiaenes/anamonaenes: Short and wide clads lead to a stout-headed architecture. Anamonaenes are most abundant in small sponges, anatriaenes most abundant in large sponges, and anadiaenes intermediate. Measures did not differ based on the number of clads, so all are pooled here. Rhabd 926–4856–10,296 (n=12) x 2–10–22 μm (n=146). Broken rhabds were measured up to 11,179 μm long. Clads 11–40–105 x 4–12–23 μm, with length:width ratios of 2– 3–7 (n=144). Clad:rhabd angle 16°–42°–56° (n=91).

Boathook anamonaenes: anamonaenes with the clad offset from the shaft. Similar in size to other anamonaenes, but measured differently due to shape (figure S7). One unbroken rhabd was measured at 2453 μm long; rhabd widths 3–6–9 μm (n=41). Length from clad tip to top of rhabd 21–30–41 μm (n=41).

Protriaenes/prodiaenes/promonaenes I: those with rhabd widths over 4 μm were placed in this class. Clads are straight, only slightly spread, averaging equal to modestly unequal in length. About half (58%, 69/120) were triaenes, while diaenes and monaenes were both common as well. The proportion of triaenes in any one sponge varied widely, from 0% to 86%. Rhabd 1007– 1952–3908 (n=14) x 4–6–14 μm (n=136). Longest clads were 15–47–87 x 2–4–9 μm (n=131), with longest:shortest clad ratios of 1.0–1.5–3.2 (n=108). Clad:rhabd angle 7°–15°–42° (n=109). Mean ratio per sponge varied from 1.1–1.7, while mean clad angle per sponge varied from 12°– 17°. Protriaene shown in figure 8L; for example prodiaene and promonaene, see figures 7K and 7L.

**Figure 7.**
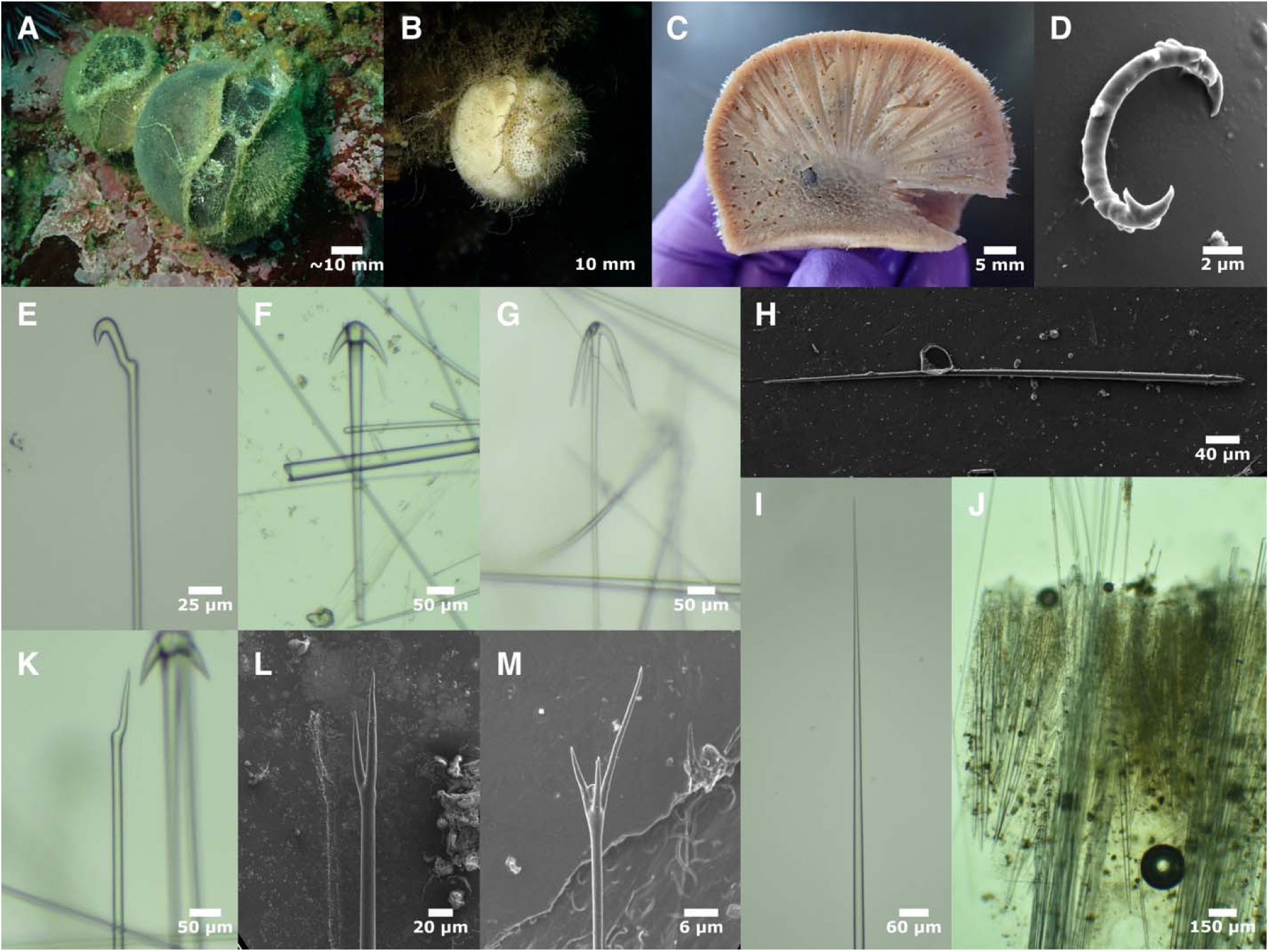
Tetilla villosa. A: uncollected sponges in situ at the collection location of TLT1203 (scale approximate); B: TLT942 on Geodia mesotriaena; C: RBC 007-00011-002, sectioned post-preservation; D: sigmaspire from TLT1203; E: boathook anamonaene from TLT1410; F: thick-cladded anatriaene from TLT1410; G: thin-cladded anatriaene from RBC 980-00335-008; H: anisoform oxea III from TLT943; I: tapered ending of oxea I from TLT1410; J: perpendicular section through ectosome showing vertical oxeas II & III, protruding protriaenes, and anatriaenes with clads inside and protruding, from TLT943; K: promonaene I from TLT1410; L: prodiaene I from TLT943; M: protriaene II from TLT943.

**Figure 8.**
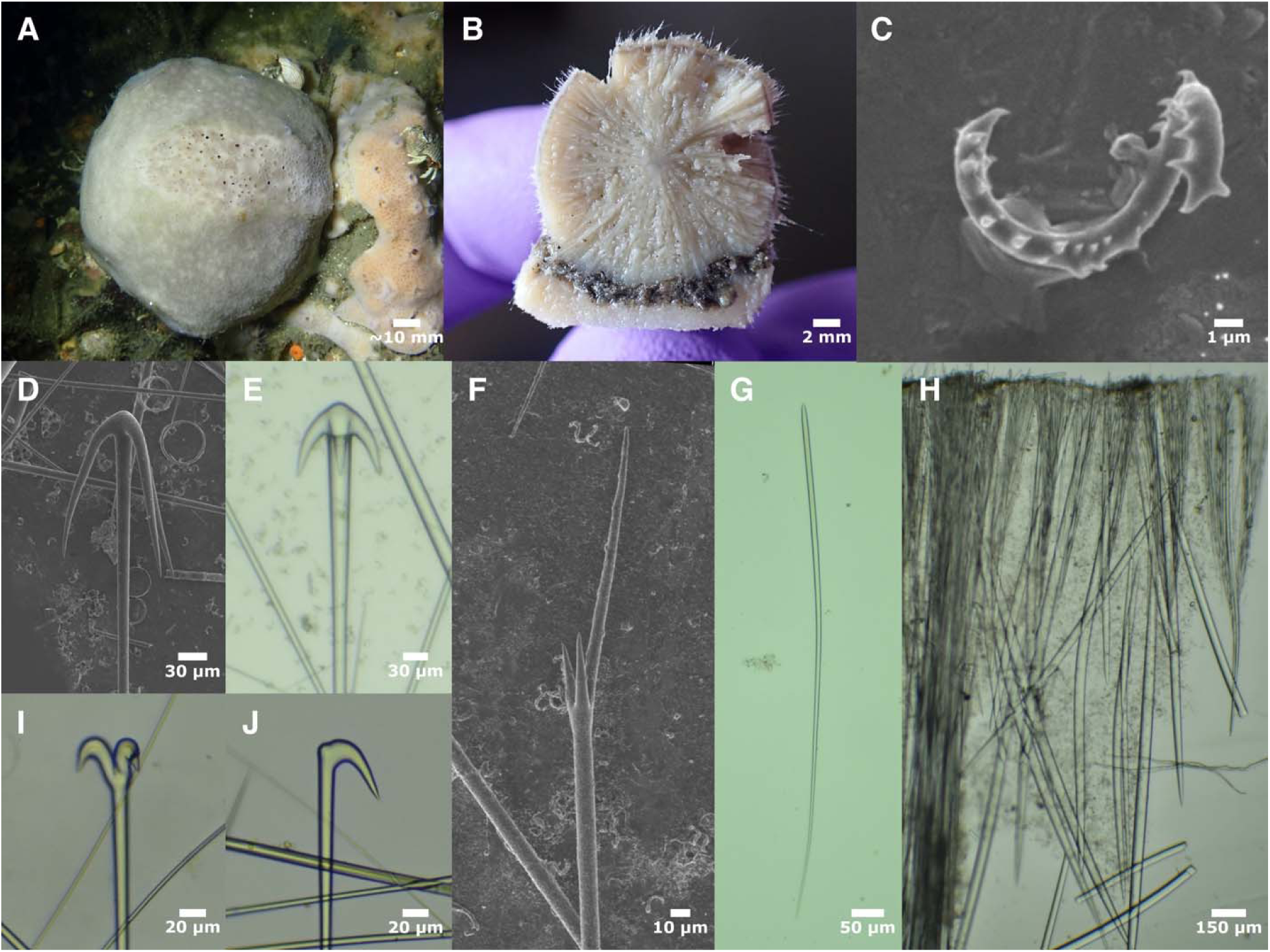
Tetilla vancouverensis. A: holotype in situ; B: sample TLT1311A sectioned post-preservation, with host sponge visible on the bottom; surface cavities are areas where spicules were sampled and not natural cavities; C: sigmaspire from holotype; D: thin-cladded anatriaene from holotype; E: thick-cladded anatriaene from TLT1623A; F: protriaene I from holotype; G: anisoform oxeas III from RBC 018-00212-001; H: perpendicular section through surface of RBC 018-00212-001, with a spicular bundle visible on the left side; I: anadiaene with clads at 90° from TLT1311A; anadiaenes with clads at 180° are more common, but both are seen regularly; J: anamonaene from TLT1311A; a spectrum of variation was seen from this “normal” shape to those more “boathook” shaped, as shown in figure 6J.

Protriaenes/prodiaenes II: those with rhabd widths under 4 μm were placed in this class. None were seen in the holotype fragment, but they were seen in all other samples investigated. Shaped similar to the larger class, but the clads are more spread apart and more unequal in length; promonaenes not seen. About half (60%, 83/138) were triaenes; the proportion of triaenes in any one sponge again varied, from 31% to 100%. Rhabd 323–726–2039 (n=18) x 1–2–4 μm (n=149). Longest clads were 8–29–49 x 1–2–3 μm (n=139), with longest:shortest clad ratios of 1.0–1.6–3.2 (n=106). Clad:rhabd angle 9°–21°–46° (n=93). Mean ratio per sponge varied from 1.2–2.5, while mean clad angle per sponge varied from 16°–24°.

Sigmaspires: curved in a C, S, or spiral-screw shape. Most have large spines scattered throughout, with the largest at the ends, but large end spines sometimes missing and some with only a few spines at all. Longest possible straight-line measurement across whole spicule 6–9–12 μm (n=75), with means per sponge 8–11 μm. Widths of shaft 0.9–1.3 μm (n=6)

#### Distribution and habitat

Found in the intertidal zone from Mendicino, Northern California, to Avila Beach, Central California; most abundant at the Southern end of this range, at locations in San Luis Obispo County. Found subtidally to at least 12 m depth, from Central California to Point Loma, Southern California (and probably farther north, but this is not confirmed). The distribution is patchy: it is common at shallow subtidal reefs around the Monterey Peninsula in Central California, and at Southern California sites around the Palos Verdes Peninsula and near San Diego, but not seen along the Santa Barbara coast despite considerable search effort. It was collected at several locations on San Nicolas Island, but was not seen at the 47 subtidal sites I searched on the other Channel Islands, and I know of no records or photos indicating it occurs on other islands.

Small individuals are found exclusively on other sponges, including members of the orders Dictyoceratida, Axinellida, Clionaida, Bubarida, Poecilosclerida, and Haplosclerida, and larger sponges are usually found attached to sponge hosts (host species are listed in table S1).

Some host sponges were extremely thin encrusting sponges, such as a *Hymedesmia* sp. that was a < 1 mm thick encrustation on a *Crassadoma gigantea* scallop valve, and hosted two large black *T. arb*.

All intertidal samples seen by the author, including those photographed by others and posted on iNaturalist, were small and yellow. The type sample, however, was a large “drab” sponge collected in the lower intertidal by de Laubenfels, so larger individuals are apparently accessible from the intertidal in some times and places.

#### Remarks

The original description of *T. arb* describes it as common in the intertidal in Central California, with distinct oscula surrounded by a palisade of spicules. Despite this being a good description for the samples described above, and not matching any other sponge known from the region, Hartman (1975) referred to the common intertidal *Tetilla* in Central California as *Tetilla* sp. B. This was due to several apparent differences between samples he examined and the type description of *T. arb*. Most importantly, de Laubenfels did not describe anamonaenes. This is now explained by this spicule primarily occurring in small individuals, though it remains unknown why de Laubenfels only encountered large (5-8 cm) sponges and not the smaller ones that are now the only ones present in the intertidal. Second, de Laubenfels described the oxeas I as “length certainly several millimeters, probably 2 or 3 cm”, which has led later authors to think that this species has extremely long oxeas (Lee *et al*. 2007; Lehnert & Stone 2011). My reexamination of the holotype finds these spicules to average 2.6 mm, to a maximum of 4.5 mm, so “cm” is a typo for “mm”. The results presented here clearly illustrate that only one species is common in the Central California intertidal, and that the spicules of the *T. arb* holotype are a very good match to large subtidal samples that genetic data confirm are of this same species. Interestingly, the de Laubenfels paratype (USNM 21497) is, in fact, a sample of *T. villosa*. This is surprising, because *T. villosa* does not have distinct oscula, nor is it known to occur in the intertidal; de Laubenfels mentions that one sample was found loose under a boulder, so perhaps this was a *T. villosa* that had washed ashore as beach wrack. Though he was familiar with the description of *T. villosa* from British Columbia, de Laubenfels does not seem to have seriously considered whether this species might occur in California (de Laubenfels, 1932).

*Tetilla arb* is differentiated from *T. villosa* and *T. vancouverensis* sp. nov. by lacking anatriaenes with clad length:width ratios > 7.5 (and often lacking anatriaenes all together), and by having 1–10 discrete, separated oscula. Also, the oxeas I average less than 3000 μm long and oxeas II average less than 1300 μm long, though this alone does not exclude small sponges of these other species. The holotype and paratype of *T. arb* are incomplete and lack the top portions of the sponge, so oscular morphology cannot be assessed. However, these spicular differences allowed me to confidently associate the *T. arb* holotype with the sponges described above, and associate the *T. arb* paratype with the species *T. villosa*, which I found to be present in the shallow subtidal in California. The presence of promonaenes and prodiaenes, and having anisoform oxeas III that are thinner than the isoform oxeas II, distinguish *T. arb* from *T. losangelensis* sp. nov. See table 2 for a comparison of the most important diagnostic features of sponge-rooting *Tetilla*.

### Tetilla villosa (Lambe, 1893)

#### Material examined

Holotype: CMNI 1900-2815, Houston Stewart Channel, Haida Gwaii, British Columbia, depth not recorded, 1879. Other samples: USNM 21497, Pescadero Point, Carmel, California, (36.56000, -121.95000), intertidal, 1-Jul-1925 (sample was previously the paratype of *Tetilla arb*); RBC 981-00216-015, Off Cape Sutil, British Columbia, (50.89167, - 128.05000), 20-25 m, 1981; RBC 980-00335-008, Harvey Cove, Mouth of Quatsino Sound, British Columbia, (50.43000, -127.92500), 15 m, 30-Jun-1980; RBC 007-00011-002, W. of Nootka Sound, British Columbia, (49.81667, -127.64333), 165 m, 6-Oct-1999; CASIZ 235692, 0.2 km NW of Boiler Bay RV Park, Oregon, (44.83191, -124.06830), 15 m, 1-Aug-2013; TLT1048, wreck of the USS Hogan, California, 37 m, 15-May-2021; TLT942 & TLT943, both collected at Lazy Days, San Diego, California, (32.69415, -117.27110), 12-25 m, 15-May-2021; TLT1203, CRABS, Carmel, California, (36.55377, -121.93840), 10-17 m, 21-Sep-2021; TLT1410, Tatoosh Island, Washington, (48.39370, -124.73305), 4-11 m, 31-Jul-2023.

#### Diagnosis

Discrete, separated oscula are not seen, and instead this species has an oscular area with many small openings covered by a pore sieve, usually surrounded by a prominent palisade of long protruding spicules. The oscular area is usually obvious in live samples, but frequently concealed in preserved samples, which can appear as featureless, fuzzy globes. Two types of anatriaenes are present: thick-cladded (ratios < 7.5) and thin-cladded (ratios > 7.5). A majority of protriaenes I are prodiaenes and promonaenes, with only 7%–40% protriaenes in any given sample; clads are not very spread (mean angle < 14°) and the longest:shortest clad ratio averages 1.1–1.7 per sample. The anisoform oxeas III are thinner than the isoform oxeas II.

#### Morphology

Holotype, and all sponges seen in situ, are spherical; some preserved samples were oblong or somewhat pyramidal. Known samples are 30–110 mm in diameter. Those seen in situ were gray or white, and retained their color in ethanol, but some preserved samples are beige. Sponges seen in situ had a large, obvious oscular area covered in a pore sieve (cribroporal oscula). In most cases, the oscular area was surrounded by a palisade of very long protruding spicules, though this was sometimes absent. The oscula were still visible on some preserved sponges, but could not be found on others, which also lacked the oscular fringe and resembled featureless, fuzzy globes. The basal portion of the sponge is covered in a mat of spicules and sediment, which is often anchored to another sponge; some sponges in situ did not appear to be attached to other sponges, and preserved samples with the basal mat may have been removed from a host or not. One sponge seen in situ had mats of apparently extruded spicules on some upper portions of the sponge as well, and some preserved samples consisted of two *T. villosa* matted together.

#### Skeleton

Skeleton radial, with bundles of spicules radiating from a central point that was closer to the bottom of the sponge than the top. The bottom of the sponge, below the gathering point of the bundles, was often softer and spongier than the top, but also cartilaginous and hard to cut through. Spicule bundles can be straight, curved, or somewhat spiraling. They contain oxeas I, anatriaenes/anadiaenes/anamonaenes with clads near the surface or slightly protruding, and protriaenes, prodiaenes, and/or promonaenes with clads mostly outside the sponge. Some spicule bundles were seen with both types of anatriaenes together, but sometimes spicules needed to be isolated from multiple regions of a sponge to find both types. Oxeas and anatriaenes sometimes protrude slightly, but most protruding spicules are protriaenes/prodiaenes/promonaenes, which protrude both at spicule bundles and across the surface of the sponge. The ectosome of the sponge is reinforced with oxeas II and oxeas III which are in an erect palisade just under the surface of the sponge or protruding slightly. Sigmaspires occur throughout the sponge.

#### Spicules

Shown in figure 7, except when noted below.

Oxeas I: the longest size-class of oxeas. The largest are fusiform, tapering very gradually to somewhat filiform ends. Smaller ones are less dramatically fusiform, but still symmetrical, thickest in the center, and very gradually tapering. Some oxeas are hard to differentiate among categories, especially small oxeas I, thin oxeas II, and large oxeas III. All sponges pooled: 429– 3626–6062 x 4–34–57 μm (n=202). Mean length per sponge 2549–4496 μm.

Oxeas II: stout ectosomal oxeas, often curved or bent, tapering abruptly to sharp tips. Rare styles and strongyles occasionally seen. All sponges pooled: 492–1156–2285 x 8–26–58 μm (n=293). Mean length per sponge 1039–1281 μm. Not shown in figure 7; see figure 6G for similar example.

Oxeas III: ectosomal anisoxeas, with one end tapering abruptly to a pointed tip and the other tapering very gradually to a filiform end; thickest near the pointed tip rather than in the center. All sponges pooled: 352–678–2429 x 5–9–23 μm (n=188). Mean length per sponge 506–881 μm.

Thin-cladded anatriaenes/anadiaenes: long and thin clads, nearly always triaenes. Rhabd 4989– 7041–9304 (n=4) x 7–12–21 μm (n=122). Many are tangled or broken, and much longer rhabds may occur. Clads 78–162–256 x 7–15–25 μm, with length:width ratios 8–11–20 (n=122). Clad:rhabd angle 16°–27°–42° (n=55).

Thick-cladded anatriaenes/anadiaenes/anamonaenes: Short and wide clads lead to a stout-headed architecture. Anatriaenes most abundant, but anadiaenes seen with the two clads either at 180° to each other or 90° to each other. A few anamonaenes seen (no small samples analyzed, where these are expected to be more common). Measures did not differ based on the number of clads, so all are pooled here. Rhabd 1644–7941–14492 (n=9) x 6–18–26 μm (n=180). Clads 27–88–165 x 7–22–39 μm, with length:width ratios 3–4–7 (n=180). Clad:rhabd angle 19°–37°–53° (n=79).

Boathook anamonaenes: anamonaenes with the clad slightly offset from the shaft. A single spicule of this type was seen in sample TLT1410, so these spicules may be common in smaller individuals when they are found.

Protriaenes/prodiaenes/promonaenes I: those with rhabd widths over 4 μm were placed in this class. Clads are straight, only slightly spread, averaging equal to modestly unequal in length. Only a minority (28%, 77/274) were triaenes, while diaenes and monaenes were both common. The proportion of triaenes in any one sponge varied but they were always in the minority (7%– 39%). Rhabd 1949–4259–6938 (n=16) x 5–10–16 μm (n=173). Longest clads 27–74–126 x 3–7–13 μm (n=167), with longest:shortest clad ratios 1.0–1.4–2.8 (n=135). Clad:rhabd angle 1°–12°– 22° (n=138). Mean ratio per sponge 1.1–1.7, while mean clad angle per sponge 10°–14°.

Protriaenes/prodiaenes II: those with rhabd widths under 4 μm were placed in this class. Shaped similar to the larger class, but the clads are more spread and less equal; promonaenes not seen. Unlike the larger class, triaenes were common (80%, 43/54). Rhabd 318–459–713 (n=16) x 1–2– 4 μm (n=80). Longest clads 10–25–51 x 1–2–5 μm (n=79), with longest:shortest clad ratios 1.0–2.1–4.8 (n=68). Clad:rhabd angle 12°–20°–35° (n=69). Mean ratio per sponge 1.7–2.8; mean angle per sponge 18°–21°.

Sigmaspires: curved in a C, S, or spiral-screw shape. Most have large spines scattered throughout, with the largest at the ends, but large end spines sometimes missing and some with few spines. Longest possible straight-line measurement across whole spicule 7–11–13 μm (n=87), with means per sponge 9–12 μm. Widths of shaft 1.0–1.7 μm (n=12).

#### Distribution and habitat

Previously known only from British Columbia, this species has a greater latitudinal and depth range that the other species in the region. It occurs from Haida Gwaii, British Columbia to San Diego, Southern California, from 10–165 m depth. One sample was apparently collected in the intertidal by de Laubenfels, but no other intertidal samples are known. It is rarely seen at diving depths: of the 123 subtidal sites I investigated between Juneau, Alaska and San Diego, California, I saw this species at only 4 locations. It was locally common, however, as multiple individuals were seen at 3 of these 4 sites. Several were found attached to other sponges including *Geodia mesotriaena* von Lendenfeld 1910 and *Suberites lambei* Austin *et al*. 2014, but most were not obviously attached to a host sponge.

#### Remarks

This species was well described by Lambe, and my analysis is highly congruent with his report (Lambe, 1893). The status and distribution of this species have been quite confused, however, as *T. villosa* outside British Columbia have previously been attributed to *T. arb*, and some British Columbia samples attributed to *T. villosa* are actually *T. vancouverensis* sp. nov.

*T. villosa* is differentiated from *T. arb* and *T. losangelensis* sp. nov. by its oscular morphology and the presence of thin-cladded anatriaenes. It has longer oxeas I and II than these species as well, but care is needed with this trait because it changes with sponge size. It is further differentiated from *T. losangelensis* sp. nov. by having promonaenes and prodiaenes, and by having the clads of these spicules be less spread (table 2). Differentiation of *T. villosa* and *T. vancouverensis* sp. nov. depends on the morphology of the large size class of protriaenes. In *T. villosa*, a minority of these are triaenes, with many monaenes, while in *T. vancouverensis* sp. nov., most of these are triaenes and there are no monaenes. Most samples can also be differentiated by the longest:shortest clad ratios of these spicules, but means for this trait are close to overlapping (table 2).

In California, *T. villosa* can be reliably identified in the field by its unique oscular morphology. It is likely that *T. villosa* and *T. vancouverensis* sp. nov. can also be differentiated in the field, but caution is needed, as they are more similar and fewer confirmed samples have been photographed. The oscular area on *T. vancouverensis* sp. nov. has many openings like *T. villosa*, but these appear larger than in *T. villosa*, lack the sieve-like covering, and either lack a spicular palisade or have one that is much reduced. In preserved samples, where the oscula are generally not visible, only the protriaenes (and the DNA) distinguish these species.

### *Tetilla vancouverensis* sp. nov

#### Material examined

Holotype: TLT1708A, Copper Cliffs, Quadra Island, British Columbia, (50.10362, -125.27982), 4-42 m, 27-Jul-2025. Paratypes: TLT1623A, Copper Cliffs, Quadra Island, British Columbia, (50.10362, -125.27982), 9-21 m, 26-Jul-2025; TLT1645, Copper Cliffs, Quadra Island, British Columbia, (50.10362, -125.27982), 4-42 m, 25-Jul-2025; TLT1688, Copper Cliffs, Quadra Island, British Columbia, (50.10362, -125.27982), 5.8 m, 25-Jul-2025. Other samples: RBC 981-00216-014, Off Cape Sutil, Nahwitti Bar, British Columbia, (50.89167, -128.05000), depth not recorded, 24-Aug-1981; RBC 018-00212-001, Point George, Shaw Island, Washington, (48.54720, -123.17550), 80-105 m, 13-Oct-2011; TLT1311A, Cape Alava, Washington, (48.17134, -124.75227), 4-12 m, 29-Jul-2023.

#### Diagnosis

Has many clustered, small oscula, not covered by a pore sieve. Possesses two types of anatriaenes: thick-cladded (ratios < 7.5) and thin-cladded (ratios > 7.5). The larger size class of protriaenes are primarily (>84%) protriaenes, with the minority being prodiaenes; promonaenes absent. These protriaenes have clads that are not very spread (mean angle < 15°), with clads that are unequal in length (longest:shortest clad ratios of 1.8–3.7 per sample). The anisoform oxeas III are thinner than the isoform oxeas II.

#### Etymology

Named for Vancouver Island.

#### Morphology

Spherical or nearly so; known samples are 18–95 mm in diameter. Those seen in situ were gray or white; gray is retained in ethanol but white turns to beige. A cluster of many oscula, 0.3–1.5 mm in size, is obvious in living samples but not in preserved material; this area may be weakly demarcated by a spicular palisade. One small sample, 18 mm in diameter, was much more hispid that the larger sponges, and appeared to have an apical oscular area, but oscula were concealed by protruding spicules. All sponges were found attached to other sponges, and often had a mat of spicules and sediment at the attachment point.

#### Skeleton

Skeleton strongly radial. All collected samples are fragments, except for the small sample, which has straight spicule bundles radiating from the approximate center of the sponge, with no apparent differences in the top and bottom halves. Spicule bundles contain oxeas I, anatriaenes/anadiaenes/anamonaenes, and protriaenes/prodiaenes. In the small sample, thick-cladded anatriaenes were mostly in the bottom of the sponge, where anamonaenes were also found, and thin-cladded anatriaenes were mostly in the top of the sponge. Anatriaene clads are found near the surface or protruding slightly, while most protruding spicules are protriaenes/prodiaenes, which protrude both at spicule bundles and across the surface of the sponge. The ectosome of the sponge is reinforced with oxeas II and oxeas III which are in an erect palisade just under the surface of the sponge or protruding slightly. Sigmaspires occur throughout the sponge.

#### Spicules

Shown in figure 8, except when noted below.

Oxeas I: the longest size-class of oxeas. The largest are fusiform, tapering very gradually to somewhat filiform ends. Smaller ones are less dramatically fusiform, but still symmetrical, thickest in the center, and very gradually tapering. Some oxeas are hard to differentiate among categories, especially small oxeas I, thin oxeas II, and large oxeas III. All sponges pooled: 721–

2720–5854 x 6–24–48 μm (n=96). Mean length per sponge 1802–4662 μm, correlated with sponge size. Not shown in figure 8; see figures 6F and 7I for indistinguishable examples.

Oxeas II: stout ectosomal oxeas, often curved or bent, tapering abruptly to sharp tips. Rare styles occasionally seen in most samples. All sponges pooled: 581–1248–2124 x 13–30–49 μm (n=76). Mean length per sponge 861–1537 μm, correlated with sponge size.

Oxeas III: ectosomal anisoxeas, with one end tapering abruptly to a pointed tip and the other tapering very gradually to a filiform end; thickest near the pointed tip rather than in the center. All sponges pooled: 509–911–1881 x 6–10–18 μm (n=64). Mean length per sponge 764–983 μm, correlated with sponge size.

Thin-cladded anatriaenes/anadiaenes: long and thin clads. Rhabd 1559–4219–7520 (n=14) x 7– 11–16 μm (n=122). Rhabds are usually tangled together and often broken, so much longer rhabds likely occur. Clads 74–151–273 x 7–13–22 μm, with length:width ratios 8–13–22 (n=123). Clad:rhabd angle 11°–26°–44° (n=73).

Thick-cladded anatriaenes/anadiaenes/anamonaenes: short and wide clads lead to a stout-headed architecture. Anatriaenes most abundant, but anadiaenes occasional. Anamonaenes common in lower portion of small sample, and occasional in other samples. Measures did not differ based on the number of clads, so all are pooled here. Rhabds tangled and often incomplete, but one was measured at 4192 μm in length. Widths 6–15–26 μm (n=59). Clads 22–60–102 x 6–17–31 μm, with length:width ratios of 3–4–7 (n=59). Clad:rhabd angle 28°–36°–45° (n=29).

Boathook anamonaenes: anamonaenes with the clad slightly offset from the shaft. Similar in size to other anamonaenes, but measured differently due to shape. Rhabd widths 9–13–16 μm, length from clad tip to top of rhabd 34–58–85 μm (n=12).

Protriaenes/prodiaenes I: those with rhabd widths over 4 μm were placed in this class. Clads are straight, parallel to rhabd or slightly spread, usually with one long clad or one short clad, and the other two of similar length. Prodiaenes rare (5%, 5/108), and only seen in 2/7 samples.

Anamonaenes not seen. Rhabd 1344–3209–4710 (n=13) x 4–10–16 μm (n=91). Longest clads 35–94–178 x 3–7–12 μm (n=89), with longest:shortest clad ratios of 1.3–2.5–5.3 (n=73). Clad:rhabd angle 4°–12°–21° (n=79). Mean ratio per sponge 1.8–3.7; mean clad angle per sponge 10°–14°.

Protriaenes II: those with rhabd widths under 4 μm were placed in this class. Shaped similar to the larger class, but the clads are more spread than the larger class; promonaenes and prodiaenes not seen. Rhabd 289–516–855 (n=9) x 2–3–5 μm (n=55). Longest clads 15–34–86 x 1–2–4 μm (n=54), with longest:shortest clad ratios 1.4–2.7–3.9 (n=33). Clad:rhabd angle 12°–20°–29° (n=44). Mean ratio per sponge 2.3–3.2; mean clad angle per sponge 15°–24°.

Sigmaspires: curved in a C, S, or spiral-screw shape. Most have large spines scattered throughout, with the largest at the ends. Longest possible straight-line measurement across whole spicule 7–10–16 μm (n=192), with means per sponge 9–10 μm. Widths of shaft 1.0–1.4 μm (n=3). Most (97%) of sigmaspires were between 7 and 13 μm, as seen in the other sponge-rooting *Tetilla* described here, with outliers present mainly in single sample (TLT1623A); these may have been foreign or a rare aberration.

One sample (RBC39) had a few spherical spicules (featureless, round balls; not shown).

#### Distribution and habitat

Known from the Northern tip of Vancouver Island to Cape Alava, Washington. They are abundant along the Quadra Island side of the Discovery Passage. Most were collected at diving depths, as shallow as 6 m, but one was dredged from 80–105 m. The Cape Alava sample was attached to *Obruta collector* Turner & Lonhart 2023, while all those sampled from Quadra Island were attached to *Scopalina jali* Turner 2021.

#### Remarks

*Tetilla vancouverensis* sp. nov. and *T. villosa* may have a parapatric distribution, as I have no verified samples of *T. villosa* from inside the Salish Sea, which is where most *T. vancouverensis* sp. nov. have been found. They do sometimes occur sympatrically, however, as they were collected together off Cape Sutil, North Vancouver Island, by Philip Lambert in 1981, and were both collected by me in the Olympic Coast National Marine Sanctuary in Washington.

The new species is visually and genetically distinct from *T. villosa*, but their spicules are extremely similar. These species can, however, be reliably differentiated by measuring the large size class of protriaenes. In this new species, these are nearly all triaenes, and there are no monaenes, while in *T. villosa*, a minority of these are triaenes, with many monaenes. Most samples are obviously different in the longest:shortest clad ratios of these spicules as well, but the species approach each other in a few samples (mean values of 1.1–1.7 in *T. villosa* and 1.8– 3.7 in *T. vancouverensis* sp. nov.; table 2). *T. vancouverensis* sp. nov. is differentiated from *T. arb* and *T. losangelensis* sp. nov. by its oscular morphology and the presence of thin-cladded anatriaenes. It is further differentiated from *T. losangelensis* sp. nov. by having protriaene clads that are less spread and having oxeas III that are thinner than oxeas II.

### *Tetilla losangelensis* sp. nov

#### Material examined

Holotype: NHMLA 15495 / BULA-0180, off old Marineland, Palos Verdes, California, (33.75170, -118.41670), 1-40 m, 21-Aug-2019.

#### Diagnosis

Spherical white sponge with a single osculum. Lacks thin-cladded anatriaenes. The larger size class of protriaenes are exclusively protriaenes, with clads that are moderately spread (mean angle = 20°). The anisoform oxeas III average thicker than the isoform oxeas II.

#### Etymology

Named for the city of Los Angeles.

#### Morphology

The only known sample is a white, slightly flattened sphere, 7.6 mm in diameter, with a single apical osculum approximately 0.5 mm across.

#### Skeleton

Skeleton strongly radial, with no distinct cortical layer. Tissues were not sectioned to examine the skeleton in detail due to the small size of the holotype. The spicule complement is the same as the three other sponge-rooting *Tetilla* described above, which do not differ in skeletal organization.

#### Spicules

Shown in figure 9, except when noted below.

**Figure 9.**
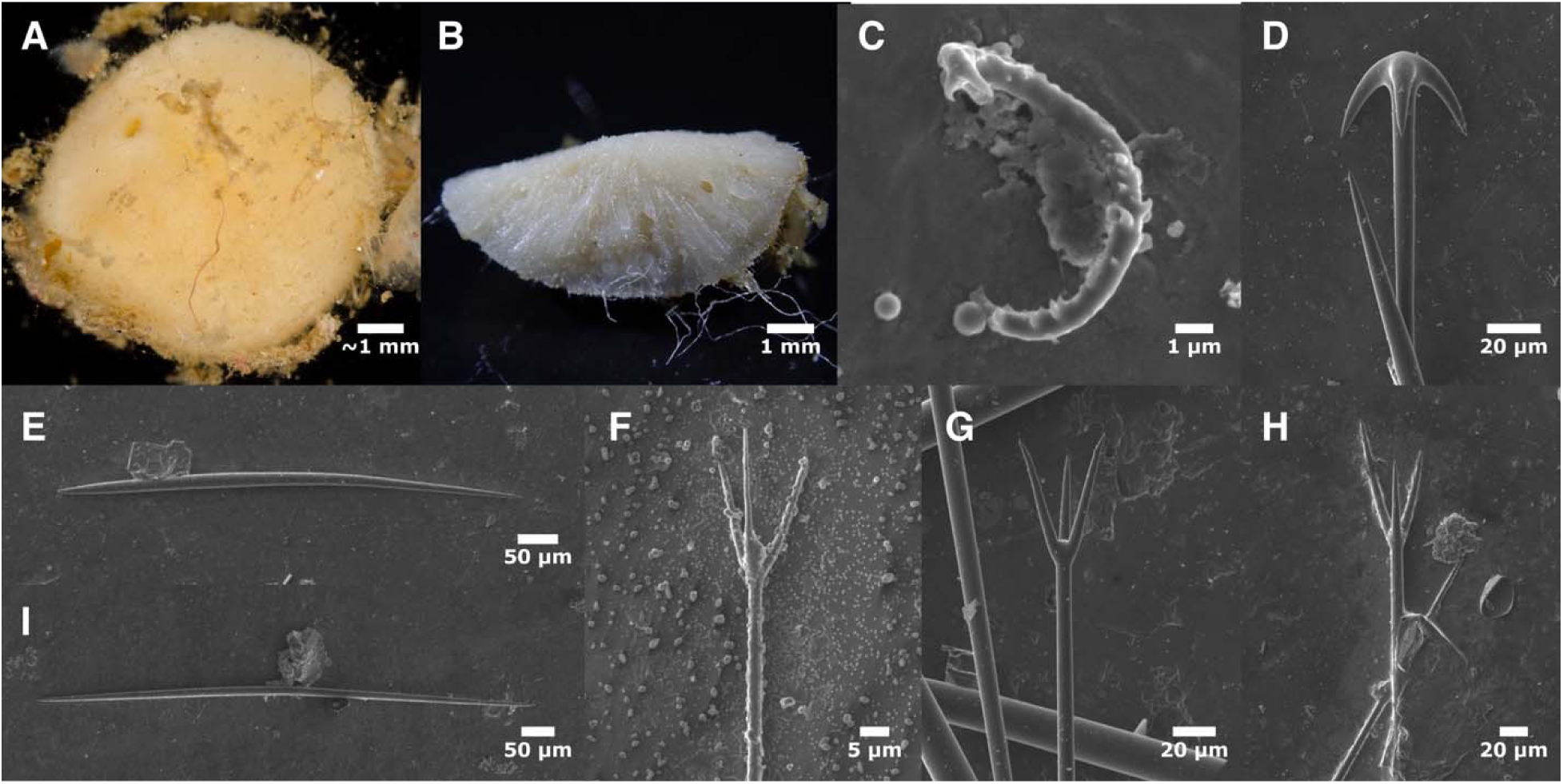
Tetilla l osangelensis. A: lab photo of sample after collection but before preservation (scale approximate); B: a subsampled portion post-preservation, with the ectosome facing down; C: sigmaspire; D: anatriaene; E: anisoform oxea III; F: protriaene II (partially encrusted with salt and/or debris); G-H: protriaenes I, both shapes typical; I: oxea II; all photos are of the holotype.

Oxeas I: the longest size-class of oxeas. Fusiform, tapering gradually to somewhat filiform ends. Smaller ones are less dramatically fusiform, but still symmetrical, thickest in the center, and very gradually tapering. Some oxeas are hard to differentiate among categories, especially small oxeas I and thin oxeas II. 1078–1574–2133 x 8–15–21 μm (n=23). Not shown in figure 9; see figures 6F and 7I for examples.

Oxeas II: ectosomal oxeas, often curved or bent, tapering abruptly to sharp tips; relatively thin on average, but differentiated from oxeas III by being symmetrical. One style seen. 336–745–973 x 6–13–21 μm (n=37).

Oxeas III: ectosomal anisoxeas, with one end tapering abruptly to a pointed tip and the other tapering gradually, often to a filiform end; thickest near the pointed tip rather than in the center. 405–681–1083 x 9–16–21 μm (n=35).

Anatriaenes/anadiaenes: Only one class seen; no anamonaenes found. Rhabds 1148–2091–3074 μm (n=10) x 4–8–11 μm (n=43). Clads 23–45–59 x 5–10–14 μm, with length:width ratios 3–5–6 (n=43). Clad:rhabd angle 31°–43°–57° (n=32).

Protriaenes I: those with rhabd widths over 4 μm were placed in this class. Clads are straight (figure 9H) or with tips that curve slightly away from the central axis (figure 9G), variably spread but averaging more than other sponge-rooting *Tetilla*. Clads are equal in length or modestly unequal. All (48/48) were protriaenes. Rhabd 1568–1940–2447 (n=8) x 5–9–14 μm (n=41). Longest clads 41–73–123 x 5–7–10 μm (n=36), with longest:shortest clad ratios of 1.0–1.4–1.9 (n=24). Clad:rhabd angle 13°–20°–29° (n=79).

Protriaenes II: those with rhabd widths under 4 μm were placed in this class. Shaped similar to the larger class, but the clads average more spread and slightly less equal in length. Rhabd 370– 587–749 (n=7) x 2–2–4 μm (n=23). Longest clads were 14–24–55 x 1–2–3 μm (n=23), with longest:shortest clad ratios of 1.0–1.3–1.8 (n=12). Clad:rhabd angle 19°–24°–32° (n=6).

Sigmaspires: curved in a C, S, or spiral-screw shape. Most have small or large spines scattered throughout, with the largest often at the ends. Longest possible straight-line measurement across whole spicule 7–9–12 μm (n=24). Widths of shaft 0.6–1.3 μm (n=10).

#### Distribution and habitat

The only known sample was collected by SCUBA divers off of the Palos Verdes Peninsula in Southern California, on a dive with a max depth of 40 m. Substrate unknown.

#### Remarks

Of the 46 sponge-rooting *Tetilla* investigated in this report, this sample was morphologically and genetically distinct, so it has been placed in its own species. In contrast to small individuals of *T. arb* and *T. vancouverensis*, this small sample lacked anamonaenes; it is not known if it was rooted in a host sponge. Combined with the lack of other samples, the lack of anamonaenes could indicate that this is not a small individual, but rather a typical individual from a small and cryptic species.

This species is differentiated from *T. villosa* and *T. vancouverensis* sp. nov. by having only thick-cladded anatriaenes and a single osculum. It is differentiated from *T. arb* and *T. villosa* by lacking prodiaenes and promonaenes. Unique among these *Tetilla*, the larger sized protriaenes are moderately flared (mean angle = 20°), and the anisoform oxeas III average thicker than the isoform oxeas II.

### Genus *Craniella* Schmidt, 1870

#### Diagnosis

Globular sponges without porocalices and with a distinct, two-layered cortex (visible to the naked eye). The outer cortical layer often has sub-dermal cavities with the inner layer made of collagen and a tight arrangement of cortical oxeas. Presence of direct-developing embryos within the sponge tissue. Paraphrased from Carella *et al*. (2016).

#### Regional diagnosis

Differentiating the *Craniella* cortex from the sponge-rooting *Tetilla* ectosomal layer can be difficult, but additional characters are useful within the temperate Northeast Pacific. *Craniella* oxeas I are usually anisoform, with a filiform tip at only one end. *Craniella* lack the small anisoform oxeas III present in sponge-rooting *Tetilla*; oxeas III, when present, are long, thin, and symmetrical. *Craniella* lack the small, thin protriaenes that are abundant in sponge-rooting *Tetilla*, though some have short-cladded forms. Most (but not all) *Craniella* also have protriaenes with widely-spread, equal-length clads and anatriaenes with pointed tops. Within the region, *Craniella* are not known south of British Columbia or from the intertidal zone.

Ten species of *Craniella* are present in the region, and table 3 summarizes the most important features for distinguishing them. These species fall into two clades in the phylogeny. One clade is more closely related to the North Atlantic *C. zetlandica* Carter 1872 than *C. wolfi* from the Galapagos Islands. This clade has considerable variation in sigmaspire morphology, and will be discussed first. The second clade, more closely related to *C. wolfi*, has little variation in sigmaspire morphology and will be discussed second; *C. spinosa* will be included here based on morphology. I will also provide morphological data on *C. craniana* de Laubenfels, 1953. This is an Arctic species, so not properly included in this revision. It is so poorly known, however, that I felt it necessary to examine the holotype in comparison to the other species, and I will include those values here. Figure 10 shows all verified collection locations for the 10 Northeast Pacific species.

**Figure 10.**
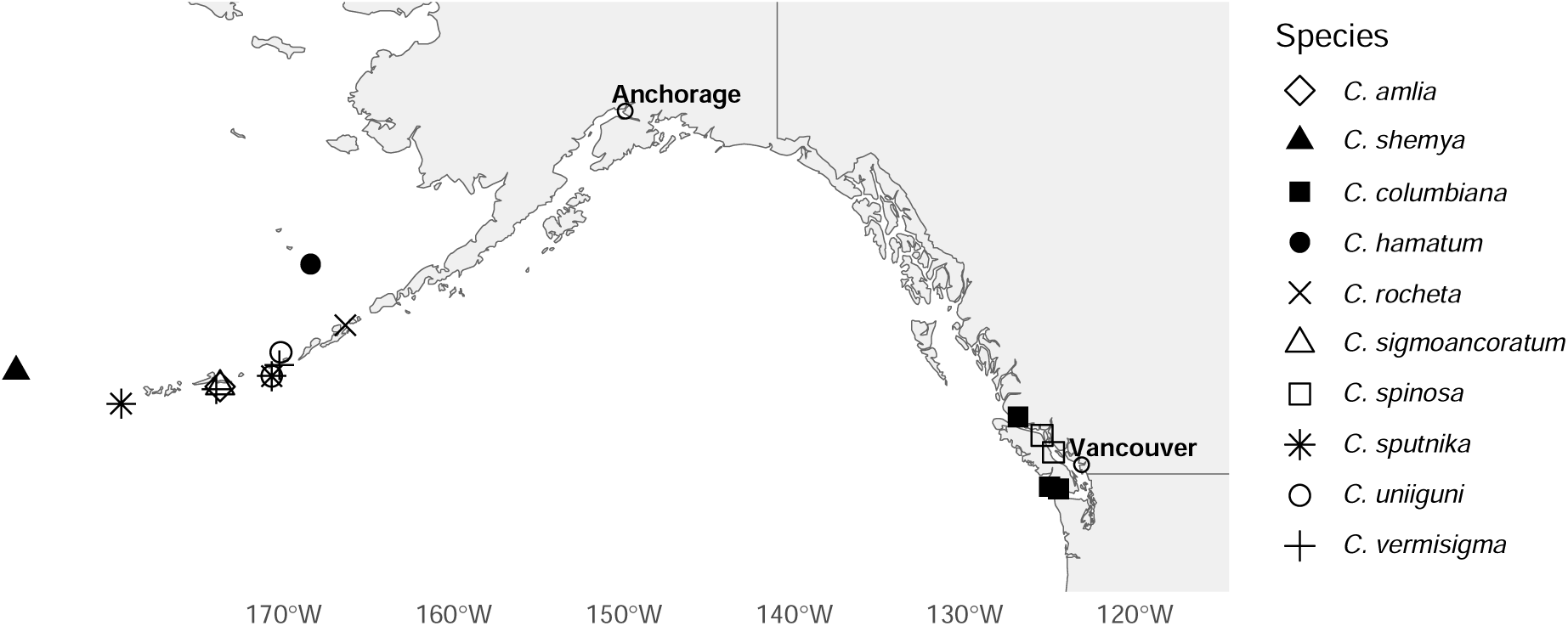
Range map of samples examined for all *Craniella* species.

### *Craniella sputnika* Lehnert & Stone, 2011

#### Material examined

USNM 1478668, S. of Islands of Four Mountains, Aleutian Islands, (52.28400, -170.66700), 157 m, 18-Jun-2012; USNM 1478669, S. of Islands of Four Mountains, Aleutian Islands, (52.27730, -170.65600), 165 m, 18-Jun-2012

#### Diagnosis

The only regional *Craniella* lacking sigmaspires and possessing centrotylote oxeas II.

#### Morphology

Spherical sponges covered in large conules with projecting spicule bundles. Sponge diameter 24–27 mm not including conules, up to 36 mm including conules. Double-layer cortex with subdermal cavities obvious in cross section, 1.0-2.5 mm thick. Cortex white and hard, interior yellowish-white and softer.

#### Skeleton

Skeleton strongly radial, with bundles travelling from a central point to the surface, either in straight lines or a spiral. Smaller, centrotylote oxeas II are found in the interior and cortex, while the larger, non-centrotylote oxeas II are only in cortex.

#### Spicules

I examined the spicules of one previously uncharacterized sample, USNM 1478668. For images of spicules, see Lehnert & Stone (2011).

Oxeas I: the longest size-class of oxeas. Fusiform, thickest in the center and gradually tapering at both ends, but anisoform due to one filiform end. 3141–4793–6565 x 36–54–74 μm (n=8). Type material previously described as 4530–5425 × 50–75 μm.

Oxeas IIa: stout cortical oxeas, fusiform and non-centrotylote. 489–772–1044 x 24–42–57 μm (n=19). Type material previously described as 540–987 × 28–63 μm.

Oxeas IIb: the smaller size class of stout oxeas, fusiform and with a ring-like centrotylote swelling. 109–246–391 x 7–16–31 μm (n=50). Type material previously described as 97–372 × 8–17 μm.

Anatriaenes: with short thick clads; most have a pointed top. One rhabd measured at 5408 μm long; widths 23–39–53 μm (n=18). Clads 81–171–280 x 23–38–50 μm, with length:width ratios of 3–4–6 (n=14). Clad:rhabd angle 36°–40°–56° (n=14). Type material previously described with rhabds 3430–8820 × 12–21 μm, clads 48–154 × 17–22 μm.

Protriaenes/prodiaenes: clads are straight, of equal lengths, and spread away from rhabd axis. Most were triaenes (10/11, 91%). Rhabd widths 18–30–43 μm (n=13). Longest clads 107–223– 351 x 9–20–34 μm (n=13), with longest:shortest clad ratios 1.0–1.1–1.3 (n=9). Clad:rhabd angle 14°–19°–22° (n=10). Type material previously described with rhabds 4520–8960 × 12–40 μm, clads 50–250 × 5–17 μm.

Short-cladded protriaenes: shaped like long-cladded protriaenes, but with a greater angle; less common and possibly immature forms of the common type. Rhabd widths 12–13; longest clad 37–44 x 7–8 μm, with longest:shortest clad ratios of 1.2. Clad:rhabd angle 33°–38° (n=2 for all). Not described from type material; see figures 14F & 19C for similar examples.

#### Distribution and habitat

Known from deep water (115–165 m) in the Central Aleutian Islands. Described from two samples collected near Amatignak Island. The samples examined here were collected South of the Islands of Four Mountains. Some, and possibly all, of these four known samples were attached to other sponges when they were collected.

#### Remarks

This species is easily identified from its spicules by the lack of sigmaspires and the presence of a small category of centrotylote oxeas. It is further differentiated from some other species by having long protriaene clads, no thin-cladded anatriaenes, and thick-cladded anatriaenes with a large clad:rhabd angle (table 3). *C. sputnika* was recently described, including SEM images, and the sample examined here was a good match to the species description.

### *Craniella vermisigma* sp. nov

#### Material examined

Holotype: USNM 1478660, S. of Islands of Four Mountains, Aleutian Islands, (52.64820, -170.21900), 235 m, 18-Jun-2012. Paratype: USNM 1478700, S. of Amlia Island, Aleutian Islands, (51.84280, -173.91000), 195 m, 6-Jul-2004.

#### Diagnosis

The finely-spined sigmaspires are unique in the region, but this is only visible with scanning electron microscopy. With light microscopy, sigmaspire length easily differentiates this species from all except *C. shemya* sp. nov. and *C. uniiguni* sp. nov. Protriaene and anatriaene clad lengths are much shorter in *C. shemya* sp. nov., and many characters are different in *C. uniiguni* sp. nov. including protriaene clad length and two types of oxeas II (table 3).

#### Etymology

Named for its unique sigmaspires, which look like fuzzy worms.

#### Morphology

Roughly spherical sponges, slightly taller (40–65 mm) than wide (54–36 mm. A basal spicule mat was present in one sample, while the other had a slightly flattened area that may have been the base. Covered in small conules, but only a few projecting spicules are visible. Oscula not apparent, though one sponge has a large area lacking conules that appeared to be the upper surface based on internal anatomy. Sectioning revealed a cartilaginous cortex layer about 1 mm thick, with a thinner dermal layer above it and a few open sub-dermal spaces. Brooded larvae or direct-developing juveniles, up to 2 mm across, were numerous deep within the sponge tissue. White alive; tan or chocolate brown post-preservation.

#### Skeleton

Skeleton strongly radial, with bundles of oxeas I travelling from a central point to the ectosome in curving bands. Anatriaenes and protriaenes also incorporated into bundles, with clads near the sponge surface, and both are present in the basal spicule mat. Protriaenes are more common on the upper surface of the sponge but also present in the lower surface. Cartilaginous cortex layer filled with oxeas II. Oxeas II are upright or angled up to 45° to surface, but not tangential. Dermal layer contains only sigmaspires, which are abundant throughout the sponge.

#### Spicules

Shown in figure 11, except when noted below.

**Figure 11.**
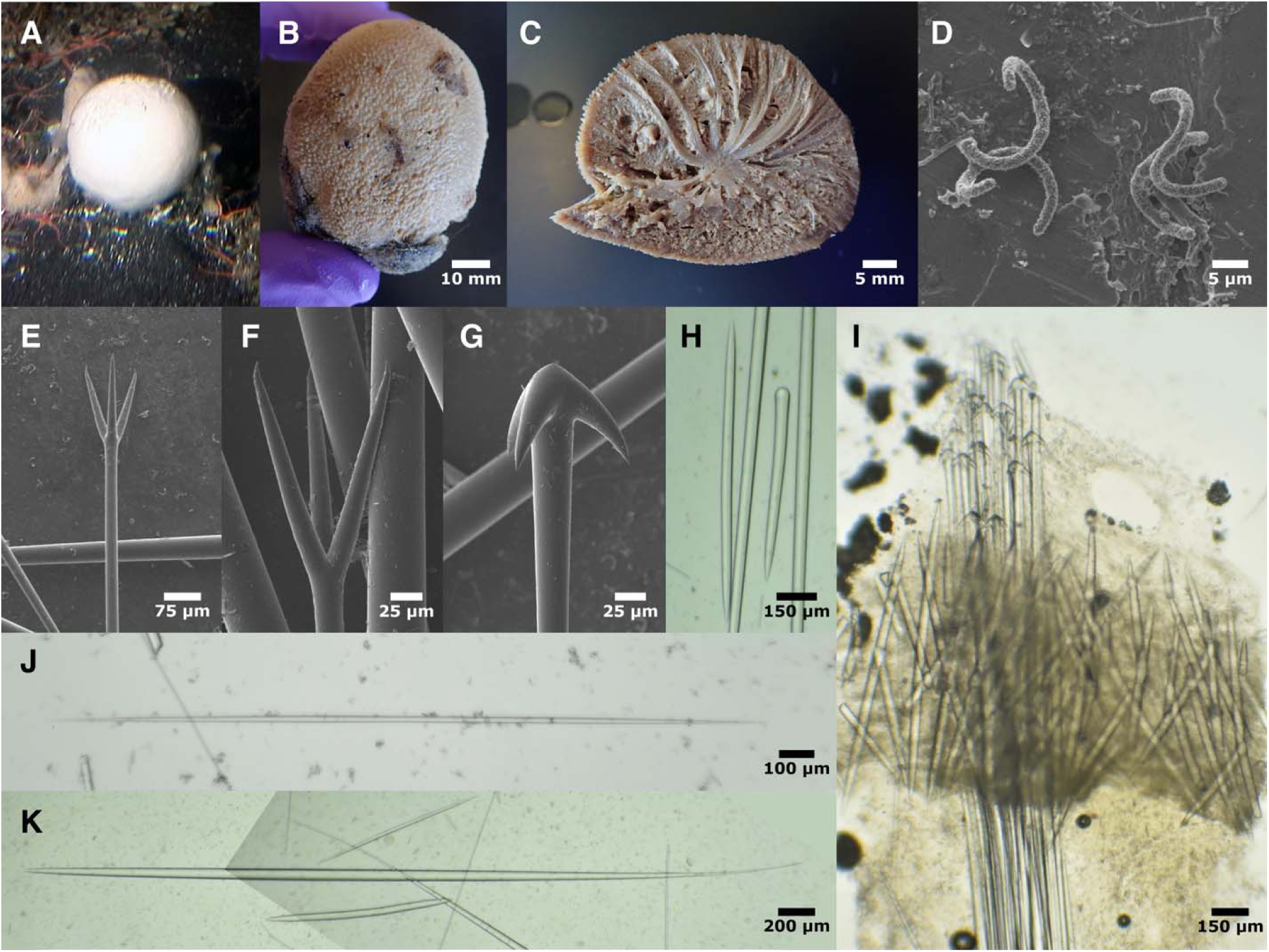
Craniella vermisigma. A: in situ photo of USNM 1478700 (curtesy of Robert Stone, NOAA); B: USNM 1478700 post-preservation; C: bisected holotype with juveniles visible; D: sigmaspires from USNM 1478700; E: protriaene from USNM 1478700; F: close-up view of protriaene showing bent tips from USNM 1478700; G: anatriaene from USNM 1478700; H: oxea II and style from USNM 1478700; I: perpendicular section through ectosome, sample USNM 1478700; J: oxea III from holotype; K: oxea I from holotype (composite of 2 images).

Oxeas I: the longest size-class of oxeas. Fusiform, thickest in the center and gradually tapering at both ends, but anisoform due to one filiform end. 2850–4248–4793 x 38–50–59 μm (n=28).

Oxeas II: stout cortical oxeas. 562–1057–1365 x 17–36–45 μm (n=54).

Styles: similar in size to oxeas II but less common. Straight or slightly subtylote. 657–796–1064 x 28–41–47 μm (n=10).

Oxeas III: long, thin, symmetrical oxeas; differentiated from oxeas I & II by being symmetrical and by being thinner relative to their length. 1027–2173–3641 x 6–18–34 μm (n=13).

Anatriaenes: with short thick clads; most have a pointed top. One anamonaene seen. Rhabds 4498–9528–16,067 (n=10) x 22–32–41 μm (n=46). Clads 79–114–147 x 24–34–45 μm, with length:width ratios of 2–3–4 (n=45). Clad:rhabd angle 24°–35°–47° (n=46).

Protriaenes/prodiaenes: clads appear straight in the light microscope, but all those examined under SEM were straight until the tips, which were angled in slightly. Clads of equal lengths, spread away from rhabd axis. Most were triaenes (28/29, 97%). Rhabds 4562–5818–7477 μm (n=10) x 16–22–26 μm (n=39). Longest clads 114–161–236 x 11–16–20 μm, with longest:shortest clad ratios 1.0–1.1–1.2 (n=38). Clad:rhabd angle 7°–20°–28° (n=39).

Sigmaspires: curved in a C, S, or spiral-screw shape. Evenly covered in very small spines. Some appeared to be very subtly centrotylote, but this was not visible in all spicules. Longest possible straight-line measurement across whole spicule 11–15–23 μm (n=91).

#### Distribution and habitat

Known from two samples, collected 8 years and 250 km apart in the Central Aleutian Islands. Trawled from deep water (195–235 m).

#### Remarks

This species is clearly differentiated from others in the region by the finely-spined sigmaspires, but this trait is only visible with scanning electron microscopy. The sizes of the sigmaspires, protriaene clads, and anatriaene clads can easily identify this species under light microscopy (see diagnosis and table 3).

The characteristics of all *Craniella* known from the Northern Hemisphere were helpfully compiled by Lehnert and Stone (2011), and the only Northern Hemisphere species described since are from the Galapagos Islands. As shown in their table, all species except *C. cranium* (Müller, 1776), *C. prosperiaradix* Tanita & Hoshino 1989, and *C. globosa* Thiele, 1898 can be excluded based on the size of the sigmaspires. *C. cranium* is an Atlantic species, and reports of its presence in the North Pacific are likely erroneous; in any case, these reports describe a sponge with much shorter anatriaene and protriaene clads (Tanita & Hoshino, 1989). *C. prosperiaradix* is excluded based on the presence of additional types of microscleres and *C. globosa* is excluded based on very short oxeas I and anatriaenes and protriaenes with much shorter clads and rhabds. Interestingly, a species recently described from the Galapagos Islands (*C. wolfi*) has finely-spined, worm-like sigmaspires resembling those of *C. vermisigma* sp. nov., though they are extremely long (62–91 μm) (Schuster *et al*., 2018). Genetic data place them in separate clades, so this finely-spined morphology has evolved convergently (figure 1).

### Craniella sigmoancoratum (Koltun, 1966)

#### Material examined

USNM 1478663, S. of Amlia Island, Central Aleutian Islands, (51.93450, - 173.71100), 103 m, 30-Jun-2012.

#### Diagnosis

The unique C-shaped sigmaspires with end denticles are much larger than any other species in the region.

#### Morphology

The only voucher examined contains five small, spherical sponges covered in conules with projecting spicules bundles. The largest is 6.5 mm in diameter; 8.8 mm including the projecting bundles. The outer 0.5–1.0 mm is a white cortical layer, though not clearly “double layer” to the naked eye; choanosome light yellow post-preservation. Beige when first collected. The original species description states that samples up to 50 mm occur.

#### Skeleton

Skeleton radial, with bundles of oxeas I and protriaenes travelling from a central point and extending into spines. Anatriaenes not seen in tissue sections, but presumed to be present in spicules bundles as seen in all related species. Sponge surface is a thin dermal layer containing only sigmaspires, which are also present in all other parts of the sponge. Supporting this layer is a thick layer of oxeas II at various angles from upright to nearly tangential.

#### Spicules

Shown in figure 12, except when noted below.

**Figure 12.**
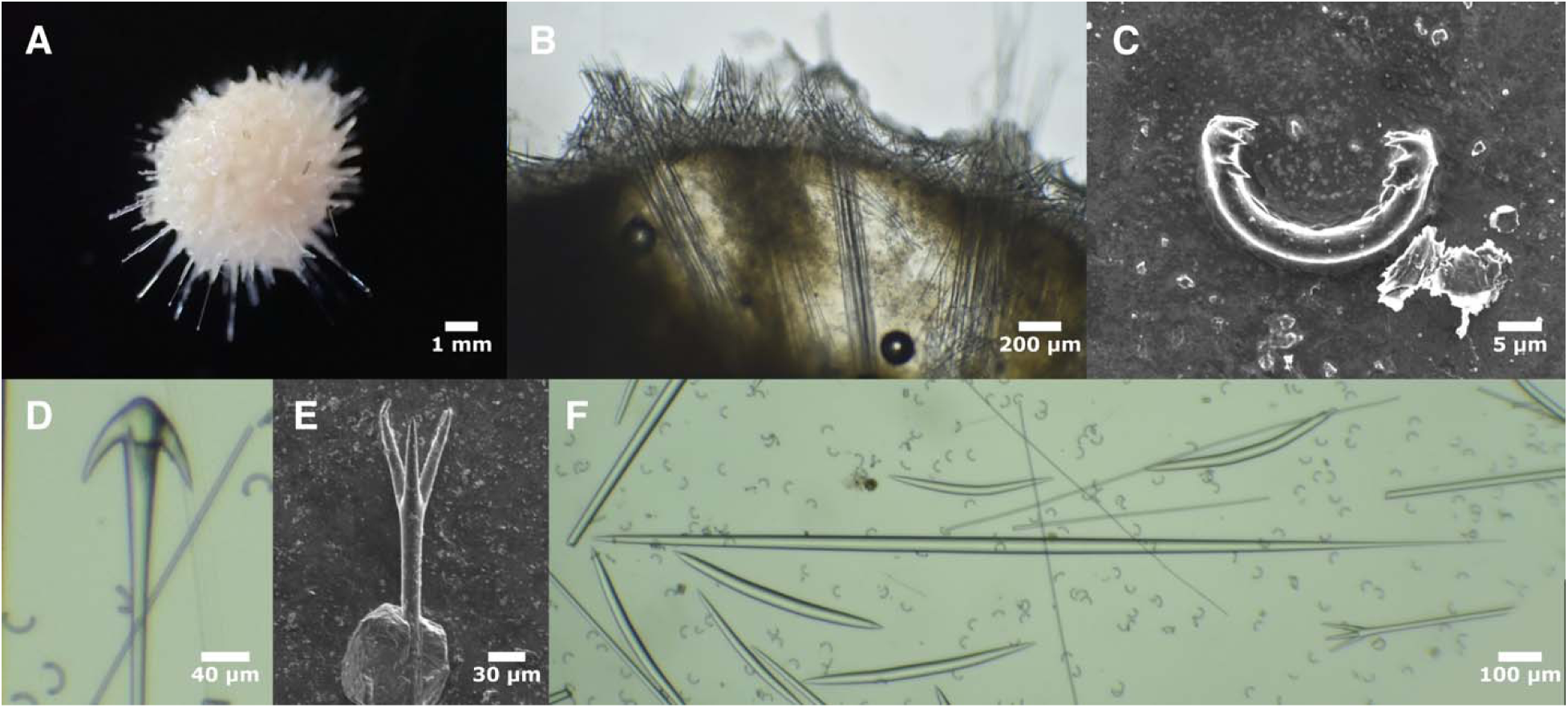
Craniella sigmoancoratum. A: Whole individual, post-preservation; B: perpendicular section through ectosome; thin dermal layer has broken away on the left, but is visible on the right as a thin layer of sigmaspires draped across the cortical oxeas II; C: sigmaspire; D: anatriaene; E: protriaene; F: anisoform oxea I surrounded by oxeas II and sigmaspires, with a single protriaene. All photos from USNM 1478663.

Oxeas I: the longest size-class of oxeas. Fusiform, thickest in the center and gradually tapering at both ends, but anisoform due to one filiform end. 1048–1597–2150 x 19–27–36 μm (n=16). Type material previously described as 3350–7100 x 50–78 μm, but discrepancy could be explained by differences in sponge size.

Oxeas II: stout cortical oxeas. 372–437–576 x 14–23–31 μm (n=24). Type material previously described as 460–1340 x 26–67 μm; differences may be explained by differences in sponge size.

Oxeas III: long, thin, symmetrical oxeas; differentiated from oxeas I by being shorter and symmetrical, and from oxeas II by being thinner relative to their length. Only one seen, 1125 x 5 μm. Not described from type material. Not shown in figure 12; see figure 14K & 16J for similar examples.

Anatriaenes: with short thick clads; most have a pointed top. Rhabd length not measured, as all spicules were broken; widths 14–21–26 μm (n=21). Clads 41–80–94 x 14–24–30 μm, with length:width ratios 3–3–4 (n=21). Clad:rhabd angle 25°–35°–42° (n=21). Type material previously described with rhabds 3000–8700 x 13–40 μm, clads 26–94 μm; type illustration shows an example with a clad angle of 30°.

Protriaenes: clads are straight, of equal lengths, and spread away from rhabd axis. All were triaenes. Rhabd length not measured, as all spicules were broken; widths 13–17–22 μm (n=21). Longest clads were 61–113–149 x 8–11–15 μm, with longest:shortest clad ratios of 1.0–1.1–1.4 (n=20). Clad:rhabd angle 17°–22°–29° (n=21). Type material previously described with clads 87–235 μm (rhabd dimensions not reported or not included in translated version available from the Fisheries Research Board of Canada). Type illustration shows equal length clads, with an angle of 23°.

Sigmaspires: C-shaped, with unspined shafts and end denticles. Longest possible straight-line measurement across whole spicule 25–27–29 x 3–3–4 μm (n=20). Type material previously described as 22–34 μm.

#### Distribution and habitat

Known from deep water (100–188 m) from the Southern Kuril Islands to Amlia Island in the Central Aleutians.

#### Remarks

This species is easily identified from its large, uniquely shaped sigmaspires. The spicules from the sample examined were a good match to the type material previously described from the nearby Kuril Islands.

### *Craniella uniiguni* sp. nov

#### Material examined

Holotype: USNM 1478662, N. of Islands of Four Mountains, Aleutian Islands, (53.08190, -170.13500), 186 m, 16-Jun-2012. Paratype: USNM 1478664, S. of Islands of Four Mountains, Aleutian Islands, (52.27730, -170.65600), 165 m, 18-Jun-2012.

#### Etymology

Named in honor of the Uniiĝun people who are native to the Islands of Four Mountains.

#### Diagnosis

The size of the sigmaspires differentiates *C. uniiguni* sp. nov. from all local species except *C. vermisigma* sp. nov. and *C. shemya* sp. nov. The species is differentiated from these two (and most others) by its protriaenes, which have very long clads, of somewhat unequal lengths, only modestly spread. The two length categories of oxeas II also differentiate this species from all but *C. sputnika*, which differs by having the smaller oxeas II bearing an enlarged central ring (table 3). This species is notable for its remarkable number of spicule types, including 5 classes of oxeas.

#### Morphology

Sponges are oblong and covered in long spines. Sponge diameter 28 mm x 20-22 mm not including spines, which are up to 6 mm long in one sample and 12 mm long in the other. A double-layer cortex 1-2 mm thick, with subdermal cavities, is obvious in cross section. Samples were brownish-white alive; post-preservation, cortex and spines are white and hard, while the sponge interior is yellowish-white and softer. No oscula or means of attachment are evident. Brooded larvae or direct-developing juveniles, up to 2 mm across, were present within the interior.

#### Skeleton

Skeleton with bundles of oxeas I travelling from a central point and extending into spines. Bundles also contain thick-cladded anatriaenes and protriaenes with their clads near the sponge surface. Bundles in the spines appear to contain oxeas and protriaenes but not anatriaenes. Sponge surface is a thin dermal layer containing only sigmaspires, which are also present in all other parts of the sponge. Supporting this layer is a thick layer of oxeas IIa at various angles from upright to about 45° relative to the surface. A smaller size-class of short oxeas (IIb) are scattered throughout the interior of the sponge. Normal anatriaenes and protriaene clads are not found in the interior, but small-cladded forms of both are found there in low numbers.

One brooded larvae/juvenile was examined and found to contain oxeas IIa and IIb in a surface layer, with interior spicule bundles containing oxeas of indeterminant category. Sigmaspires were seen throughout.

#### Spicules

Shown in figure 13, except when noted below.

**Figure 13.**
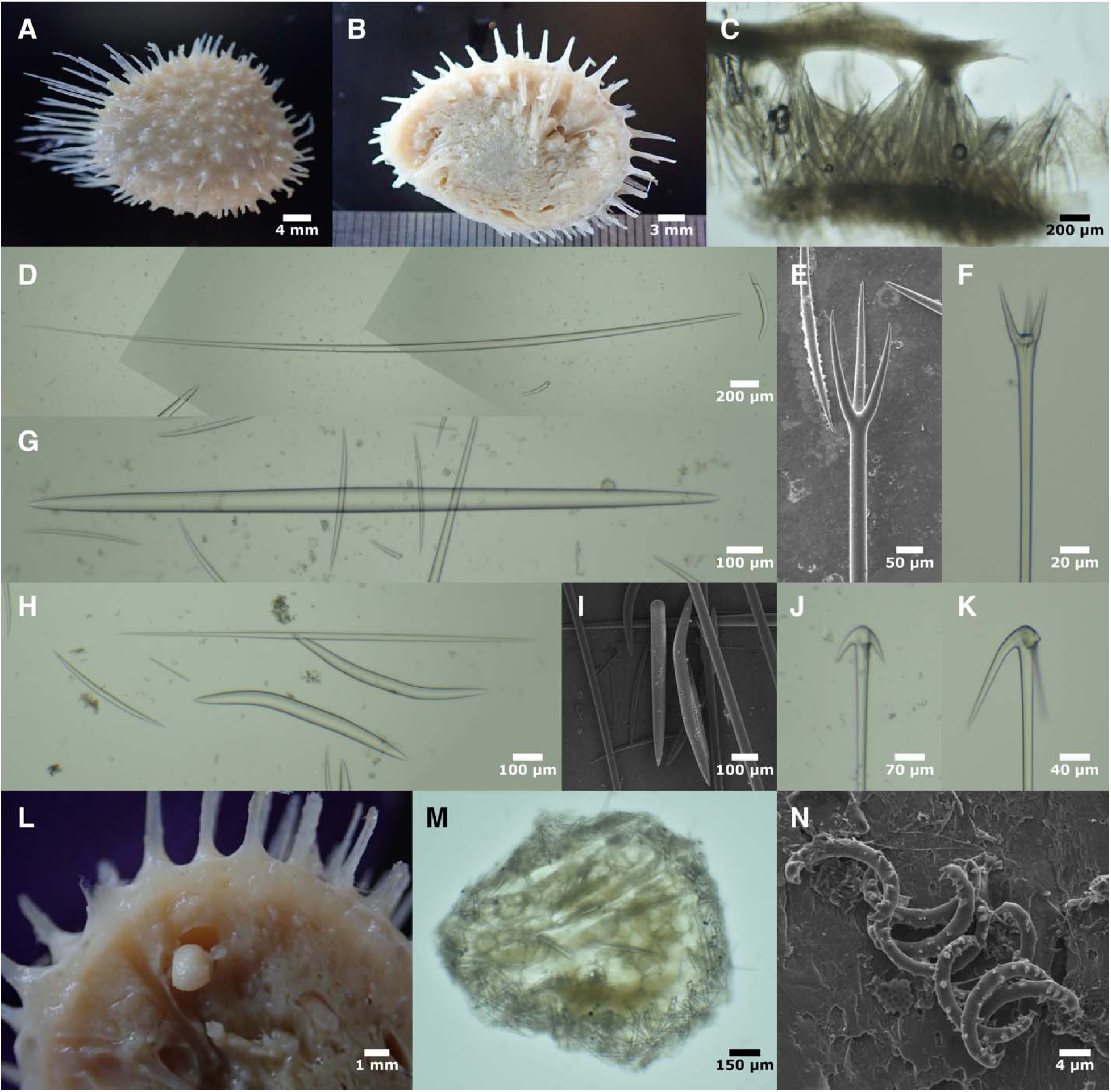
Craniella uniiguni. A: USNM 1478664 post-preservation; B: bisected holotype, post-preservation; C: perpendicular section through cortex of holotype, showing dermal layer (top) supported by oxeas IIa; D: anisoform oxea Ia (composite of three images; a complete oxea IIa also visible, from USNM 1478664); E: protriaene from holotype; F: short-cladded protriaene from USNM 1478664; G: oxea Ib (several complete oxea IIb also visible; from USNM 1478664); H: comparison of oxea III (longest), oxeas IIa (thickest, with bends), and oxea IIb (short and thin), from holotype; I: style and oxea IIa from holotype; J: thick-cladded anatriaene from USNM 1478664; K: thin-cladded anatriaene from holotype (one clad broken); L: closeup of partially dislodged larva/juvenile within bisected holotype; M: the larva/juvenile from L, bisected and digested to show skeleton; N: sigmaspires from holotype.

Oxeas I: the longest size-class of oxeas. Fusiform, thickest in the center and gradually tapering at both ends, but anisoform due to one filiform end. 2971–5743–7783 x 39–63–76 μm (n=26).

Oxeas IIa: stout cortical oxeas, often with one or two pronounced bends. 476–785–1099 x 28– 48–64 μm (n=49).

Oxeas IIb: short interior oxeas; separated from oxeas IIa by their position in the sponge; if all are considered together, length distribution is bimodal. 225–442–658 x 7–19–36 μm (n=71).

Oxeas III: long, thin, symmetrical oxeas; differentiated from oxeas I by being shorter and symmetrical, and from oxeas II and IV by being thinner relative to their length. 586–1416–3240 x 5–14–26 μm (n=44).

Styles: similar to oxeas IIa, but less common. 566–620–731 x 53–58–64 μm (n=7).

Oxeas IV: Fusiform, symmetrical, thickest in the center and gradually tapering at both ends; seen in low numbers in the sponge interior. Differentiated from oxeas I by being shorter, thicker for a given length, and lacking the one filiform end; differentiated from oxeas IIa by being longer and found in the interior; differentiated from oxeas IIb by being much larger; differentiated from oxeas III by being much thicker for a given length. 1669–1922–2164 x 51–64–75 μm (n=7).

Thick-cladded anatriaenes: seen in lower abundance than the other *Craniella* described here, and lack the pointed tops seen in most of those species. Rhabd 2947–4301–6822 μm (n=8) x 21–30– 37 μm (n=25). Clads 79–115–164 x 27–34–41 μm, with length:width ratios 3–3–6 (n=26). Clad:rhabd angle 25°–34°–47° (n=26).

Thin-cladded anatriaenes: only two broken examples seen, so these are possibly foreign. One was found in the choanosome of the paratype, and one was found within the larva/juvenile of the holotype. Rhabd widths 10–13–17 μm. Clads 165–192–219 x 19–22–25 μm, with length:width ratios 7–9–11. Clad:rhabd angle 28°–30°–31° (n=2 for all).

Short-cladded anatriaenes: a few anatriaenes with short clads and wide clad angles were seen in the sponge interior; these were possibly immature versions of other spicules. Rhabd 3200–3293– 3385 μm (n=2) x 7–12–18 μm (n=4). Clads 16–32–56 x 7–11–17 μm, with length:width ratios 2– 3–3 (n=4). Clad:rhabd angle 49°–55°–59° (n=4). Not shown in figure 13.

Protriaenes/prodiaenes: very long clads, slightly curving, of somewhat unequal lengths, only modestly spread away from rhabd axis. Clads had a slight tendency for the tips of the clads to angle inward, but this was only visible under SEM. Most were triaenes (44/46, 96%), but two diaenes seen. Rhabds 3533–5847–10733 μm (n=21) x 17–27–34 μm (n=43). Longest clads 109– 262–355 x 10–18–43 μm, with longest:shortest clad ratios 1.0–1.4–1.8 (n=43). Clad:rhabd angle 9°–16°–25° (n=40).

Short-cladded protriaenes: very short clads, of approximately equal length, with large clad angle. One complete rhabd measured at 3534 μm, widths 8–11–17 μm (n=12). Longest clads 13–43–95 x 2–6–11 μm, with longest:shortest clad ratios 1.0–1.2–1.4 (n=43). Clad:rhabd angle 19°–29°– 47° (n=12).

Sigmaspires: C-shaped, sometimes with a slight twist; scattered large spines throughout, sometimes with larger spines at the ends. Longest possible straight-line measurement across whole spicule 14–15–18 μm (n=40).

#### Distribution and habitat

Known from two samples, collected at different locations, both near the Islands of Four Mountains in the Central Aleutians. Depths 165–186 m.

#### Remarks

The sizes of the sigmaspires, together with protriaene clad length and shape, differentiate this species from all others known from the region. The species is also notable for its many categories of spicules, including 5 types of oxeas, not seen in any other regional species. Of these, the easiest to use as a diagnostic trait are the two types of oxeas II; a similar case is seen only in *C. sputnika*, but in that case the smaller type are centrotylote.

When comparing this species to all *Craniella* known from other locations in the Northern Hemisphere (Lehnert & Stone 2011), all but *C. cranium*, *C. prosperiaradix*, and *C. globosa* can be excluded based on the size of the sigmaspires. *C. cranium* is an Atlantic species, and reports of its presence in the North Pacific are likely erroneous; in any case, it lacks the two categories of oxeas II seen in the new species. *C. prosperiaradix* is excluded based on the presence of additional types of microscleres and *C. globosa* is excluded based on having only one category of oxeas II and smaller clads on protriaenes and anatriaenes.

### *Craniella amlia* sp. nov

#### Material examined

Holotype: USNM 1478667, S. of Amlia Island, Aleutian Islands, (51.93450, -173.71100), 103 m, 30-Jun-2012.

#### Etymology

Named for Amlia Island.

#### Diagnosis

The small sigmaspires differentiate *C. amlia* sp. nov. from all Northeast Pacific species except *C. rocheta* sp. nov. These species are differentiated based on *C. rocheta* sp. nov. having uniquely-shaped thick-cladded anatriaenes and thin-cladded anatriaenes. The spicules of *C. columbiana* sp. nov. are similar to this species, but the sigmaspires and protriaene clads are significantly different in length (table 3).

#### Morphology

The only sample examined is spherical, 43 mm in diameter, and covered in low conules with projecting spicule bundles. An area with several oscula, up to 1.5 mm across, is present on the top of the sponge, and spicule bundles in this area protrude farther than they do in other areas. A double-layer cortex, approximately 1.5 mm thick, is apparent in cross-section.

Post-preservation, all parts of the sponge are light brown. Brooded larvae or direct-developing juveniles, up to 2 mm across, were present in the interior.

#### Skeleton

Radial skeleton, with bundles of oxeas I, anatriaenes, and protriaenes spiraling from a central point and extending into conules. Sponge surface is a thin dermal layer containing only sigmaspires, which are also present in other parts of the sponge. Supporting this layer is a thick layer of oxeas II at various angles from upright to about 45° relative to the surface.

#### Spicules

Shown in figure 14, except when noted below.

**Figure 14.**
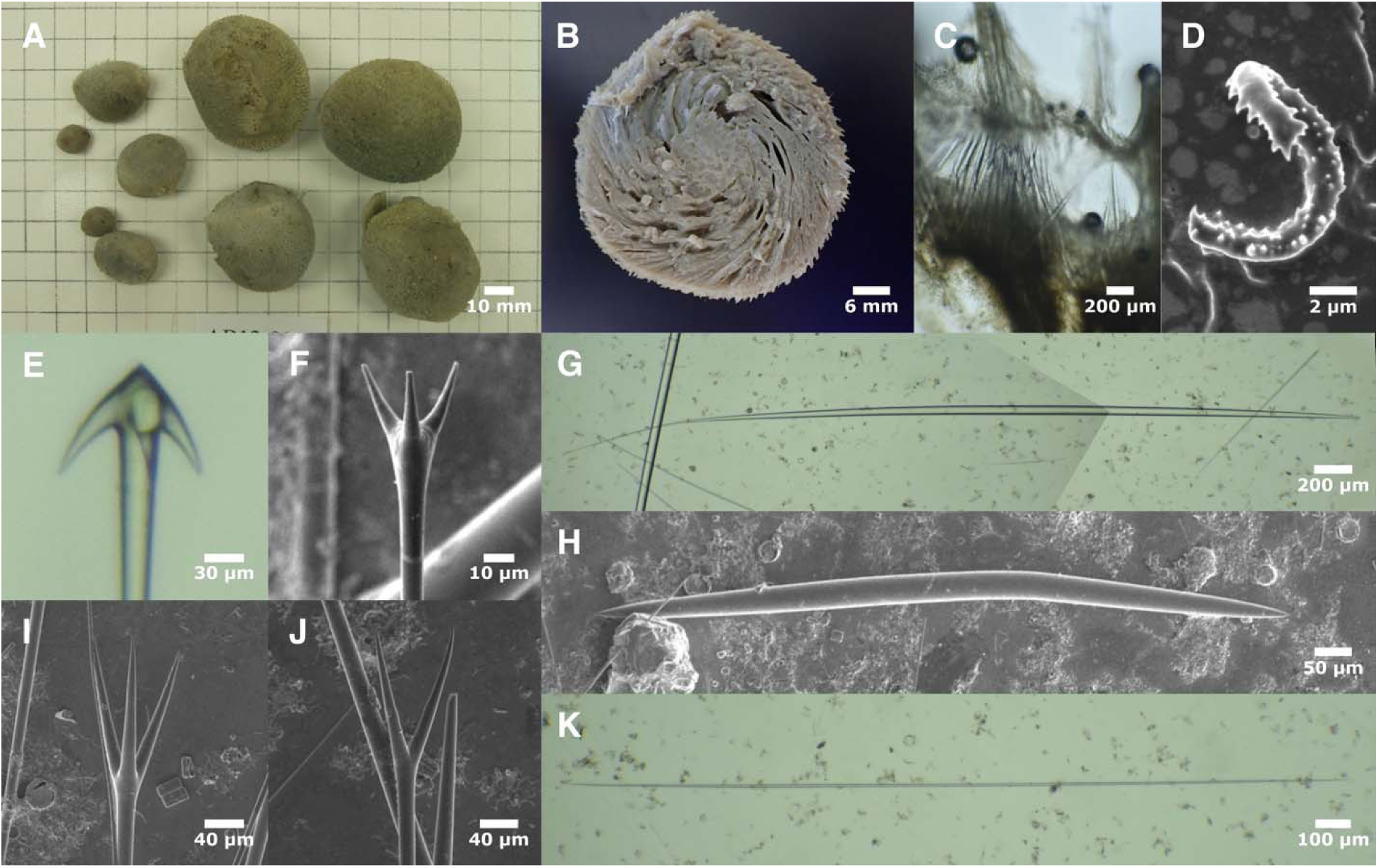
Craniella amlia. A: Freshly collected samples (photo curtesy of Robert Stone; top center individual appears to be the sample vouchered as USNM 1478667 examined here, which is now the holotype); B: bisected sample with juveniles visible, post-preservation; C: perpendicular section through cortex; D: sigmaspire; E: anatriaene; F: short-cladded protriaene; G: oxea I (composite of two images); H: oxea II; I: protriaene; J: prodiaene; K: oxea III. All photos from holotype.

Oxeas I: the longest size-class of oxeas. Fusiform, thickest in the center and gradually tapering at both ends, but anisoform due to one filiform end. 3400–4033–4643 x 26–43–49 μm (n=14).

Oxeas II: stout cortical oxeas. 580–982–1256 x 20–37–50 μm (n=32).

Oxeas III: long, thin, symmetrical oxeas; differentiated from oxeas I by being shorter and symmetrical, and from oxeas II by being thinner relative to their length. 460–1220–2268 x 5–10– 18 μm (n=15).

Thick-cladded anatriaenes: usually with pointed tops. Rhabd 3472–5193–7168 μm (n=5) x 20– 25–29 μm (n=29). Clads 62–101–124 x 17–30–40 μm, with length:width ratios 3–3–4 (n=29). Clad:rhabd angle 28°–34°–42° (n=9).

Protriaenes/prodiaenes: clads are straight, of equal lengths, and spread away from rhabd axis. Most were triaenes (22/25, 88%). Rhabds 3338–5366–6345 μm (n=5) x 12–18–24 μm (n=24). Longest clads 93–128–159 x 8–14–20 μm, with longest:shortest clad ratios 1.0–1.1–1.2 (n=24). Clad:rhabd angle 17°–21°–27° (n=8).

Short-cladded protriaenes: very short clads, of approximately equal length, with wide clad angle. Rhabds 3489–2701–2921 μm (n=3) x 7–12–19 μm (n=7). Longest clads 15–37–62 x 3–9–13 μm.

Sigmaspires: C, S, or spiral-screw shaped, with modest spines throughout and larger ones at the ends. Longest possible straight-line measurement across whole spicule 6–8–11 μm (n=54).

#### Distribution and habitat

The only examined sample was trawled near the Central Aleutian Island of Amlia at 103 m depth. Field photos indicate that multiple sponges matching the morphology of the examined sampled were trawled at this location (figure 14A, provided by Robert Stone, NOAA).

#### Remarks

For this species, DNA sequencing was successful at two regions of the cox1 locus, which place it in the *C. zetlandica* clade (figure S9; this is not shown in figure 1 because sequencing was unsuccessful at the 28S locus). These cox1 regions were insufficient to genetically differentiate this species from *C. uniiguni* sp. nov., but the spicules and morphology of these species are very different, so I am confident they are not conspecific. The spicules of this sponge are more similar to species in the *C. wolfi* clade, but at least one spicule measurement differentiates *C. amlia* sp. nov. species from each of these distantly related species (table 3).

Most *Craniella* species known from the Northern Hemisphere are differentiated from this new species by sigmaspire length (Lehnert & Stone 2011). Exceptions are two Japanese species, *C. ovata* and *C. lentisimilis*. There are other differences between the new species and *C. lenisimilis*, including smaller clads on the anatriaenes, no short-cladded protriaenes, and no oxeas III; this species is also described as having a dense mat of tangential oxeas in the cortex, and the oxeas I are not anisoform. Differences between the new species and *C. ovata* include a broad equatorial stripe of papillae, smooth upper half, root of spicules, a lack of oxeas III, and a lack of small-cladded protriaenes in *C. ovata*. This species is also an unlikely match due to differences in habitat.

Field notes shared by Robert Stone (NOAA) indicate that 9 samples matching the examined voucher were collected at the same location; I am unsure where the others are vouchered, but some may be available at the Zoologische Staatssammlung München. These samples may be identified as *C. spinosa*, which was the previous identification of the sample examined here.

### *Craniella shemya* sp. nov

#### Material examined

Holotype: USNM 1478666, S. of Shemya Island, Near Islands, (52.44320, 174.30300), 228 m, 27-Jun-2012.

#### Etymology

Named for Shemya Island.

#### Diagnosis

Sigmaspire length easily differentiates this species from all others in the Northeast Pacific except *C. vermisigma* sp. nov. and *C. uniiguni* sp. nov. Protriaene and anatriaene clad lengths are much longer in *C. vermisigma* sp. nov., which also has different sigmaspire morphology. Many characters are different in *C. uniiguni* sp. nov. including protriaene clad length and shape and two length classes of oxeas II (table 3).

#### Morphology

The only known sample is spherical, about 8 mm in diameter, and covered in conules with projecting spicule bundles. No oscula are apparent, but a small spicule mat is present that may have attached the sponge to a substrate. Despite the small size of the sample, a double-layer cortex is obvious in cross-section, approximately 1.5 mm thick on the upper parts of the sponge, but thinner and less clear on the bottom.

#### Skeleton

Spicule bundles radiate from the center of the sponge to the periphery. Due to the small size of the sample, and the lack of significant variation in skeletal anatomy of the other *Craniella* describe here, tissue sectioning was limited, but a dense layer of chaotically arranged, mostly upright oxeas II is present in the cortex.

#### Spicules

Shown in figure 15, except when noted below.

**Figure 15.**
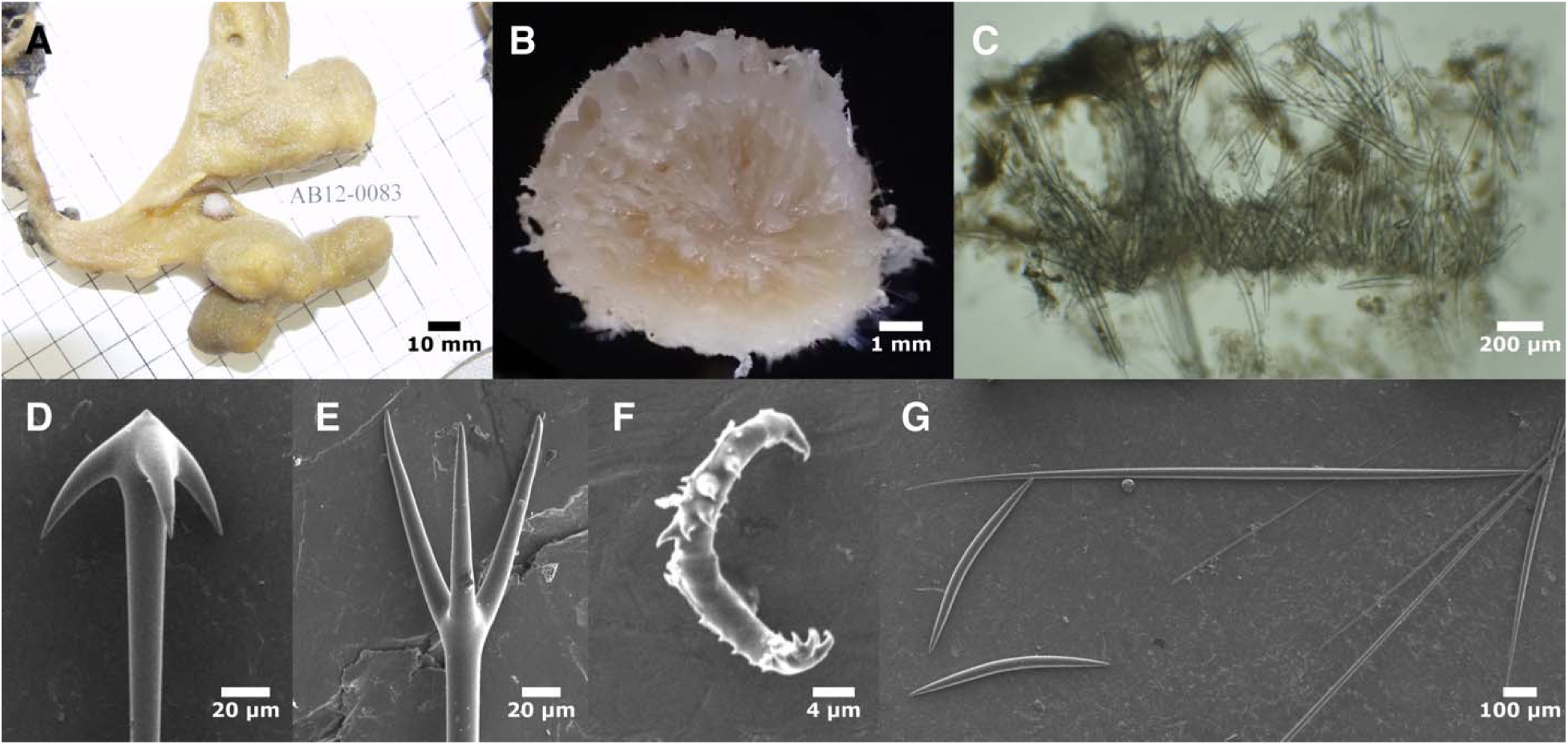
Craniella shemya. A: Freshly collected sample attached to another sponge (photo curtesy of Robert Stone); B: bisected sample post-preservation; C: perpendicular section through cortex; D: anatriaene; E: protriaene; F: sigmaspire; G: oxea I with two oxeas II. All photos from holotype.

Oxeas I: the longest size-class of oxeas. Fusiform, thickest in the center and gradually tapering at both ends. Often slightly anisoform due to one filiform end, but less consistently anisofom than most other *Craniella* described here, with many nearly symmetrical. 1125–1950–2307 x 8–29– 39 μm (n=20).

Oxeas II: stout cortical oxeas. 489–619–718 x 18–28–36 μm (n=23).

Thick-cladded anatriaenes: usually with pointed tops. Rhabd 1176–4157–7459 μm (n=3) x 11– 18–27 μm (n=19). Clads 40–77–124 x 12–22–42 μm, with length:width ratios 3–4–5 (n=19). Clad:rhabd angle 30°–38°–50° (n=16).

Protriaenes/prodiaenes/protetraenes: clads are straight, of equal lengths, and spread away from rhabd axis. Only 64% (7/11) were triaenes, with others evenly split between prodiaenes and protetraenes. One rhabd measured at 2725 μm long; widths 15–16–18 μm (n=10). Longest clads 69–106–132 x 9–12–17 μm, with longest:shortest clad ratios 1.0–1.1–1.2 (n=10). Clad:rhabd angle 11°–25°–54° (n=11).

Short-cladded protriaenes: very short clads, of approximately equal length, with wide clad angle. Rhabd widths 14–15–17 μm. Longest clads 38–46–54 x 9–10–11 μm (n=2). Not shown in figure 15; see figures 14F & 19C for similar examples.

Sigmaspires: C, S, or spiral-screw shaped, with substantial spines throughout; some with accentuated spines at the ends. Longest possible straight-line measurement across whole spicule 13–16–19 μm (n=51).

#### Distribution and habitat

The only known sample was collected in the Near Islands, South the island of Shemya, 228 m depth. Robert Stone (NOAA; personal communication) reports that this sample was found attached to *Auletta krautteri* Austin *et al*. 2014.

#### Remarks

Though only one small sample is available from this species, it is quite genetically distinct from all others with data available. This is especially true at cox1, where the species is an outgroup to the other Pacific species in the *C. zetlandica* clade, with roughly 2% sequence divergence to all of them. Moreover, the large sigmaspires make this species morphologically distinct from all others except *C. vermisigma* sp. nov. and *C. uniiguni* sp. nov., which are morphologically distinct in multiple other ways (table 3).

The new species can also be distinguished from all other *Craniella* known from the Northern Hemisphere using the table compiled by Lehnert and Stone (2011). All but *C. cranium*, *C. prosperiaradix*, and *C. globosa* can be excluded based on the size of the sigmaspires. *C. cranium* is an Atlantic species, and reports of its presence in the North Pacific are likely incorrect; this species was not considered further. *C. prosperiaradix* is excluded based on the presence of additional types of microscleres and *C. globosa* is excluded based on anatriaenes with much shorter clads, much shorter oxeas II (despite the small size of *C. shemya*), and habitat.

Note that the original voucher label incorrectly reported this sample as having been collected at a decimal longitude of -174.303, rather than 174.303 (west of the dateline, when in fact it was east of the dateline). The correction was made by referring to the original data records, as communicated by Robert Stone (NOAA).

### *Craniella hamatum* (Koltun, 1966)

#### Material examined

USNM 1478661, Pribilof Canyon, Bering Sea, (56.04840, -168.37900), 225 m, 21-Jun-2016.

#### Diagnosis

The abundant harpoon-shaped anamonaenes are diagnostic.

#### Morphology

The only sample examined was oblong, 38 mm high and 25 mm wide, and covered in large (up to 2.5 mm) conules with protruding spicule bundles. An oscular area at the apex appears to have about 20 small oscula, while the base has a spicular mat. A double layer cortex is obvious in cross-section; it is about 3.5 mm thick on one side, but only 1.0 mm on the other. Beige post-preservation. Samples described from the Kuril Islands are similar in shape and size (30 x 20 mm).

#### Skeleton

Skeleton with bundles of oxeas I and harpoon-like anamonaenes travelling from a central point and extending into conules, with protriaenes piercing the surface. Cortex sparsely populated with cortical oxeas II, which are at varying angles, from upright to nearly tangential. Anatriaenes seen only in the basal spicule mat, where protriaenes were also present. Sigmaspires abundant throughout.

#### Spicules

Shown in figure 16, except when noted below.

**Figure 16.**
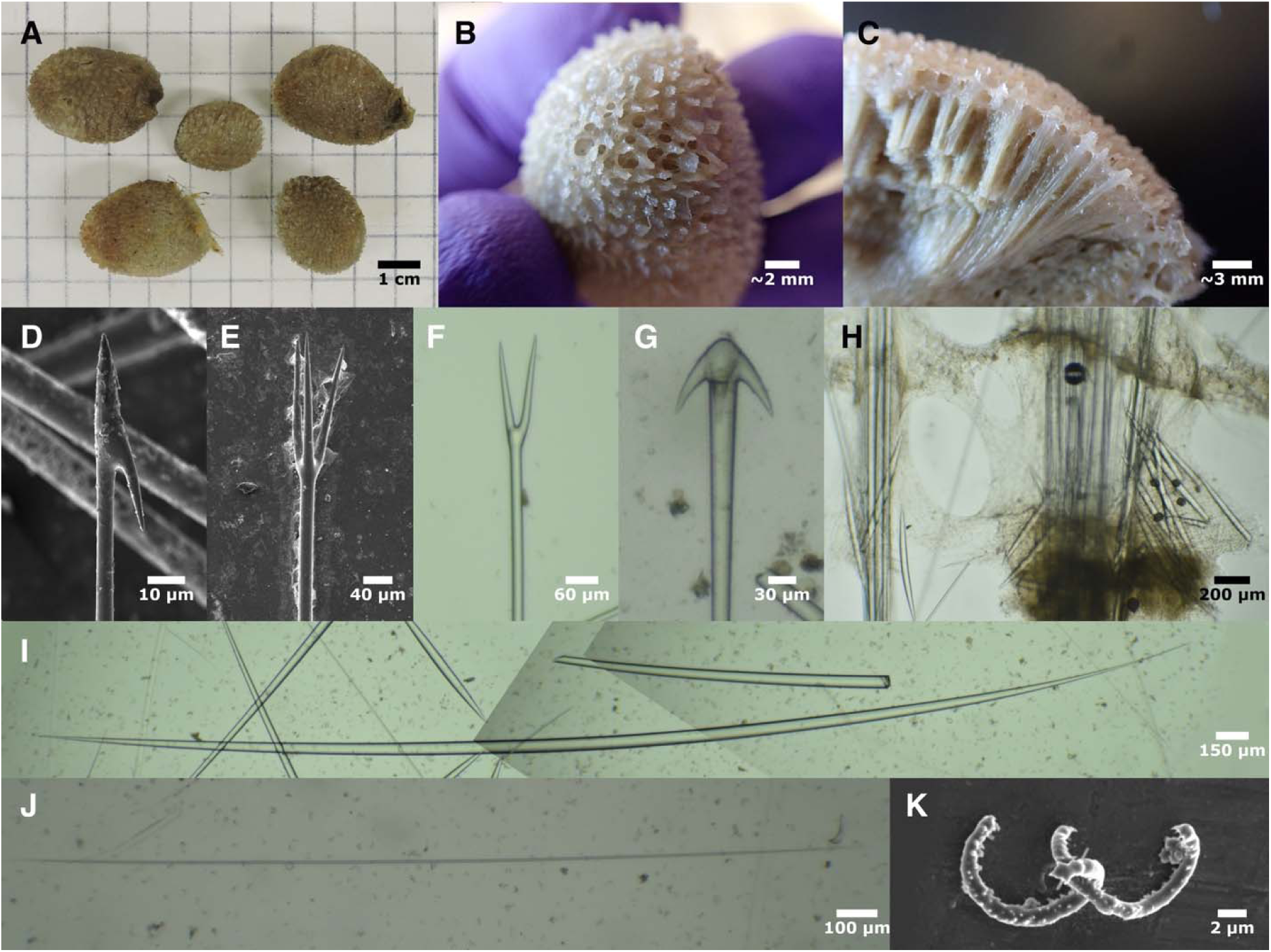
Craniella hamatum. A: Freshly collected whole individuals; sample on the upper right appears to be USNM 147866, the voucher examined (photo curtesy of Robert Stone) ; B: apical oscula on preserved sample; C: preserved, bisected sample; D: harpoon-shaped anamonaene; E: protriaene; F: prodiaene; G; anatriaene; H: perpendicular section through cortex showing scattered, mostly upright oxeas II and spicule bundles; I: oxea I (composite of three images); J: oxea III; K: sigmaspires. B-K from USNM 1478661.

Oxeas I: the longest size-class of oxeas. Fusiform, thickest in the center and gradually tapering at both ends. Often slightly anisoform, but less consistently anisoform than most other *Craniella* described here. 3418–5139–6472 x 39–53–66 μm (n=15). Type material previously described as “variously ended”, 3350–4700 x 32–58 μm.

Oxeas II: stout cortical oxeas. 337–760–1206 x 21–40–50 μm (n=28). Type material previously described as 600–940 x 29–46 μm.

Oxeas III: long thin oxeas, thinner than oxeas I or II for a given length. 869–1796–3587 x 6–7– 14 μm (n=12). Not described from type material.

Harpoon-shaped anamonaenes: with a pointed top and a single small clad. Rhabd 2792–5156– 6380 μm (n=9) x 4–7–12 μm (n=19). Clads 51–57–66 x 3–4–6 μm. Type material previously described with rhabds 2680–5700 x 9–11 μm.

Thick-cladded anatriaenes: usually with pointed tops. One rhabd measured at 6513 μm long; widths 20–26–30 μm (n=27). Clads 84–98–116 x 24–30–40 μm, with length:width ratios 3–3–4 (n=27). Clad:rhabd angle 27°–35°–41° (n=26). Type material previously described with rhabds 5000 x 21 μm.

Short-cladded anatriaenes: not seen in the sample examined, but described from type material. Rhabd 1000–1500 x 5–7 μm, clads 20–40 μm long.

Protriaenes/prodiaenes/protetraenes: clads are straight, of equal lengths, and only slightly spread away from rhabd axis. Most (89%, 40/45) were triaenes, with others diaenes except for one tetraene. Rhabds 4350–4906–5186 (n=5) x 13–18–24 μm (n=45). Longest clads 109–174–209 x 9–14–18 μm, with longest:shortest clad ratios 1.0–1.1–1.3 (n=28). Clad:rhabd angle 8°–14°–22° (n=29). Type material previously described with rhabds 1300–3400 x 8–16 μm, clads 60–140 μm.

Sigmaspires: C, S, or spiral-screw shaped, spines throughout, often with the largest spines at the ends. Longest possible straight-line measurement across whole spicule 8–11–13 μm (n=65) x 1.1–1.4–1.6 μm (n=13). Type material previously described as 8–12 μm.

#### Distribution and habitat

The sample examined was trawled from 225 m depth at Pribilof Canyon, near St. George Island in the Bering Sea. Previously known from the Pacific Coast of the Southern Kuril Islands, 414 m.

#### Remarks

This species is easily identified from its harpoon-shaped monanenes, and other spicule measurements were a good match to the previously described Russian material. Koltun (1966) and I both found the cortical oxeas to be of a low density, which is in contrast to all other *Craniella* examined here except for *C. rocheta* sp. nov., where they are even sparser. One notable discrepancy between the type description and the sponge examined here is that the oxeas II were previously described as tangential, while the Aleutian sample had mostly upright oxeas.

I successfully sequenced two small regions of the cox1 locus, which were identical in this sample and *C. columbiana* sp. nov. I also obtained 28S data from these species, and surprisingly, these sequences were also identical (though most of this sequence was from the less variable, D3D5 region). Despite this, I think it is unlikely that these samples are conspecific. I searched extensively for harpoon-shaped spicules in *C. columbiana* sp. nov. and found none, and in contrast to the monanenes that were present only in small sponge-rooting *Tetilla*, the *C. hamatum* sample was the largest of these *Craniella* examined. *C. hamatum* also differed from these other species in the shapes and sizes of the anatriaenes, and in the angles of the protriaenes (table 3).

Notes shared by Robert Stone (NOAA) indicate that 5 samples were trawled at the same time as the one I examined, and that the largest was vouchered at Zoologische Staatssammlung München.

### *Craniella columbiana* sp. nov

#### Material examined

Holotype: RBC 980-00259-006, Dunsany Passage, British Columbia, (50.90167, -126.83500), 20-100 m, 23-Mar-1980. Paratypes: RBC A-094-00006, Swiftsure Bank, British Columbia, (48.55667, -124.98333), 55 m, 27-Jan-1965; RBC 975-00072-024, Owen Point, British Columbia, (48.47167, -124.44500), 200 m, 28-Jan-1975.

#### Diagnosis

The sigmaspires of this species are similar in size to two other Northeast Pacific species: *C. spinosa* and *C. hamatum*. The much larger, uniquely-shaped protriaenes in *C. spinosa* serve to differentiate that species, and *C. hamatum* has harpoon-shaped monanenes. The sigmaspires of *C. amlia* sp. nov. are only slightly smaller than this species, but *C. amlia* sp. nov. is further differentiated by its protriaene clad lengths. Likewise, the sigmaspires of *C. rocheta* sp. nov. are only slightly smaller than *C. columbiana* sp. nov., but that species is differentiated by its unique anatriaene shape and the presence of thin-cladded anatriaenes (table 3).

#### Etymology

Named for British Columbia.

#### Morphology

Holotype consists of two spherical beige sponges, 23-30 mm in diameter, covered in conules with protruding spicule bundles. No oscula or basal spicule mat is evident, but the sponges are covered in considerable obscuring debris. A double-layer cortex is obvious in cross-section, about 3 mm thick. One of the paratypes is a smaller (10 mm) spherical sample, but the other (RBC A-094-00006) is mealy a small fragment.

#### Skeleton

Skeleton radial, with bundles of spicules radiating from a central point that is closer to the bottom of the sponge than the top. The bottom of the sponge, below the gathering point of the bundles, is less firm than the top, but also cartilaginous and hard to cut through. Spicule bundles contain oxeas I, anatriaenes, and protriaenes, the latter two types have their clads near the sponge surface or outside it. Cortex provisioned with oxeas II, varying from upright to about 45° angles to the surface. Sigmaspires occur throughout the sponge.

#### Spicules

Shown in figure 17, except when noted below. In contrast to species like *C. uniiguni* sp. nov., *C. rocheta* sp. nov., *T. vancouverensis* sp. nov., and *T. villosa* sp. nov., the thin- and thick-cladded anatriaenes were difficult to separate into classes. To increase comparability with these other species, they were separated based on having clad:rhabd ratios greater than or less than 4.5, but further work is needed to determine if there are truly two separate types.

**Figure 17.**
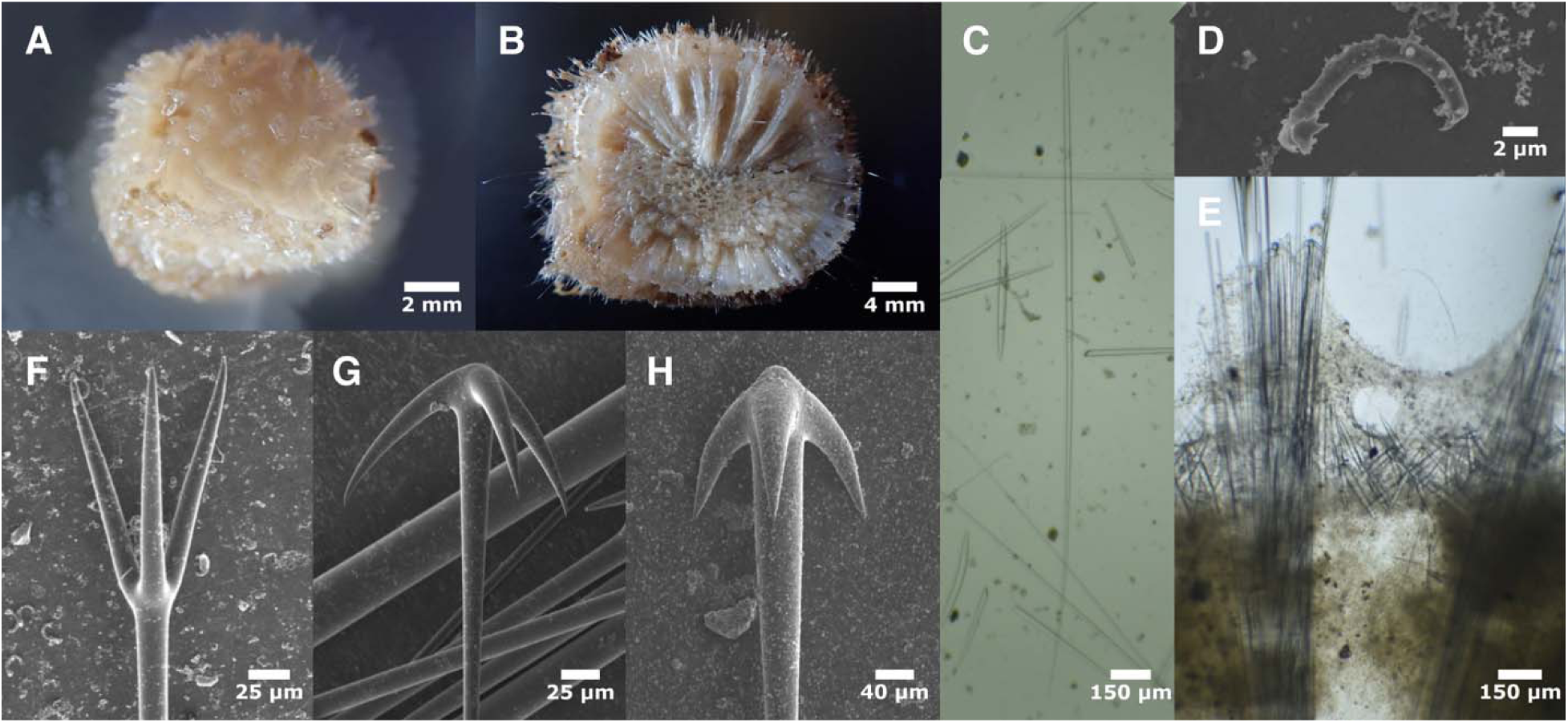
Craniella columbiana. A: Sample RBC 975-00072-024 post-preservation; B: holotype, bisected, post-preservation; C: anisoform oxea I and small oxeas II, from RBC 975-00072-024; D: sigmaspire from holotype; E: perpendicular section through cortex from RBC 975-00072-024; F: protriaene from holotype; G: thin-cladded anatriaene from RBC 975-00072-024; H: thick-cladded anatriaene from holotype.

Oxeas I: the longest size-class of oxeas. Fusiform, thickest in the center and gradually tapering at both ends; most are anisoform due to one filiform end. 1670–2857–4390 x 17–37–57 μm (n=19).

Oxeas II: stout cortical oxeas. 363–610–927 x 18–35–53 μm (n=60).

Oxeas III: long, thin, symmetrical oxeas; differentiated from oxeas I & II by being thinner relative to their length, and lacking the asymmetry of oxeas I. 1023–1627–2128 x 7–12–17 μm (n=7). Not shown in figure 17; for similar examples, see figures 14K & 16J.

Thick-cladded anatriaenes/anadiaenes: some have a pointed top, but this was frequently absent. Hard to separate from thin-cladded anatriaenes in some samples; clad:rhabd ratios < 4.5 were used as a criterion. One complete rhabd measured at 2344 μm long; widths 14–31–45 μm (n=42). Clads 61–135–204 x 17–35–57 μm, with length:width ratios 3–4–4 (n=42). Clad:rhabd angle 28°–37°–53° (n=17).

Thin-cladded anatriaenes: hard to separate from thin-cladded anatriaenes in some samples; clad:rhabd ratios > 4.5 were used as a criterion. Rhabd length not measured; widths 9–20–41 μm (n=73). Clads 75–131–232 x 12–23–52 μm, with length:width ratios 5–6–8 (n=73). Clad:rhabd angle 27°–40°–50° (n=72).

Protriaenes/prodiaenes: clads are of equal lengths, some are straight and some curve out slightly. Curved clads are reminiscent of *C. spinosa*, but are much shorter, less dramatically curved, and have a greater angle with the rhabd. The tips of the clads point inward slightly, but this is only visible under SEM. Most (88%, 50/57) were triaenes, with the others prodiaenes. Rhabds 2143– 3103–4024 μm (n=7) x 10–18–37 μm (n=86). Longest clads 78–151–219 x 6–13–30 μm, with longest:shortest clad ratios 1.0–1.1–1.2 (n=76). Clad:rhabd angle 14°–21°–34° (n=74).

Sigmaspires: S, C, or spiral-screw shaped, spiny throughout, often with larger spines at ends. Longest possible straight-line measurement across whole spicule 7–11–14 (n=80) x 1.3–1.4–1.6 μm (n=10).

A few spherical spicules were also seen, roughly 100 μm in diameter (not shown).

#### Distribution and habitat

Known from British Columbia, both North and South of Vancouver Island, 55–200 m.

#### Remarks

Eight of the 10 *Craniella* described in this report have a unique spicular trait that makes diagnosis straightforward, but *C. columbiana* sp. nov. is not one of those species. It can be differentiated from several similar species by measurement of the sigmaspires, anatriaenes, and protriaenes, as detailed in the diagnosis and table 3. The most similar species is the sympatric *C. spinosa*, and the lack of DNA data from that species raises the possibility that *C. columbiana* sp. nov. is conspecific. I could not find thick-cladded anatriaenes were not found in *C. spinosa*, but I was only able to examine a small piece of a syntype, they are sometimes found only in some regions of the sponge. Support this, only thin-cladded anatriaenes were mentioned in the original description of *C. spinosa*; in the same report, *T. villosa* anatriaenes are noted as occurring in both thin and thick-cladded forms (Lambe, 1893). Additionally, the protriaenes were conspicuously different in both size and shape in *C. spinosa*, and this is similar to the differences seen between species for which we have DNA data. The thin-cladded anatriaenes were also subtly, but significantly, different, adding additional evidence for species-level differentiation (table 3). We also note that *C. amlia sp. nov.* differs from *C. columbiana* sp. nov. mainly in small (but significant) differences in sigmaspire size; these species are also well-differentiated at the cox1 locus. Finally, as explained in the *C. hamatum* remarks above, I was unable to genetically separate *C. columbiana* sp. nov. and *C. hamatum* with the limited data obtained. Despite this, I think it is unlikely that these samples are conspecific based on their substantial morphological differences.

When comparing all *Craniella* previously known from the Northern Hemisphere to *C. columbiana* sp. nov., all but two Japanese species, *C. ovata* and *C. lentisimilis* can be excluded based on the size of the sigmaspires (Lehnert & Stone 2011). There are other differences between the new species and *C. lenisimilis*, including smaller clads on the anatriaenes and protriaenes; this species is also described as having a dense mat of tangential oxeas in cortex, and the oxeas I are not anisoform. Differences between the new species and *C. ovata* include a broad equatorial stripe of papillae, a root of spicules, and no oxeas III. This species is also an unlikely match due to differences in habitat.

### *Craniella rocheta* sp. nov

#### Material examined

Holotype: USNM 1517729, South of Cape Kaletka, Aleutian Islands, (53.98970, -166.35400), 1-15 m, 7-Sep-2018.

#### Diagnosis

The shape of the thick-cladded anatriaenes is unique among *Craniella* in the region, with pointy tops and clads that have very low angles. Additional diagnostic characters include having few cortical oxeas, having both thick and thin-cladded anatriaenes (shared with only *C. uniiguni* sp. nov. and *C. spinosa*, which both have much longer protriaene clads, among other differences) and having small sigmaspires (shared with only *C. amlia* sp. nov., though close to several others; table 3).

#### Etymology

Named for the uniquely-shaped thick-cladded anatriaenes, which resemble rocket ships.

#### Morphology

Holotype is a white, relatively large (40 x 30 mm) ovoid sponge covered in conules with long protruding spicule bundles. A cortex (1.0-1.5 mm thick) is obvious in cross-section, though it isn’t clearly double-layer due to a lack of sub-cortical spaces visible to the naked eye (though these spaces are visible in tissue sections). Oscules not evident; basal mat of spicules present.

#### Skeleton

Skeleton radial, with bundles of spicules radiating from a central point to the top and sides of the sponge. The bottom half of the sponge is spongy, cartilaginous, and less radial. Spicule bundles contain oxeas I, thin-cladded anatriaenes, and protriaenes. Anatriaene clads are near the surface or slightly protruding, while protriaenes are mostly protruding. Cortex consists of a thin dermal layer containing debris and sigmaspires and a thicker sub-dermal layer. The upright palisade of oxeas II seen in the cortex of other *Craniella* appears vestigial, with only a few upright oxeas scattered through the cortex. Thick-cladded anatriaenes only seen in bottom regions of the sponge and in basal spicule mat; thin-cladded anatriaenes found throughout. Sigmaspires abundant in all regions of the sponge.

#### Spicules

Shown in figure 18, except when noted below.

**Figure 18.**
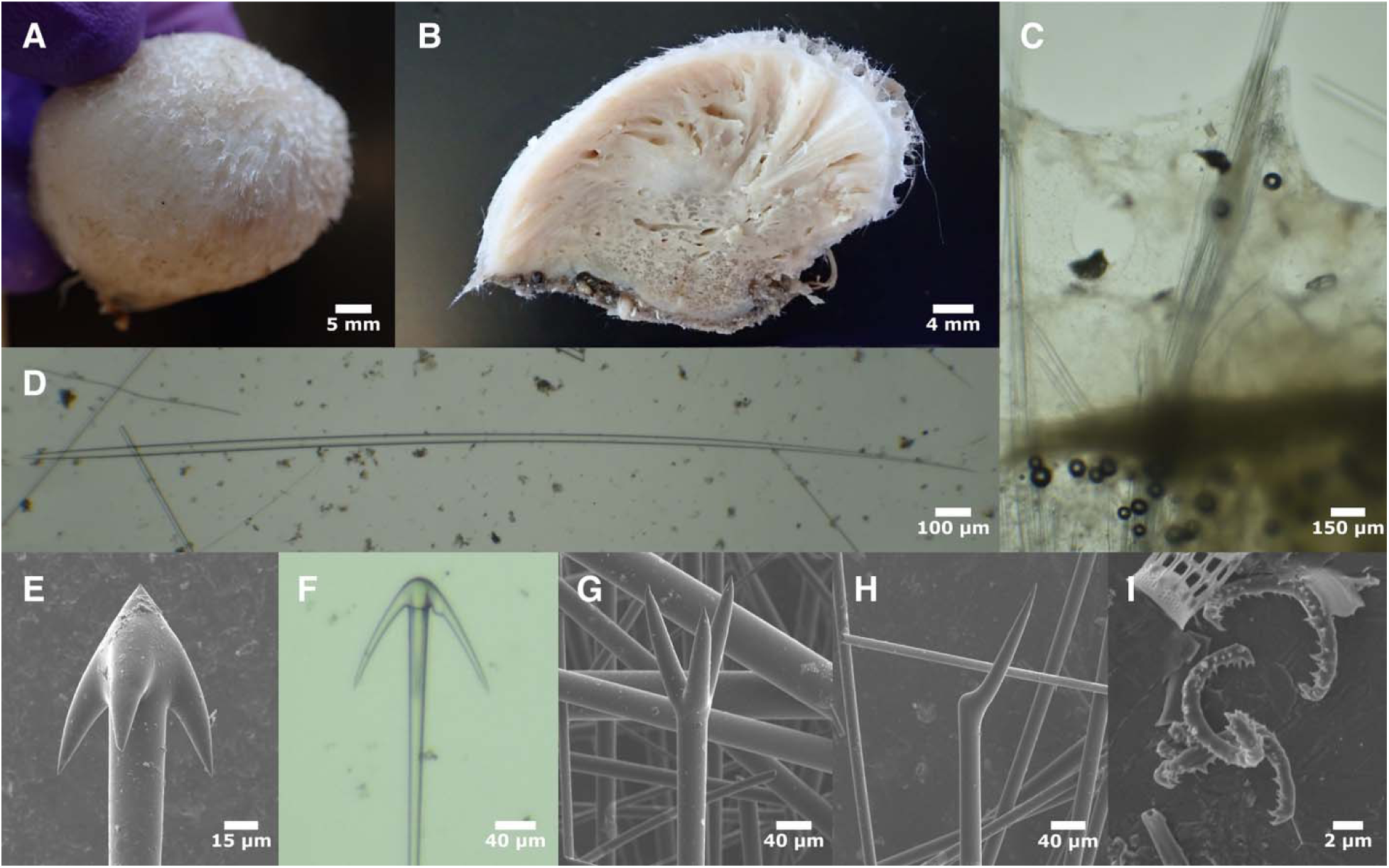
Craniella rocheta. A: Whole individual, post-preservation; B: bisected sample; C: perpendicular section through cortex, note that there are only a few scattered oxeas II; D: oxea I; E: thick-cladded anatriaene with “rocket ship” shape; F: thin-cladded anatriaene; G: protriaene; F: promonaene; G: sigmaspires. All photos from USNM 1517729.

Oxeas I: the longest size-class of oxeas. Fusiform, thickest in the center and gradually tapering at both ends; some are anisoform due to one end being more filiform, but many are symmetrical. 2710–4308–6657 x 23–34–56 μm (n=18).

Oxeas II: stout cortical oxeas; not abundant. 330–578–736 x 10–25–39 μm (n=23). Partially visible in figure 18C; see figure 19D for similar example.

Oxeas III: long, thin, symmetrical oxeas; differentiated from oxeas I and II by being thinner relative to their length. 686–1851–4565 x 6–11–22 μm (n=23). Not shown in figure 18; for similar examples, see figures 14K & 16J.

Thin-cladded anatriaenes: One rhabd measured at 7153 μm long, widths 6–14–22 μm (n=24). Clads 122–151–197 x 14–20–28 μm, with length:width ratios of 4–8–10 (n=24). Clad:rhabd angle 21°–29°–35° (n=24). One was seen with a forked clad.

Thick-cladded anatriaenes: One rhabd measured at 6548 μm long, widths 19–25–30 μm (n=16). Clads 71–93–112 x 24–31–37 μm, with length:width ratios of 3–3–4 (n=16). Clad:rhabd angle 18°–26°–35° (n=16).

Short-cladded anatriaenes: uncommon, small anatriaenes with very small clads and a large angle; may be immature form of other anatriaenes. One rhabd measured at 3299 μm long, widths 6–8– 10 μm (n=6). Clads 21–46–80 x 6–10–14 μm, with length:width ratios of 3–4–6 (n=6). Clad:rhabd angle 31°–41°–51° (n=16). Not shown in figure 18.

Protriaenes/prodiaenes/promonaenes: clads are straight, of equal lengths, and modestly spread away from rhabd axis. Most (81%, 26/32) were triaenes, but prodiaenes relatively common and one promonanene seen. Rhabd lengths not measured; width 14–19–24 μm (n=26). Longest clads were 109–154–188 x 10–13–17 μm, with longest:shortest clad ratios of 1.0–1.1–1.3 (n=26). Clad:rhabd angle 12°–18°–29° (n=25). One was seen with a forked clad.

Short-cladded protriaenes: shaped like long-cladded protriaenes, but with a greater angle. one rhabd measured at 2696 μm long; width 6–11–14 μm (n=8). Longest clads were 29–67–91 x 3– 7–9 μm, with longest:shortest clad ratios of 1.0–1.1–1.3 (n=8). Clad:rhabd angle 17°–27°–38° (n=8). Not shown in figure 18; see figures 14F & 19C for similar examples.

Sigmaspires: Most are roughly C-shaped, with a slight spiraling, spiny throughout, often with larger spines at ends. Longest possible straight-line measurement across whole spicule 7–9–11 (n=34).

#### Distribution and habitat

The only known sample was collected by divers, in less than 15 m of water, Unalaska Island, Alaska. Of the Aleutian *Craniella* examined here, this is the only one known from shallow water.

#### Remarks

The “rocket ship” shaped anatriaenes in the basal regions of this sponge are diagnostic, and differentiate it from all others in the region. It is also unique in having very few cortical oxeas. Even in incomplete samples, where these traits could be difficult to assess, the size of the sigmaspires and shape of the thin-cladded anatriaenes and protriaenes serve to differentiate this species from others in the region (table 3). At the genetic level, this species is closely related to *C. hamatum* and *C. columbiana* sp. nov, but 1-2 SNPs differentiate it from these species at both loci.

Of the *Craniella* previously known from the Northern Hemisphere, two Japanese species have similar sigmaspires: *C. ovata* and *C. lentisimilis*. Neither of these species have long, thin-cladded anatriaenes.

### *Craniella spinosa* Lambe, 1893

#### Material examined

Syntype: CMNI 1900-2814, Elk Bay, Discovery Passage, British Columbia, 36-45 m, 23-Jul-1885.

#### Diagnosis

The shape of the protriaenes is unique among *Craniella* in the region, with long clads that are parallel with the rhabd at first, then diverge mid-clad. Clads are also longer than all sympatric species save *C. sputnika* (which lacks sigmaspires) and *C. uniiguni* sp. nov. (which has larger sigmaspires and two classes of oxea II).

#### Morphology

I examined only a small fragment of a syntype, but the original description states that this species is ovoid, 1.8–19.0 mm in greatest diameter, and grayish-yellow post-preservation. It has up to three apical oscula 1-2 mm in diameter. Bundles of spicules protrude 5 mm from conules covering the sponge.

#### Skeleton

Skeleton radial, with bundles of oxeas, protriaenes, and anatriaenes radiating from a central point and protruding several millimeters from the sponge surface. Cortex consists of a thin dermal layer containing debris and sigmaspires and a thicker sub-dermal layer with a dense palisade of upright cortical oxeas.

#### Spicules

Shown in figure 19, except when noted below.

**Figure 19.**
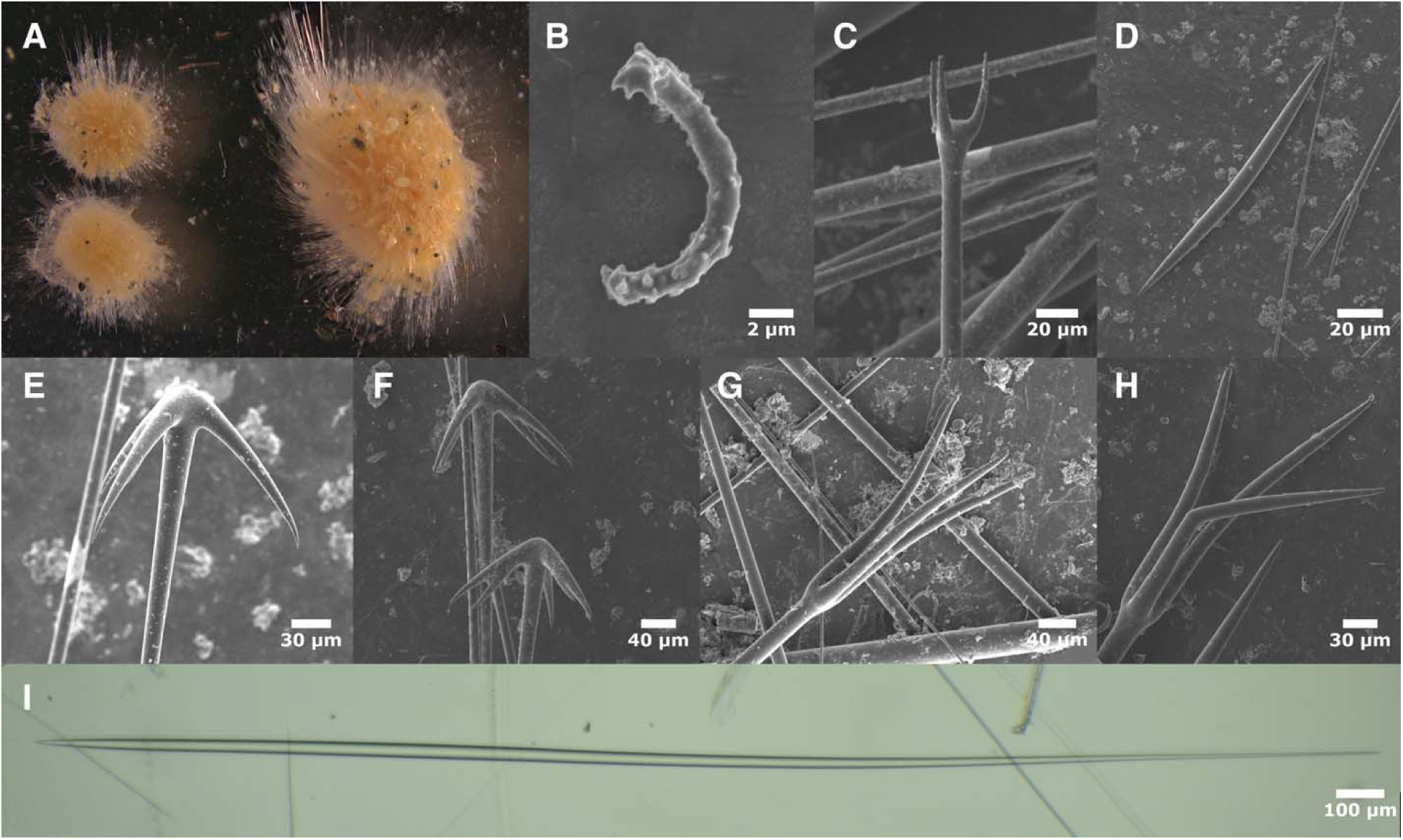
Craniella spinosa. A: Whole individuals (photo curtesy of Jean-Marc Gagnon); B: sigmaspire; C: short-cladded protriaene; D: oxea II; E: anatriaene; F: anatriaenes with forked clads; G: protriaene; H: protriaene with one bend clad; I: oxea I. All photos are of the holotype.

Oxeas I: the longest size-class of oxeas. Fusiform, thickest in the center and gradually tapering at both ends; strongly anisoform due to one filiform ending. 2861–4088–6952 x 29–36–43 μm (n=18). Type material previously described as 2730–4950 x 37 μm.

Oxeas II: stout cortical oxeas; one style was also seen. 428–837–1139 x 22–42–54 μm (n=26). Type material previously described as 726–1100 x 51–54 μm.

Oxeas III: long, thin, symmetrical oxeas; differentiated from oxeas I and II by being thinner relative to their length. Only one seen, 1795 x 13 μm. Not previously described from type material. Not shown in figure 19; see figures 14K & 16J for similar examples.

Thin-cladded anatriaenes: A large fraction (55%, 41/74) has at least one clad that is forked. Rhabds 3632–7686–9817 (n=3) x 11–20–24 μm (n=34). Clads 65–145–183 x 14–22–29 μm, with length:width ratios 4–7–10 (n=33). Clad:rhabd angle 30°–42°–49° (n=33). Type material previously described with rhabds 8220 x 16 μm, clads 131 x 13 μm; forked clads not noted.

Short-cladded anatriaenes: rare anatriaenes with very small clads and a large angle; possibly immature forms of other anatriaenes. Rhabd lengths not measured; widths 10–11–12 μm (n=3). Clads 31–42–48 x 8–9–11 μm, with length:width ratios 4–5–6 (n=3). Clad:rhabd angle 41°–50°– 57° (n=3). Not shown in figure 19.

Protriaenes/prodiaenes: as noted in the original description, clads are strongly curved: they are initially parallel with rhabd, then diverge mid-clad. Clads of equal lengths, and only modestly spread away from rhabd axis. Clad tips were angled in slightly, but this was only visible under SEM. Nearly all (97%, 33/34) were triaenes, but one diaene seen. About 10% were slightly deformed, with one clad bent at a sharp angle; angles were measured separately for these. Rhabds 3489–4335–5318 (n=6) x 13–18–23 μm (n=34). Longest clads 167–249–341 x 9–13–18 μm, with longest:shortest clad ratios 1.0–1.1–1.3 (n=34). Clad:rhabd angle 5°–12°–20° (n=29), or 27°–39°–48° (n=5) when one clad was sharply bent. Type material previously described with rhabds 5700 x 26 μm, clads 267 μm long.

Short-cladded protriaenes: with short, widely spread clads. Only one seen, rhabd 10 μm wide, longest clad 50 x 5 μm, longest:shortest clad ratio 1.2, Clad:rhabd angle 21°. Not previously described from type material.

Sigmaspires: C-shaped or spiraled, spiny throughout. Longest possible straight-line measurement across whole spicule 9–11–13 (n=15). Previously described as 13 μm long.

#### Distribution and habitat

Known from near Elk Bay and Comox, both in the Discovery Passage, Vancouver Island, British Columbia, depth 36–73 m.

#### Remarks

The unique shape of the protriaenes, noted in the original description, is diagnostic for this species. Branching anatriaene clads and bent protriaene clads were not noted in the original description, perhaps indicating they were an aberration present only in one sample. Additional traits, such as long protriaene clad lengths, and the apparent absence of large thick-cladded anatriaenes serve as additional traits to differentiate this species, as mentioned in the diagnostic section and table 3. I was unable to amplify any DNA from this species, so it is not included in the molecular phylogenies.

### *Craniella craniana* de Laubenfels, 1953

#### Material examined

Holotype: USNM 23233, 19.5 km North of Point Barrow, Beauford Sea, 226 m, 17-Aug-1949.

#### Diagnosis

The sigmaspires average 19 μm: smaller than *C. sigmoancoratum*, but larger than all other species in the Northern Hemisphere save *C. longipilis* (Topsent, 1904) and *C. polyura* Schmidt, 1870. *C. longipilis* is described from the Azores, and has much longer protriaene and anatriaene clads. The Arctic *C. polyura* has centrotylote sigmaspires lacking in *C. craniana*.

#### Morphology

I examined only a small, dried fragment of the holotype. The original description states that this species is subspherical to oval, 4-8 cm in largest dimension. The lower half is described as hairy while the upper half is covered in large, acute, cone-like projections up to 8 mm in length. No oscula were seen.

#### Skeleton

This was difficult to assess in the small dried fragment I examined. The species was previously described as having a radial skeleton, with spicular columns 1 mm wide radiating from the center of the sponge. No observations were reported regarding a cortex.

#### Spicules

Shown in figure 20, except when noted below.

**Figure 20.**
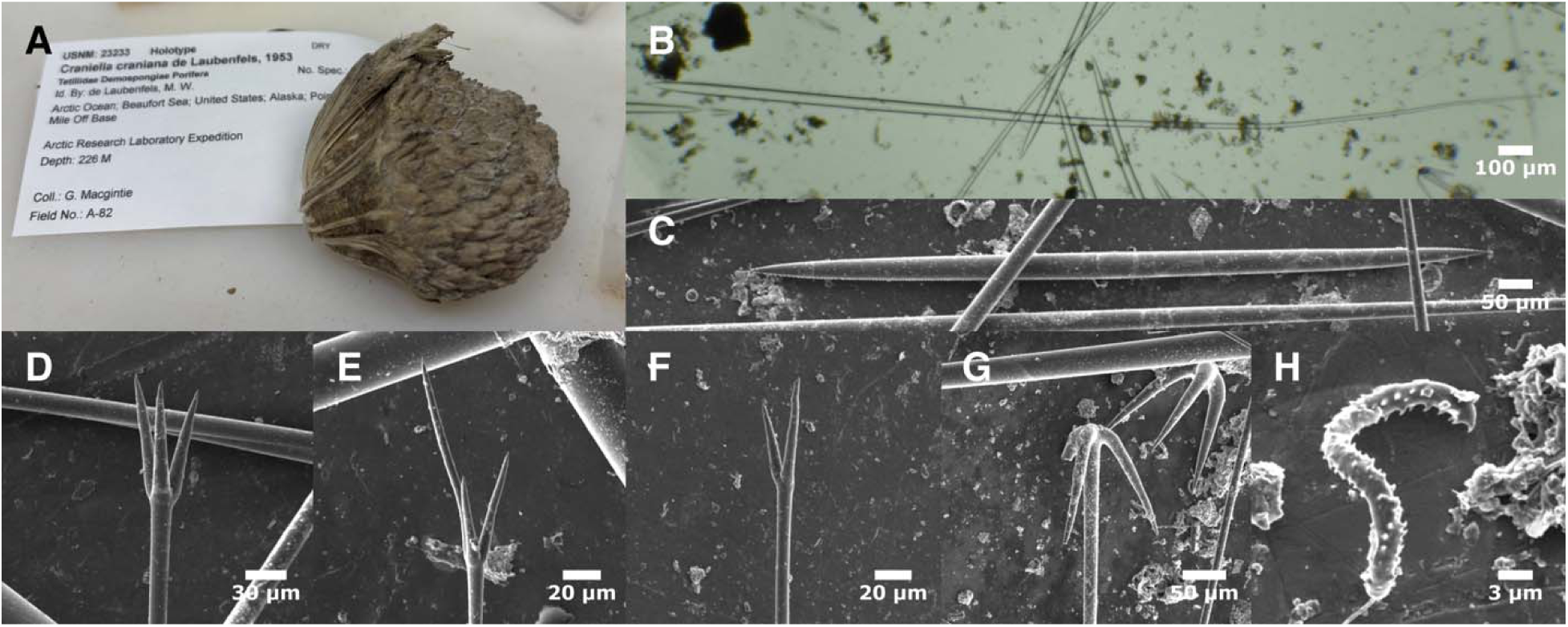
Craniella craniana. A: Dried holotype (photo curtesy of Abigail Reft); B: anisoform oxea Ib; C: oxea II; D-E: protriaene variation; F: prodiaene; G: anatriaenes; H: sigmaspire. All photos from holotype.

Oxeas Ia: the longest size-class of oxeas. Fusiform, thickest in the center and gradually tapering at both ends; variously ended, with one, two, or no filiform endings, but generally close to symmetrical. 2629–5070–6867 x 34–47–65 μm (n=17). Type material previously described as having oxeas several mm long and often over 100 μm thick. Not shown in figure 20.

Oxeas Ib: an additional class of large oxeas which were strikingly anisoform: one end with a blunt tip, one end filiform, and thickest near the blunt tip. 2232–3146–5350 x 25–39–89 μm (n=13). Not mentioned in original description.

Oxeas II: stout oxeas, presumably cortical. 669–1064–1855 x 19–35–49 μm (n=40). Not mentioned in original description.

Oxeas III: long, thin, symmetrical oxeas; differentiated from oxeas I and II by being thinner relative to their length. Only one seen, 2505 x 9 μm. Not mentioned in original description; not shown in figure 20.

Thin-cladded anatriaenes: Rhabds 4394–5130–5866 (n=2) x 9–16–21 μm (n=29). Clads 94–164– 245 x 13–19–28 μm, with length:width ratios 5–9–15 (n=28). Clad:rhabd angle 13°–29°–37° (n=33). Type material previously described as variable, with some clads 200 x 16 μm. Drawing shows a thin-cladded anatriaene (length:width ratio 8, clad:rhabd angle 30°).

Short-cladded anatriaenes: a few anatriaenes with smaller, thick clads were seen. One rhabd measured at 5853 μm long; widths 13–15–17 μm (n=4). Clads 50–66–78 x 17–21–23 μm, with length:width ratios 3–3–4 (n=4). Clad:rhabd angle 38°–40°–41° (n=4). Not mentioned in original description. Not shown in figure 20.

Protriaenes/prodiaenes/promonaenes: with fairly straight clads, varying from equal length to quite unequal, and only modestly spread away from rhabd axis. Only 65% (26/40) were triaenes, with many diaenes and one monaene seen. Rhabds 1115–2968–4821 (n=5) x 5–8–17 μm (n=45). Longest clads were 25–69–115 x 2–6–13 μm, with longest:shortest clad ratios of 1.0–1.5–2.5 (n=26). Clad:rhabd angle 8°–17°–29° (n=33). Type material previously described as having only prodiaenes, no measurements reported. The type drawing shows a prodiaene with equal-length clads and a small angle (16.8°)

Short-cladded protriaenes: with short, widely spread clads. Only one seen, rhabd 1768 x 3 μm, longest clad 10 x 2 μm, longest:shortest clad ratio 1.1, Clad:rhabd angle 30°. Not mentioned in original description. Not shown in figure 20.

Sigmaspires: C, S, or spiral-screw shaped, spiny throughout; larger end denticles absent. Longest possible straight-line measurement across whole spicule 14–19–25 (n=31) x 1.9–2.3–2.9 (n=9). Previously described as 17-22 μm long.

#### Distribution and habitat

Known only from the type location near Point Barrow, 226 m depth.

#### Remarks

This species is known from the Arctic, rather from the Northeast Pacific, and does not belong in this revision. The original description of this species contained little information, however, and I felt it was necessary to examine the type material in order to confirm that it was not present in the North Pacific as well. The spicule dimensions are provided here to aid future researchers. I also attempted DNA sequencing, but not even the smallest amplicons were successful.

The spicules of this species proved to have several differences compared to all Northeast Pacific species, including the sizes of the sigmaspires, the very small clads on the protriaenes, and the two classes of large oxeas.

### Dichotomous Key to the Tetillidae of the temperate Northeast Pacific

1A. Sponge globular/clavate rather than roughly spherical; no strongly radial skeleton; all oxeas are thin (maximum widths < 20 μm). Root of spicules anchor the sponge to soft sediment: Sediment-rooting *Tetilla*. …2

1B. Sponge spherical, strongly radial skeleton, some oxeas thicker than 20 μm: …3

2A. Sigmaspires present (may be rare); mean clad:rhabd angle of anatriaenes is < 40°: *Tetilla japonica*.

2B. Sigmaspires absent; mean clad:rhabd angle of anatriaenes is > 40°: *Tetilla mutabilis*.

3A. Pigmented ectosomal layer may be present, but no double-layer cortex with open spaces between layers. Longest oxeas sometimes filiform at only one end, but not strongly anisoform. Short, thin size class of consistently anisoform oxeas III present. Small (rhabd width < 5 um) size class of protriaenes abundant; large size class of protriaenes with clad lengths usually averaging < 100 um and with clads that are usually not widely spaced (angles < 20°) and of usually unequal (ratio ≥ 1.4) lengths: Sponge-rooting *Tetilla.* …4

3B. Double-layer cortex usually obvious in cross-section; no small class of anisoform oxeas III, but largest size class of oxeas are usually strongly anisoform; lacking small (width < 5 um) protriaenes; large protriaenes have long (> 100 um) clads. Not known South of British Columbia. Genus *Craniella*. …7

4A. Thin-cladded anatriaenes (length:width ratio > 7) present; bearing a field of small oscula that may not be visible post-preservation; thick-cladded anatriaenes with long (> 60 um) clad lengths: …5

4B. Thin-cladded anatriaenes (length:width ratio > 7) absent; sponge bearing 1–10 discrete oscula, still visible after preservation; thick-cladded anatriaenes with short (< 60 um) chord lengths: …6

5A. Large size-class of protriaenes are primarily (> 80%) triaenes, with monaenes absent; these protriaenes have clads of unequal lengths (ratio ≥ 1.8). Live sponges display an oscular area with many small oscula, 0.3-1.5 mm in size, not covered by a pore sieve: *Tetilla vancouverensis* sp. nov.

5B. Large size-class of protriaenes are a mix of tri-, di- and monaenes, with triaenes in the minority; these spicules have clads that are equal or modestly unequal in length (ratios 1.1–1.7).

Oscular pores very small and covered by a pore sieve, often concealed in preserved sponges; oscular area usually surrounded by prominent spicule fringe: *Tetilla villosa*.

6A. Anisoform oxeas III are thinner than oxeas II; large size-class of protriaenes are a mix of tri- and diaenes with rhabd:clad angles < 17°. Common in California and known from both intertidal and subtitidal zones; colors gray, black, and/or yellow alive: *Tetilla arb*.

6B. Anisoform oxeas III average thicker than oxeas II; large size-class of protriaenes are exclusively triaenes with rhabd:clad angles >17°. Rare. Only known sample was white alive with a single oscule: *Tetilla losangelensis* sp. nov.

7A. Sigmaspires absent. Oxeas II present in two size classes; smaller size class are centrotylote: *Craniella sputnika.*

7B. Abundant sigmaspires and no centrotylote oxeas: …8

8A. Harpoon-shaped anamonanenes abundant: Craniella hamatum.

8B. Harpoon-shaped anamonanenes absent:

…99A. Sigmaspires average > 21 μm; they are C-shaped with unspined shafts and large end denticles: *Craniella sigmoancoratum*.

9B. Sigmaspires average < 21 μm: …10

10A. Protriaene clads average > 200 μm: …11

10B. Protriaene clads average < 200 μm: …12

11A. Protriaenes clads are nearly parallel with the rhabd at first, then diverge mid-clad; sigmaspires 9–13 μm; only one size class of oxeas II: *Craniella spinosa*.

11B. Protriaene clads straight; sigmaspires 14–18 μm; oxeas II in two size classes; sponge covered in long spines: *Craniella uniiguni* sp. nov.

13A. Protriaene clads average > 130 μm; anatriaene clads average > 100 μm; sigmaspires evenly covered in fine spines (only visible with SEM): *Craniella vermisigma* sp. nov.

12A. Sigmaspires average > 13 μm: …13

12B. Sigmaspires average < 13 μm: …14

13B. Protriaene clads average < 130 μm; anatriaene clads average < 100 μm; sigmaspires with scattered large spines (only visible with SEM): *Craniella shemya* sp. nov.

14A. Thick-cladded anatriaenes have a unique “rocket ship” shape with low rhabd:clad angles (< 30°): *Craniella rocheta* sp. nov.

14B. Thick-cladded anatriaenes have a typical shape with rhabd:clad angles > 30° …15 15A. Sigmaspires average < 9 μm; protriaene clads average < 136 μm long: *Craniella amlia* sp. nov.

15B. Sigmaspires average > 9 μm; protriaene clads average > 136 μm long: *Craniella columbiana* sp. nov.

## Conclusions

This study presents the first comprehensive systematic analysis of the family Tetillidae in the Northeast Pacific. It resolves long-standing taxonomic puzzles—such as the identity of the intertidal tetillid in California—while clarifying the status of previously described species and uncovering many new ones. Despite the large number of species, I found that morphological taxonomy was sufficient to distinguish all of them, in contrast to groups with simpler spicules (Xavier *et al*., 2010; Turner *et al*., 2025). This was made possible only by making many measurements and including new traits like anatriaene angles, quantification of the inequality of protriaene lengths, and the proportion of prodiaenes among them. I suggest that morphological studies in this family include such measurements in the future. I have also included all raw measurement data, as well as explicitly stating how measurements were made. This allows future statistical comparisons among species, and should become the norm. Despite success at the morphological level, DNA data proved essential for confirming species boundaries, as some morphological characters varied within species in unexpected ways. It also would have been difficult to confirm such a large number of species of *Craniella* from so few samples without DNA support for nearly all of them.

In contrast to species diagnosis, genus-level morphological taxonomy of the Tetillidae appears to be on shaky ground. Generic definitions in Tetillidae rely heavily on ectosomal morphology and the arrangement of pores and oscula on the sponge surface. My results suggest some of these features (oscular morphology) evolve rapidly, while others (having a pseudocortex) are prone to convergent evolution. Addressing this challenge will likely require incorporating DNA characters into the taxonomic framework, as has been proposed for other sponge groups (Cárdenas *et al*., 2011; Turner *et al*., 2024, 2025; van der Sprong *et al*., 2025).

Additional, striking findings include the discovery of eight *Craniella* species from just 11 vouchers collected in the Aleutian Islands, suggesting this archipelago may be a hotspot of *Craniella* diversity. Further sampling will almost certainly reveal additional species. I also document what appears to be the first species introduction in the family: *T. japonica* in California. This highlights yet another reason to integrate DNA data into systematic studies of Tetillidae, as this may uncover further introductions.

Overall, this work transforms our understanding of Northeast Pacific tetillids. What was once a poorly defined and confusing assemblage is now a set of clearly diagnosed species, with characters that allow them to be distinguished in the field, in the lab, and/or with DNA data. Still, this represents only a first step. The biology of these sponges remains largely unexplored, and I hope this study provides a solid foundation for future research on these remarkable animals.

## Acknowledgements

I am grateful for the help and support of many people in UCSB’s Marine Science Institute, especially Robert Miller, Clint Nelson, Christoph Pierre, and Christian Orsini. Steve Lonhart and The Monterey Bay National Marine Sanctuary, was instrumental in facilitating collections in Central California. The Olympic Coast National Marine Sanctuary was instrumental in facilitating collections in Washington. The Natural History Museum of Los Angeles’ DISCO program facilitated collections in Los Angeles County. I am grateful that the Alaska Fisheries Science Center (NOAA) collected the Alaskan sponges and deposited them in museum collections for future use. Allen Collins, Hugh MacIntosh, Christina Piotrowski, Kathy Omura, and Jean-Marc Gagnon graciously provided access to their collections at the Smithsonian National Museum of Natural History, the Royal British Columbia Museum, the California Academy of Sciences, the Natural History Museum of Los Angeles, and the Canadian Museum of Nature, respectively; Jeff Goddard and Brandon Stidum also provided crucial samples. Several of the California Academy of Sciences samples were collected by the Coral Reef Research Foundation under contract to the US National Cancer Institute. I also thank the Makah Tribal Nation for supporting the study of sponges from Tatoosh Island, Washington.

## Data Availability Statement

The data underlying this article are available in the article and in its online supporting information.

## Supporting Information

Table S1. Metadata for all samples investigated.

Table S2. Measurement data from all spicules; “n/a” indicates a measurement was not taken for that spicule, or that the measure was not relevant for that spicule type.

Table S3. Sanger sequencing primers used.

Figure S1. Illustrated guide to measuring protriaene spicule widths.

Figure S2. Illustrated guide to measuring protriaene spicule lengths.

Figure S3. Illustrated guide to measuring protriaene spicule angles.

Figure S4. Illustrated guide to measuring anatriaene lengths and widths.

Figure S5. Illustrated guide to measuring anatriaene spicule angles.

Figure S6. Illustrated guide to measuring normal anamonaene spicules.

Figure S7. Illustrated guide to measuring boathook anamonaene spicules.

Figure S8. Maximum likelihood phylogeny of the 28S locus. New sequences are shown in bold, along with collection location and voucher number. Genbank data includes accession numbers and collection locations (when known). Node confidence is indicated with bootstrap values. Scale bar indicates substitutions per site.

Figure S9. Maximum likelihood phylogeny of the cox1 locus. New sequences are shown in bold, along with collection location and voucher number. Genbank data includes accession numbers and collection locations (when known). Node confidence is indicated with bootstrap values. Scale bar indicates substitutions per site.

Figure S10. Sanger sequencing success vs. sample age. The largest attempted amplicon (700 bp or larger) was successful on most recently collected material, but failed on most samples over 20 years old. A small random number (jitter) has been added to the dates in order to see points that would otherwise overlap. This result should be considered hypothesis-forming rather than hypothesis-testing, as these points are non-independent due to partial correlations between age, collection location, collector, museum, and taxonomic subgroup.

Figure S11. Oxeas in the holotype of *T. arb.* When plotting the length and width of oxeas (in microns) in this and other sponge-rooting *Tetilla*, oxeas I (black) and oxeas II (blue) are clearly distinct. Oxeas III (red) are harder to distinguish by size, and are identified by their anisoform nature. It is possible that oxeas III are oxeas I that are still forming.

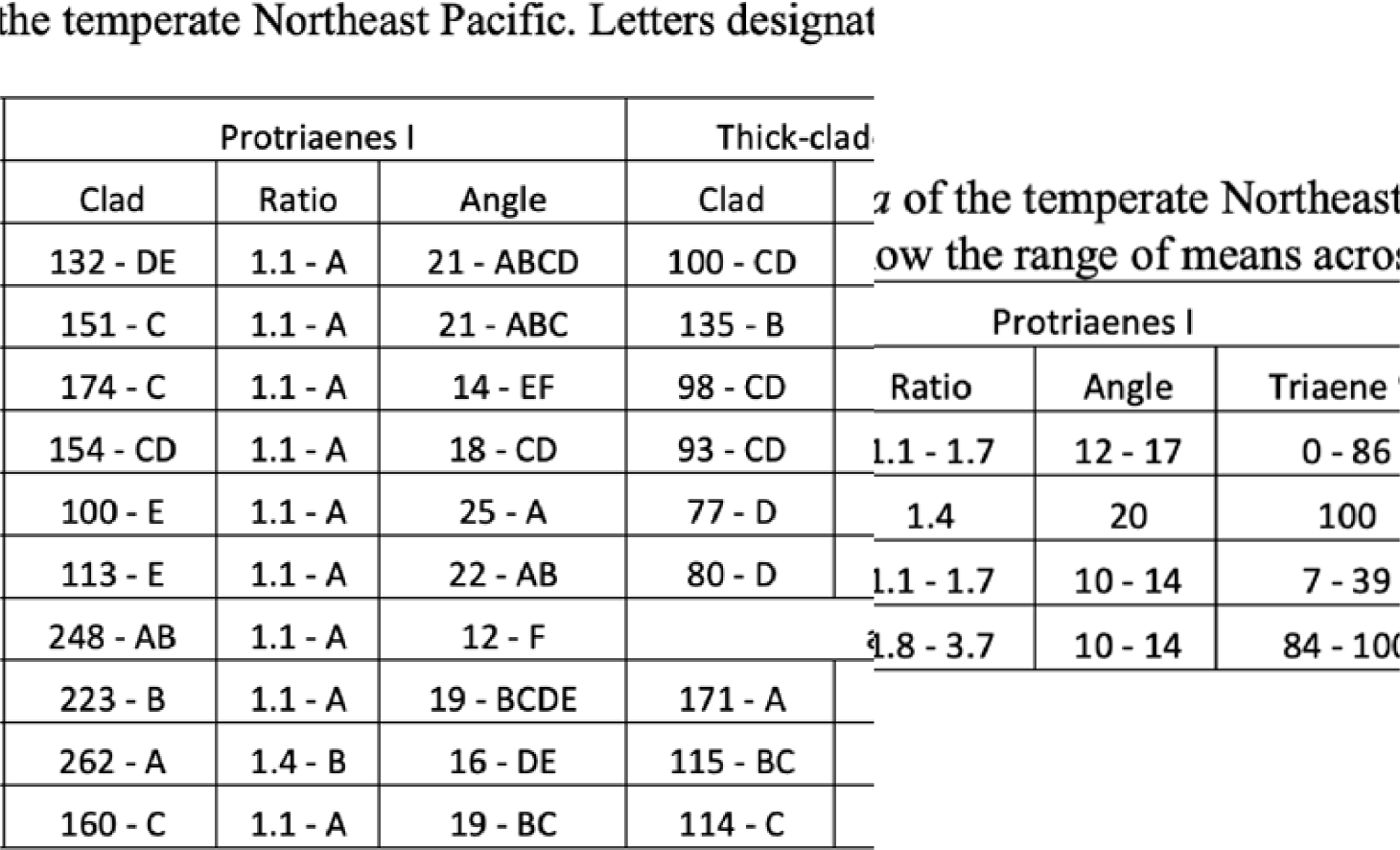

