## Supplemental figures for "Systematic revision of the family Tetillidae (Porifera: Demospongiae) in the temperate Northeast Pacific"

**Supplementary Figures**

Figure S1. Illustrated guide to measuring protriaene spicule widths

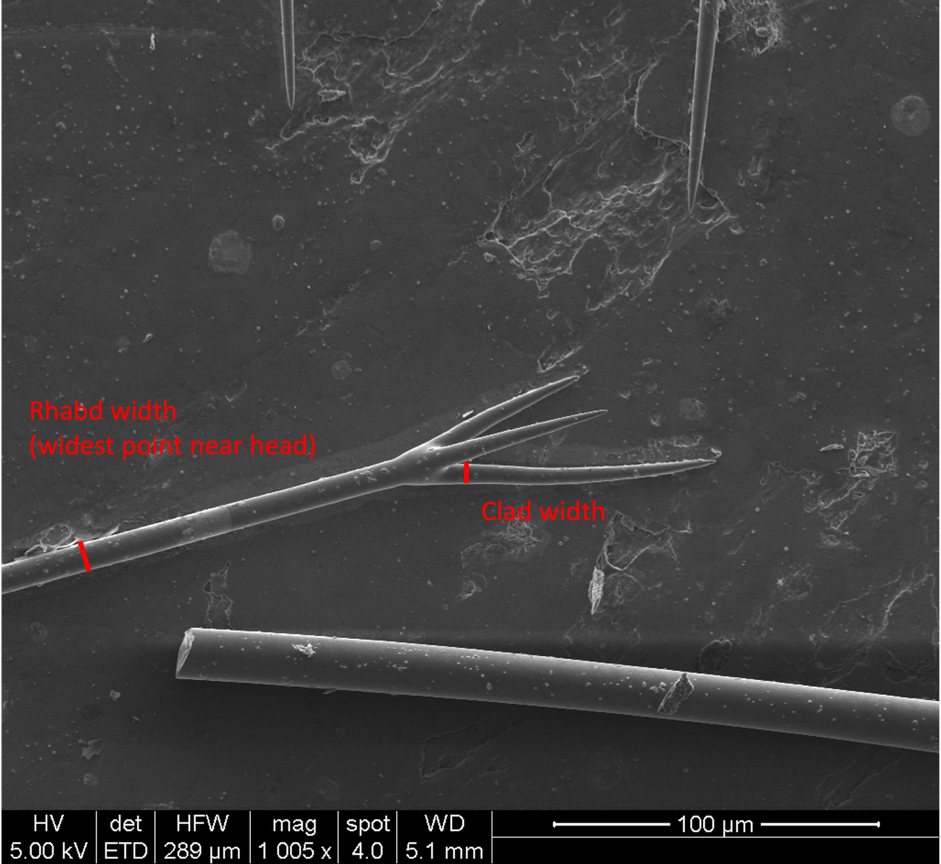

Figure S2. Illustrated guide to measuring protriaene spicule lengths.

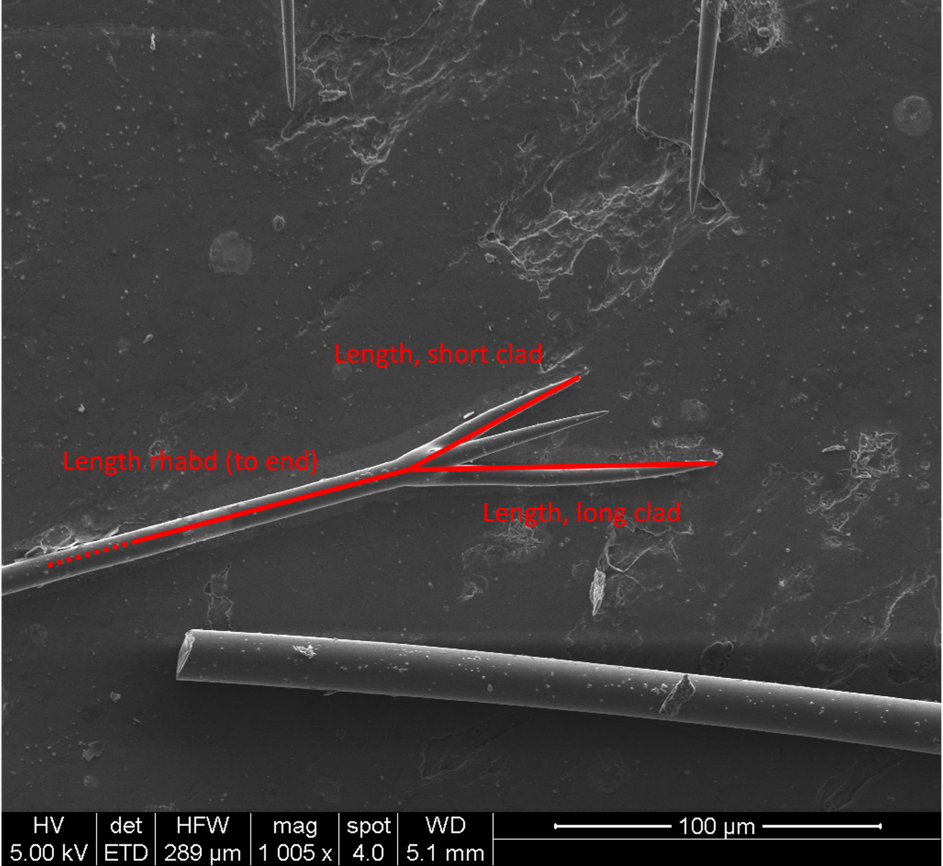

Figure S3. Illustrated guide to measuring protriaene spicule angles.

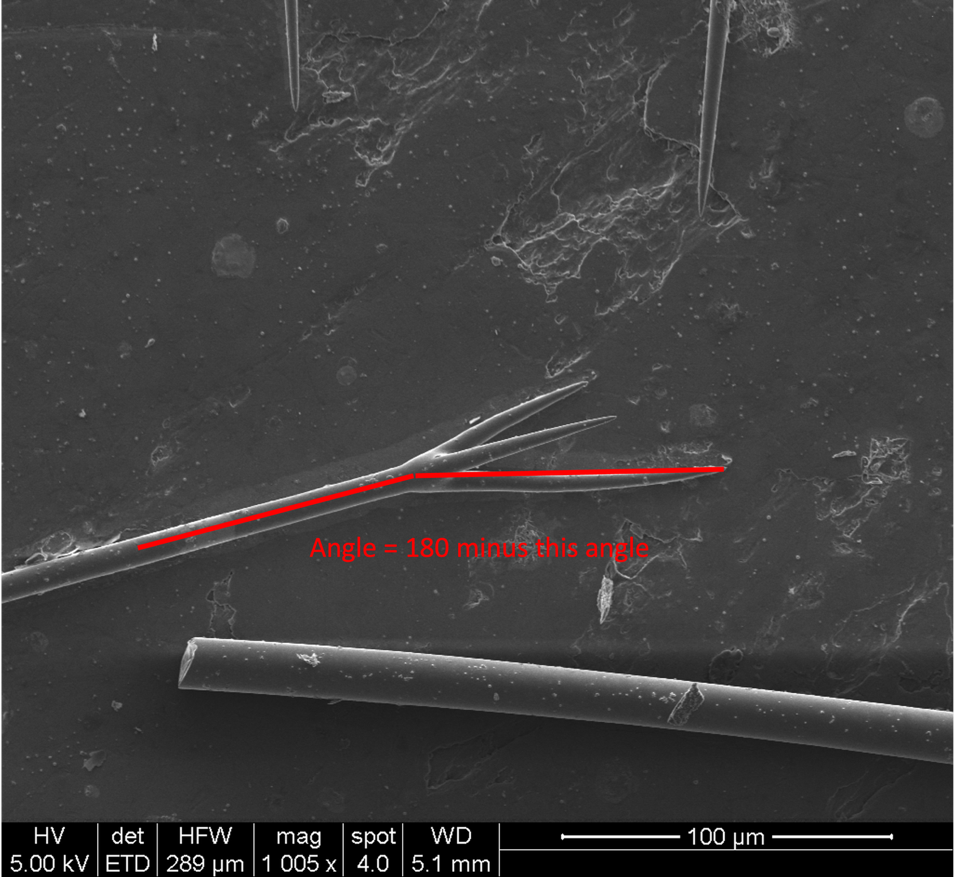

Figure S4. Illustrated guide to measuring anatriaene lengths and widths.

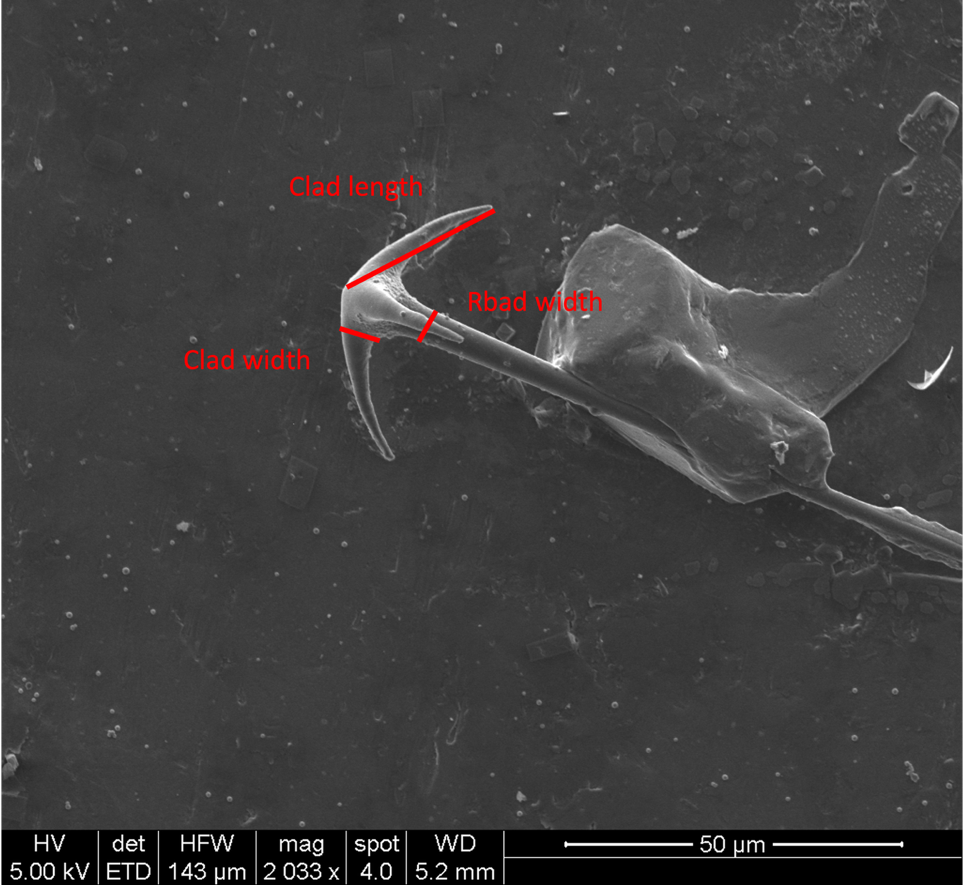

Figure S5. Illustrated guide to measuring anatriaene spicule angles.

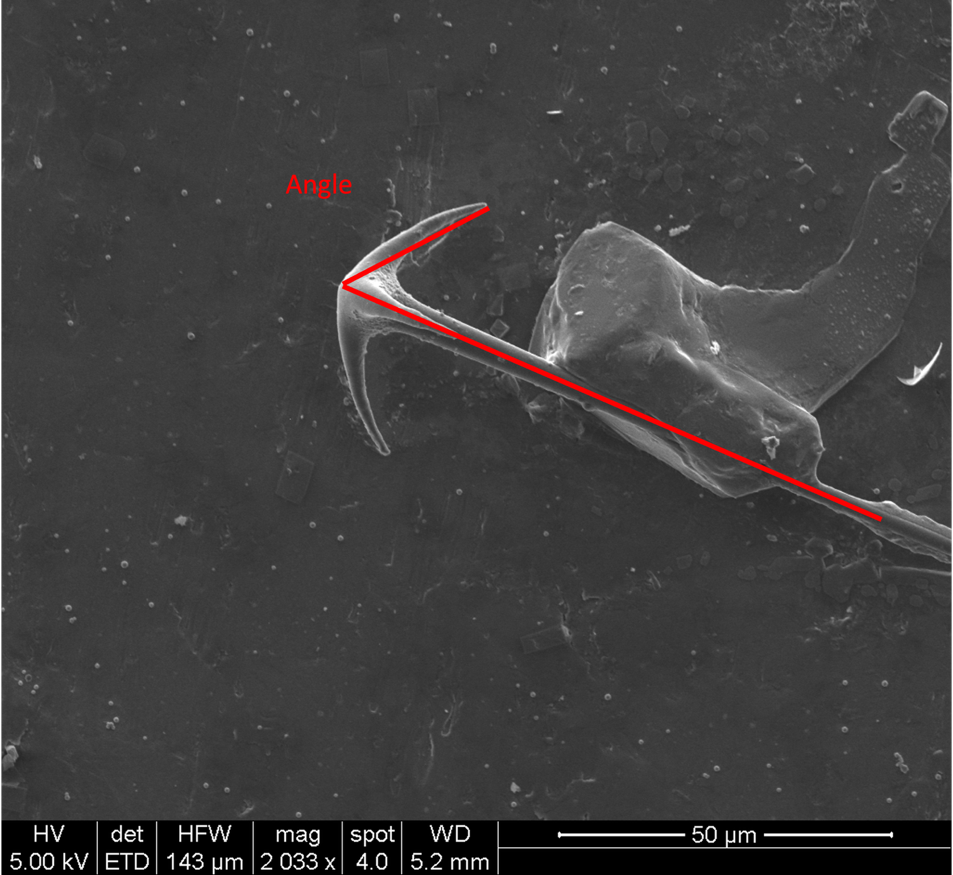

Figure S6. Illustrated guide to measuring normal anamonaene spicules.

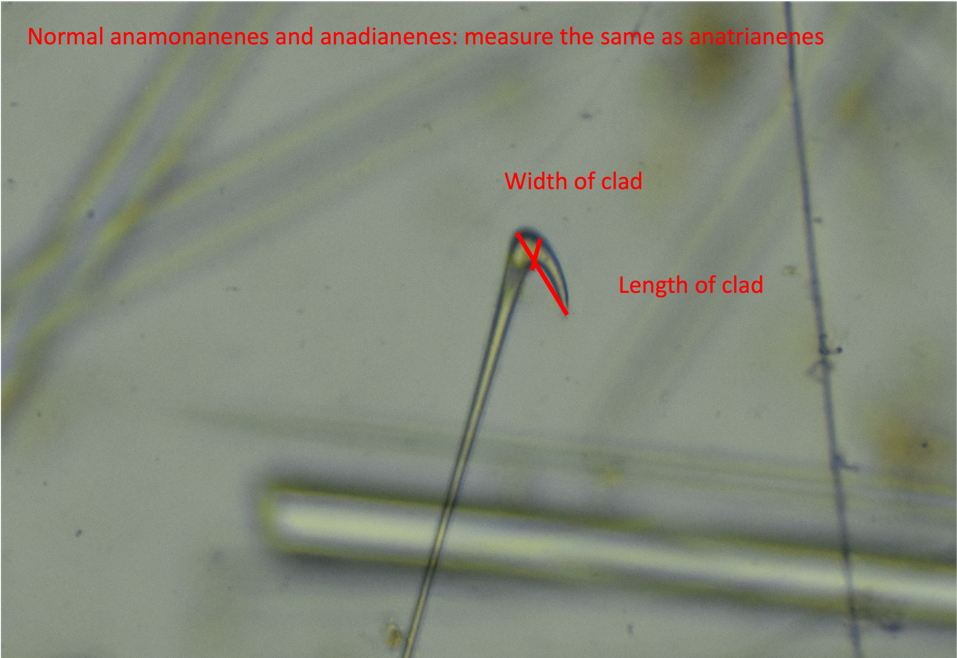

Figure S7. Illustrated guide to measuring boathook anamonaene spicules.

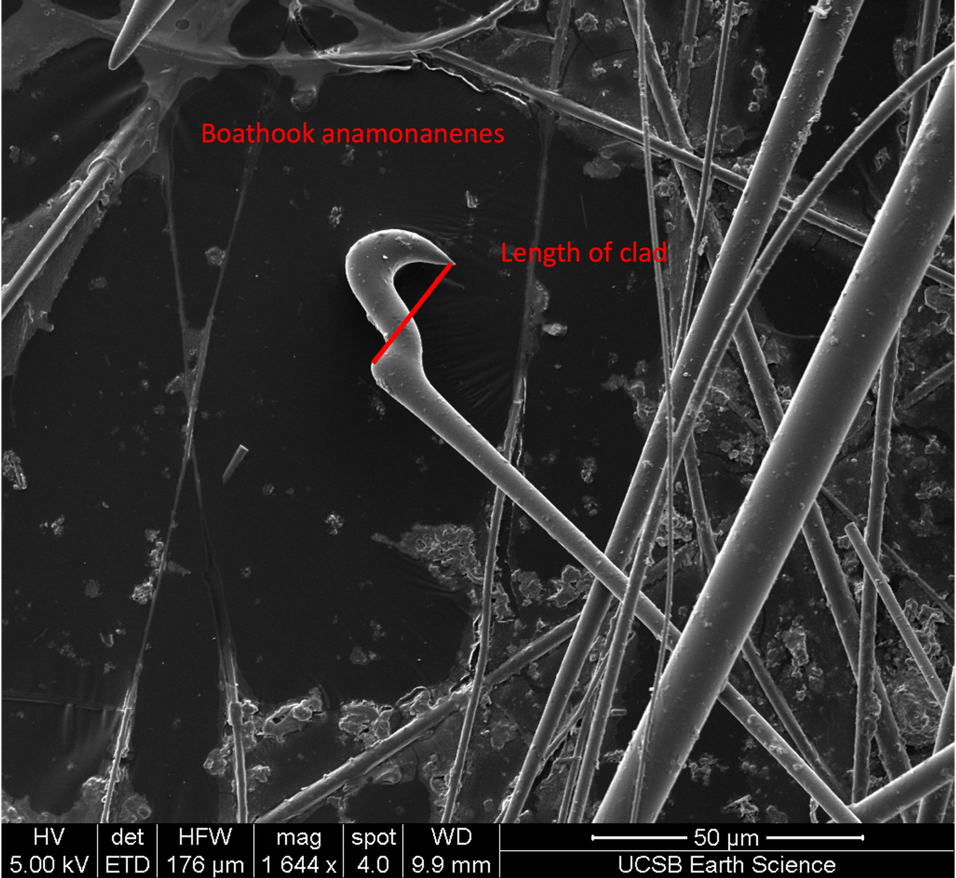

Figure S8. Maximum likelihood phylogeny of the 28S locus. New sequences are shown in bold, along with collection location and voucher number. Genbank data includes accession numbers and collection locations (when known). Node confidence is indicated with bootstrap values. Scale bar indicates substitutions per site.

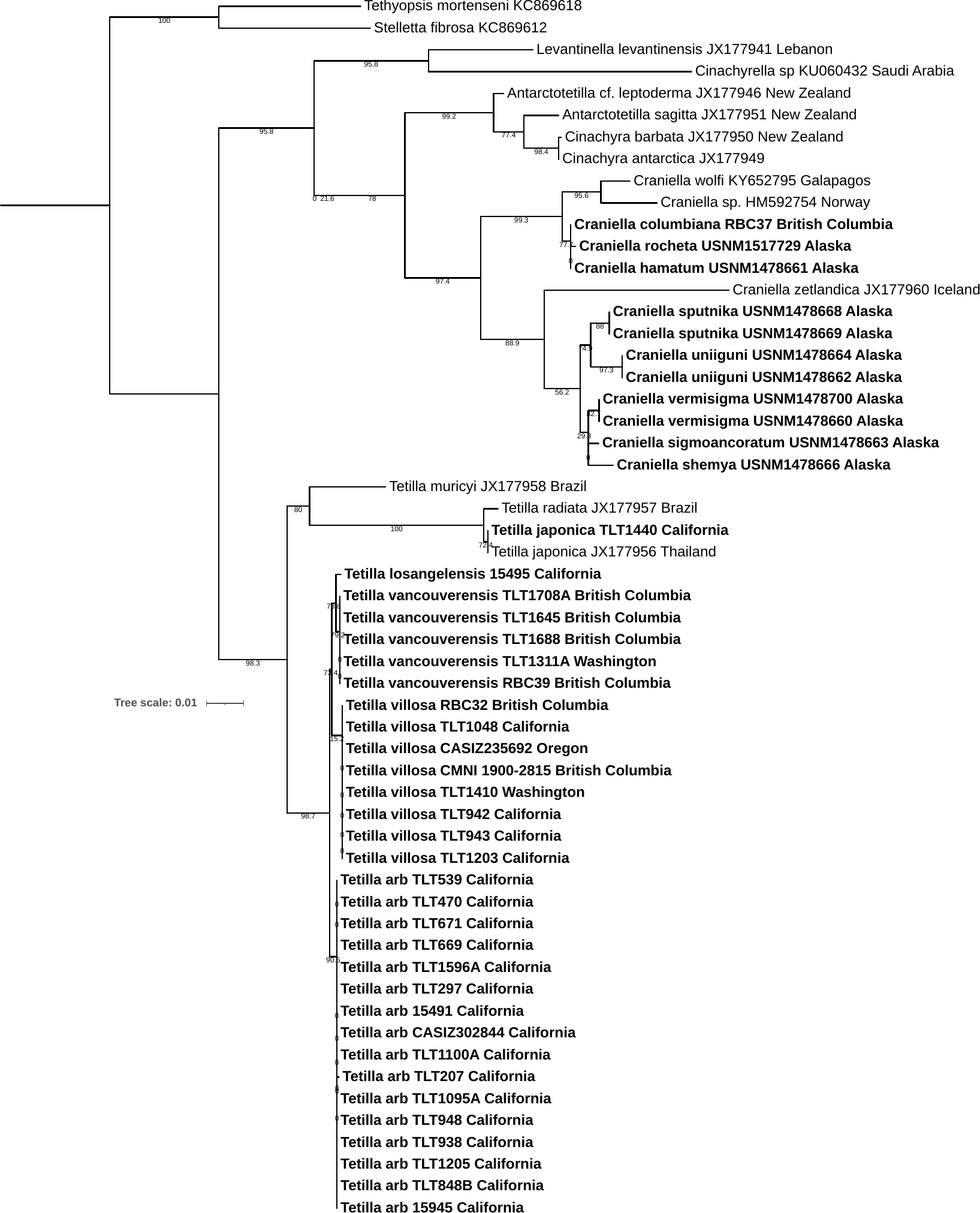

Figure S9. Maximum likelihood phylogeny of the cox1 locus. New sequences are shown in bold, along with collection location and voucher number. Genbank data includes accession numbers and collection locations (when known). Node confidence is indicated with bootstrap values. Scale bar indicates substitutions per site.

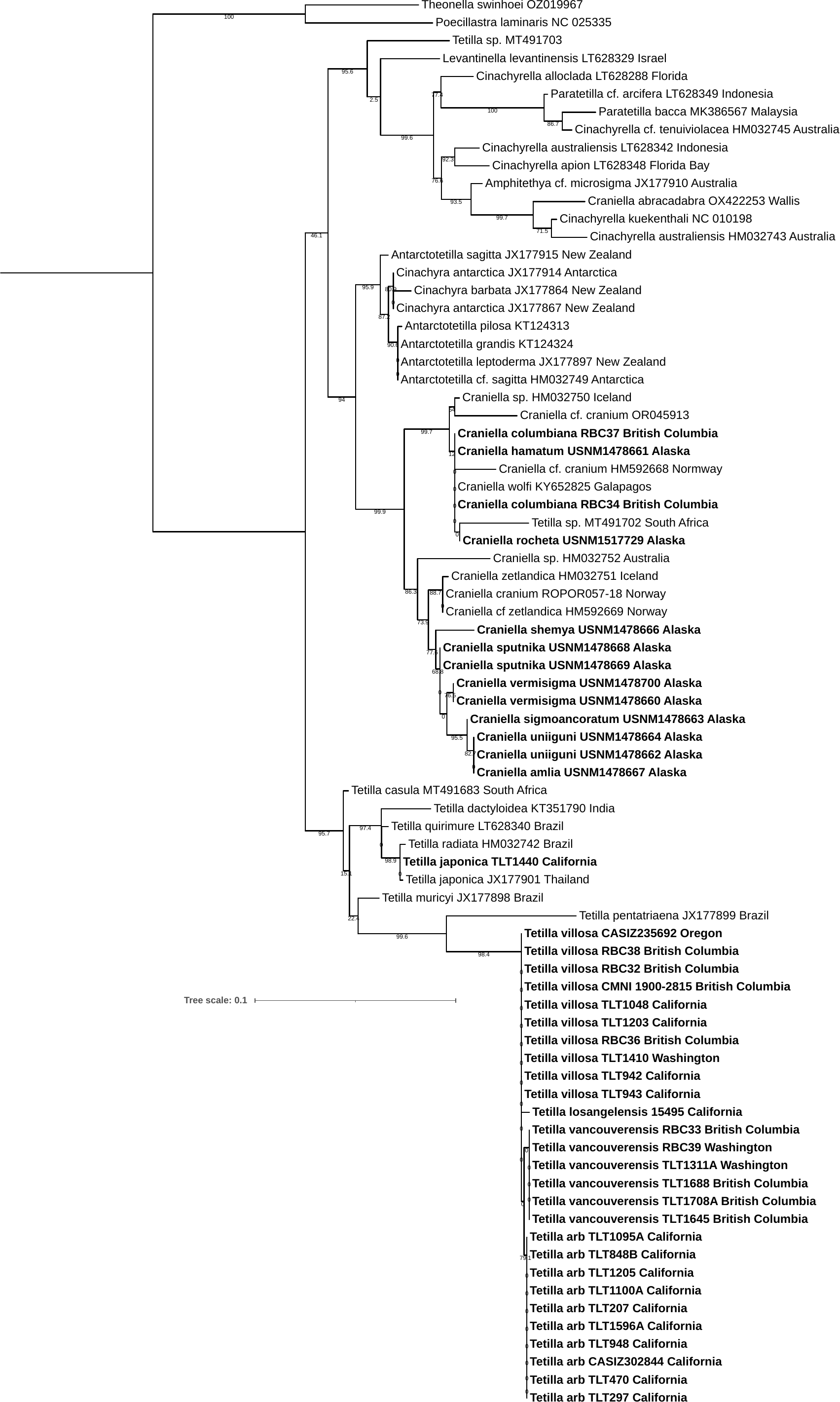

Figure S10. Sanger sequencing success vs. sample age. The largest attempted amplicon ( 700 bp or larger) was successful on most recently collected material, but failed on most samples over 20 years old. A small random number (jitter) has been added to the dates in order to see points that would otherwise overlap. This result should be considered hypothesis-forming rather than hypothesis-testing, as these points are non-independent due to partial correlations between age, collection location, collector, museum, and taxonomic subgroup.

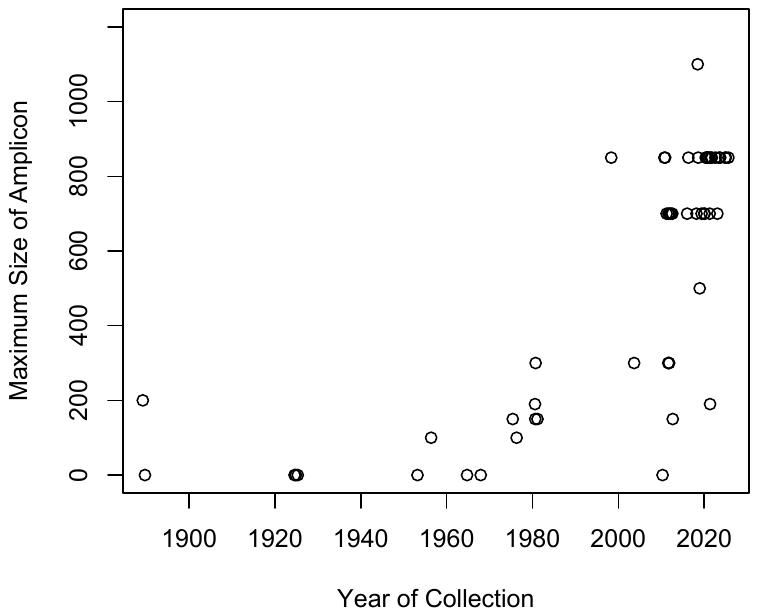

Figure S11. Oxeas in the holotype of *T. arb.* When plotting the length and width of oxeas (in microns) in this and other sponge-rooting *Tetilla*, oxeas I (black) and oxeas II (blue) are clearly distinct. Oxeas III (red) are harder to distinguish by size, and are identified by their anisoform nature. It is possible that oxeas III are oxeas I that are still forming.

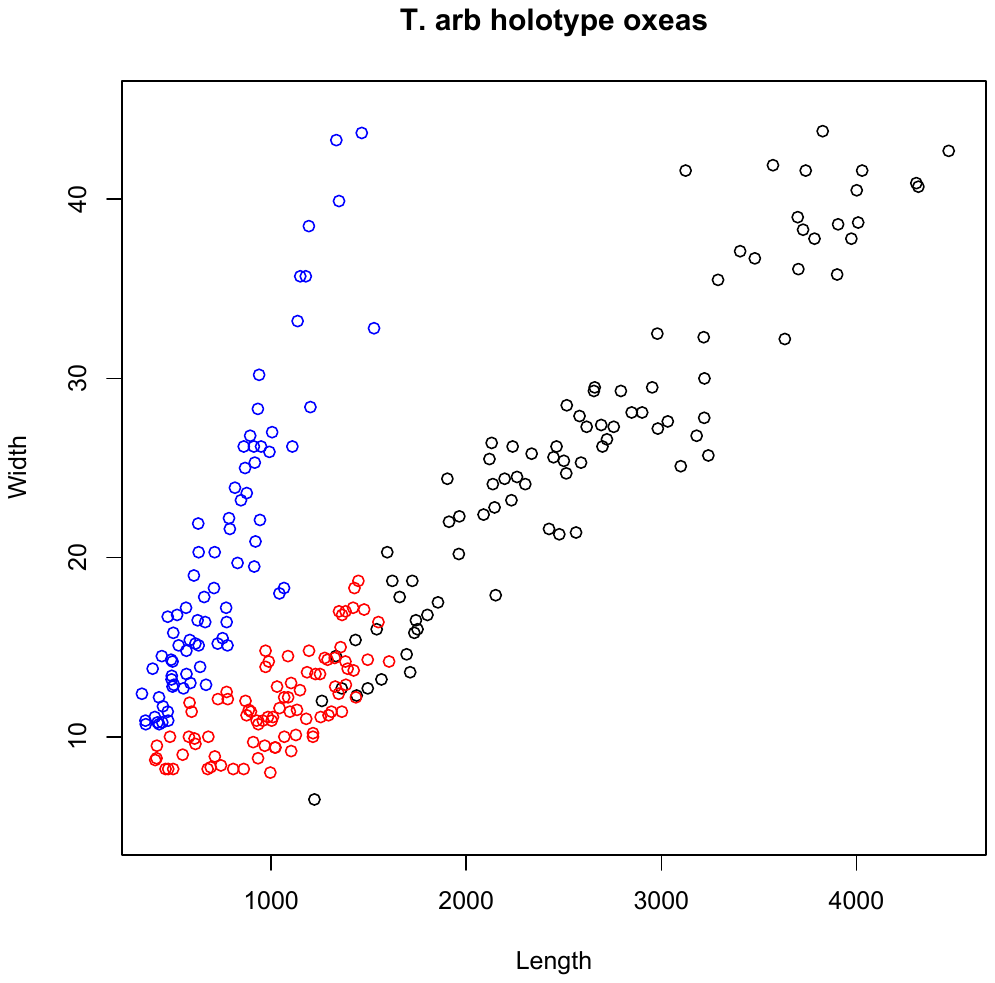
