## Supplemental table 3 for "Systematic revision of the family Tetillidae (Porifera: Demospongiae) in the temperate Northeast Pacific"

Table S3. Sequencing primers used.

| Locus | Name | Sequence | Notes |
| --- | --- | --- | --- |
| 28S D3-D5 | Por28S-830F | CATCCGACCCGTCTTGAA | Porifera ‘universal’, ~700 bp |
| 28S D3-D5 | Por28S-1520R | GCTAGTTGATTCGGCAGGTG | Porifera ‘universal’; pair with above |
| 28S D1-D2 | Tetra-15F | GCGAGRYGACCCGCYGAAC | Worked in *Tetilla,* ~700 bp |
| 28S D1-D2 | Por28S-878R | CACTCCTTGGTCCGTGTTTC | Porifera ‘universal’; pair with above |
| 28S C2-D2 | Tetra-C2 | GAAAAGYACTTTGGAAAGAGAGT | Worked in *Tetilla,* ~500 bp within D1-D2 |
| 28S C2-D2 | D2 | TCCGTGTTTCAAGACGGG | Porifera ‘universal’; pair with above |
| 28S-150bp | Tetilla_28S_150L | TTCTGTCCGTTGAGCCTCC | Worked in *Tetilla*, ~150 bp within D1-D2; Pair with D2 |
| 28S-200bp | Tetilla_28S_200L | TGTACGGGCACTCTCACG | Worked in *Tetilla*, ~200 bp within D1-D2 |
| 28S-200bp | Tetilla_28S_200R | CTGCGCTACCTACCCGAC | Pair with above |
| 28S-300bp | Cran-cox300L | GGYGCCCGCATATTAAAGATAGT | Worked in *Craniella,* ~300 bp within D1-D2 |
| 28S-300bp | Cran-cox300R | SGGKTTCGGTAATTGAATGGT | Pair with above |
| 28S_Cramini | Tetra-Dmini | ACTCCTTGGTCCGTGTTTCA | Worked in *Craniella,* ~200 bp within D1-D2 |
| 28S_Cramini | Cran-Cmini | GAACTTACAGCCGWCRGTCY | Pair with above |
| cox1_Folmer | LCO1490 | GGTCAACAAATCATAAAGAYATYGG | Porifera ‘universal’, ~650 bp; only worked in *T. japonica* |
| cox1_Folmer | HCO2198 | TAAACTTCAGGGTGACCAAARAAYCA | Pair with above |
| cox1_Folmer+ | Tetilla_cox1L | CCTTTATATCGGGGCGCCA | Worked in *Tetilla,* ~850 bp |
| cox1_Folmer+ | Tetilla_cox1R | ACTACTTGCTACCACGATTCC | Pair with above |
| cox1_nanoA | Tetilla_nanoAL | GATAATCGCCGTACCTACCG | Worked in *Tetilla* and *Craniella*, pair with Tetilla_cox1R, ~150bp |
| cox1_nanoC | Tetilla_nanoCR | AGATAGCCGCATCGACTGAA | Worked in *Tetilla*, pair with Tetilla_cox1L, ~190bp |
| cox1_300 | Cran-cox300L | GGYGCCCGCATATTAAAGATAGT | Worked in *Craniella,* ~300 bp within Folmer region |
| cox1_300 | Cran-cox300R | SGGKTTCGGTAATTGAATGGT | Pair with above |
